# DCIS genomic signatures define biology and clinical outcome: Human Tumor Atlas Network (HTAN) analysis of TBCRC 038 and RAHBT cohorts

**DOI:** 10.1101/2021.06.16.448585

**Authors:** Siri H Strand, Belén Rivero-Gutiérrez, Kathleen E Houlahan, Jose A Seoane, Lorraine M King, Tyler Risom, Lunden A Simpson, Sujay Vennam, Aziz Khan, Luis Cisneros, Timothy Hardman, Bryan Harmon, Fergus Couch, Kristalyn Gallagher, Mark Kilgore, Shi Wei, Angela DeMichele, Tari King, Priscilla F McAuliffe, Julie Nangia, Joanna Lee, Jennifer Tseng, Anna Maria Storniolo, Alastair M Thompson, Gaorav P Gupta, Robyn Burns, Deborah J Veis, Katherine DeSchryver, Chunfang Zhu, Magdalena Matusiak, Jason Wang, Shirley X Zhu, Jen Tappenden, Daisy Yi Ding, Dadong Zhang, Jingqin Luo, Shu Jiang, Sushama Varma, Lauren Anderson, Cody Straub, Sucheta Srivastava, Christina Curtis, Rob Tibshirani, Robert Michael Angelo, Allison Hall, Kouros Owzar, Kornelia Polyak, Carlo Maley, Jeffrey R Marks, Graham A Colditz, E Shelley Hwang, Robert B West

**Affiliations:** Department of Pathology, Stanford University School of Medicine, Stanford, CA 94305, USA; Department of Molecular Medicine, Aarhus University Hospital, 8200 Aarhus N, Denmark; Stanford Cancer Institute, Stanford University School of Medicine, Stanford, CA 94305, USA; Vall d’Hebron Institute of Oncology, Barcelona, 08035, Spain; Department of Surgery, Duke University School of Medicine, Durham, NC 27708, USA; School of Life Sciences, Arizona State University, Tempe, AZ 85281, USA; Department of Pathology, Montefiore Medical Center, Bronx, NY 10467, USA; TBCRC Loco-Regional Working Group; Department of Pathology, Mayo Clinic, Rochester, MN 55902, USA; Department of Surgery, University of North Carolina at Chapel Hill, Chapel Hill, NC 27599, USA; Department of Pathology, University of Washington, Seattle, WA 98195, USA; Department of Pathology, University of Alabama at Birmingham, Birmingham, AL 35294, USA; Department of Medicine, University of Pennsylvania, Philadelphia, PA 19104, USA; Breast Oncology Program, Dana-Farber Cancer Institute, Boston, MA 02215, USA; Department of Surgery, Brigham and Women’s Hospital, Boston, MA 02115, USA; Department of Surgery, University of Pittsburgh, Pittsburgh, PA 15213, USA; Dan L. Duncan Comprehensive Cancer Center, Baylor College of Medicine, Houston TX 77030, USA; Department of Surgery, MD Anderson Cancer Center, Houston, TX 77030, USA; Department of Surgery, University of Chicago, Chicago, IL 60637, USA; Department of Medicine, Indiana University, Indianapolis, IN 46202, USA; Department of Surgery, Baylor College of Medicine, Houston, TX 77030, USA; Department of Radiation Oncology, University of North Carolina at Chapel Hill, Chapel Hill, NC 27599, USA; TBCRC, The EMMES Corporation, Rockville, MD 20850, USA; Department of Medicine, Washington University School of Medicine, St. Louis, MO 63108, USA; Departments of Pathology & Immunology, Washington University School of Medicine, St. Louis, MO 63108, USA; Department of Surgery, Washington University School of Medicine, St. Louis, MO 63110, USA; Department of Biomedical Data Science, Stanford University, Stanford, CA 94305, USA; Duke Cancer Institute, Duke University School of Medicine, Durham, NC 27708, USA; Department of Medicine and Genetics, Stanford University, Stanford, CA 94305, USA; Department of Statistics, Stanford University, Stanford, CA 94305, USA; Department of Pathology, Duke University School of Medicine, Durham, NC 27708, USA; Department of Biostatistics & Bioinformatics, Duke University School of Medicine, Durham, NC 27708, USA; Department of Medical Oncology, Dana-Farber Cancer Institute, Boston, MA 02215, USA

**Keywords:** Ductal carcinoma in situ, RNA gene expression profiling, whole genome sequencing, multiplex immunohistochemistry, invasive breast cancer, precancer, outcome, human tumor atlas network, breast, tumor microenvironment, classifier, prognosis, recurrence, progression

## Abstract

Ductal carcinoma *in situ* (DCIS) is the most common precursor of invasive breast cancer (IBC), with variable propensity for progression. We have performed the first multiscale, integrated profiling of DCIS with clinical outcomes by analyzing 677 DCIS samples from 481 patients with 7.1 years median follow-up from the Translational Breast Cancer Research Consortium (TBCRC) 038 study and the Resource of Archival Breast Tissue (RAHBT) cohorts. We identified 812 genes associated with ipsilateral recurrence within 5 years from treatment and developed a classifier that was predictive of DCIS or IBC recurrence in both cohorts. Pathways associated with recurrence include proliferation, immune response, and metabolism. Distinct stromal expression patterns and immune cell compositions were identified. Our multiscale approach employed *in situ* methods to generate a spatially resolved atlas of breast precancers, where complementary modalities can be directly compared and correlated with conventional pathology findings, disease states, and clinical outcome.

**HIGHLIGHTS:**

⍰ Development of a new classifier for DCIS recurrence or progression
⍰ Outcome associated pathways identified across multiple data types and compartments
⍰ Four stroma-specific signatures identified
⍰ CNAs characterize DCIS subgroups associated with high risk invasive cancers

## INTRODUCTION

As nonobligate precursors of invasive disease, precancers provide a unique vantage point from which to study the molecular pathways and evolutionary dynamics that lead to the development of life-threatening cancers. Breast ductal carcinoma in situ (DCIS) is one of the most common precancers across all tissues, with almost 50,000 women diagnosed each year in the U.S. alone (American Cancer Society, 2019). Current treatment of DCIS involves surgical excision with either breast conserving surgery or mastectomy, with the goal of preventing invasive cancer. However, DCIS consists of a molecularly heterogeneous group of lesions, with highly variable risk of invasive progression. An improved understanding of which DCIS is likely to progress could thus spare a subgroup of women unnecessary treatment.

Identification of factors associated with disease progression has been the subject of substantial study. Epidemiologic models of cancer progression indicate that clinical features such as age at diagnosis, tumor grade, and hormone receptor expression may have some prognostic value; however, they have limited ability to identify the biologic conditions that govern whether DCIS will progress to invasive cancer. Previous molecular analyses of DCIS have studied either 1) cohorts of pure DCIS with known outcomes (*e.g.,* disease-free versus recurrent), or 2) cross-sectional cohorts of DCIS that either do or do not exhibit adjacent areas of invasive cancer (Allinen et al., 2004, Gil Del Alcazar et al., 2017, Heselmeyer-Haddad et al., 2012, Lesurf et al., 2016, Newburger et al., 2013, Gorringe et al., 2015, Casasent et al., 2018, Abba et al., 2015, Vincent-Salomon et al., 2008a). Both of these approaches have tested potentially divergent assumptions: recurrence of the DCIS as invasive breast cancer (IBC) may arise from neoplastic cells that were left behind when the DCIS was removed, be related to an initial field effect, or may develop from independent events. Longitudinal cohorts have provided a perspective of cancer progression over time. Analysis of DCIS found adjacent to invasive cancer assumes that these preinvasive areas are a good model for pure DCIS tumors and are the ancestors of the invasive cancer cells, with synchronous lesions inferring progression. To date, these studies have not produced clear evidence for a common set of events that are associated with invasion.

This diversity is mirrored in DCIS where few genomic aberrations have been identified that can differentiate DCIS from IBC (Johnson et al., 2012, Heselmeyer-Haddad et al., 2012, Newburger et al., 2013, Gorringe et al., 2015, Yao et al., 2006, Pareja et al., 2020) and microenvironmental processes, including collagen organization, myoepithelial changes, and immune suppression, may contribute to IBC development (Lesurf et al., 2016, Allinen et al., 2004, Gil Del Alcazar et al., 2017). Presently, it remains unknown how these different molecular axes together contribute to DCIS evolution.

Here, as part of the NCI Human Tumor Atlas Network (HTAN) we collected and curated two of the largest DCIS cohorts to date: from the Translational Breast Cancer Research Consortium (TBCRC) 038 study and the Resource of Archival Breast Tissue (RAHBT), on which to conduct multimodal molecular analyses. We performed a comprehensive integrated profile of these complementary, clinically annotated, longitudinally sampled DCIS cohorts, in order to understand the spectrum of molecular changes in DCIS and to identify both tumor and stromal predictors of subsequent events. We used multidimensional and multiparametric approaches to address the central conceptual themes of cancer progression, ecology and evolutionary biology, and molecular subtypes. Multiple data types were applied to create a platform for complex multi-dimensional data representation. We hypothesize that the breast precancer atlas (PCA) presented here will allow for the application of phylogenetic tools that can reconstruct the relationship between DCIS and IBC, the natural history of DCIS, and factors that underlie progression to invasive disease.

## RESULTS

### Study Design and Cohorts

We generated two retrospective case-control study cohorts of patients initially diagnosed with pure DCIS. Each cohort was composed of cases with DCIS with or without a subsequent ipsilateral breast event (iBE, either DCIS or IBC) after surgical treatment. We used identical eligibility criteria for these two cohorts for outcome analysis. The RAHBT cohort used for outcome analysis has 97 cases with median diagnosis at age 53 and median time to recurrence of 40 months. Over half (66.0%) had lumpectomy with radiation, 10.3% had lumpectomy without radiation, and 35% were identified as black. A total of 216 patients were included in the TBCRC cohort, with median diagnosis at age 52 and median time to recurrence of 48 months. More than half (55.5%) had lumpectomy with radiation, 15.3% had lumpectomy without radiation, and 30.0% were identified as black (**Table 1**). **Figure 1** shows an outline of the cohorts and analyses used in this paper. **Table 1** summarizes the RAHBT and TBCRC cohorts used for outcome analysis, **Table S1** summarizes the RAHBT LCM cohort, and **Table S2** summarizes the assays used in this study by cohort.

**Figure 1.**
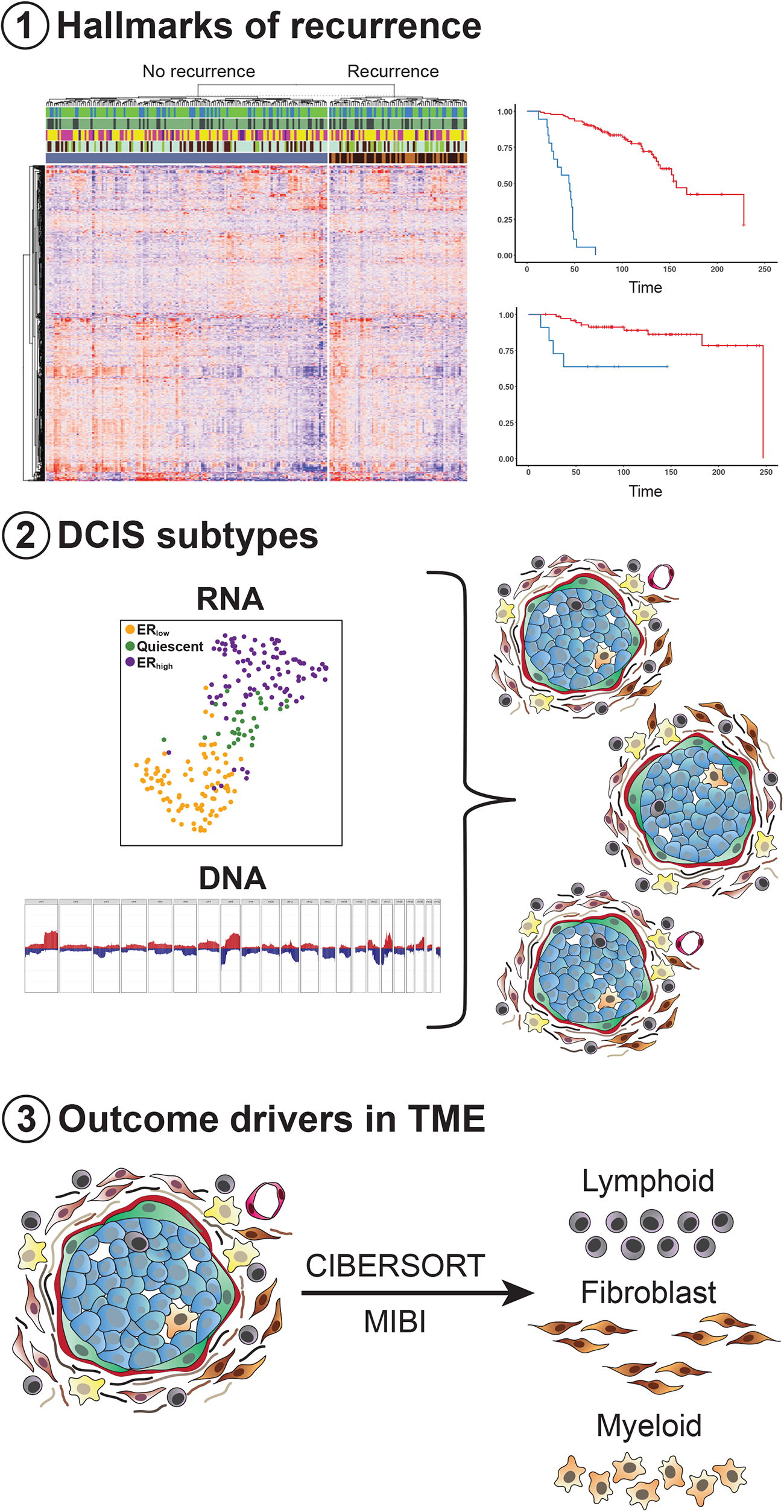
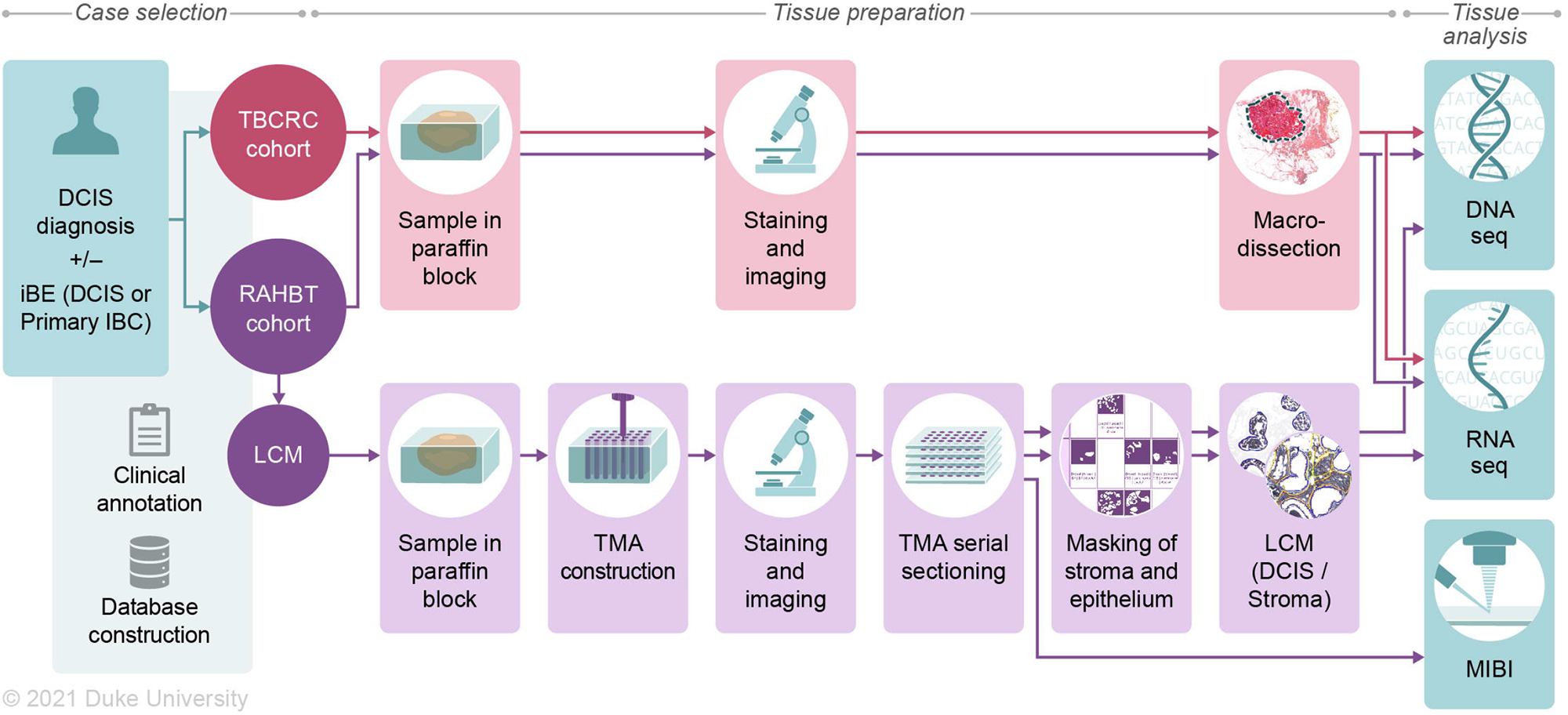
Cohorts and methods Outline. Two retrospective study cohorts (RAHBT, TBCRC) were generated, consisting of DCIS patients with either a subsequent ipsilateral breast event (iBE) or no later events after surgical treatment. TBCRC samples were macrodissected for downstream RNA and DNA analyses. RAHBT samples were 1) macrodissected like TBCRC, or 2) organized into a TMA from which serial sections were made for RNA, DNA and protein (MIBI) analysis (RAHBT LCM cohort). For RNA and DNA sequencing of these samples, TMA cores were laser capture microdissected after tissue masking by a pathologist, to ensure pure epithelial and stromal components.

**Table 1.**
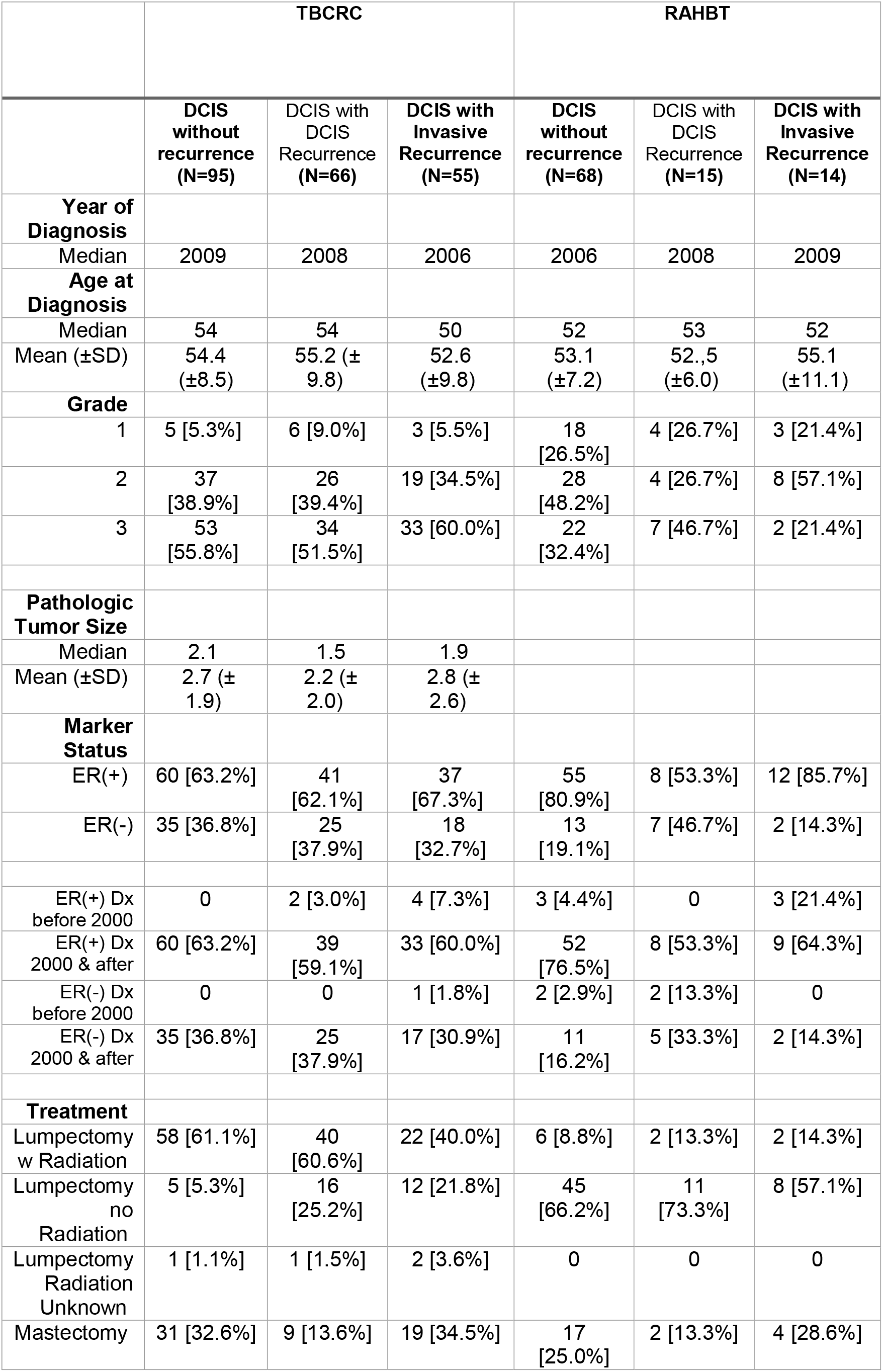
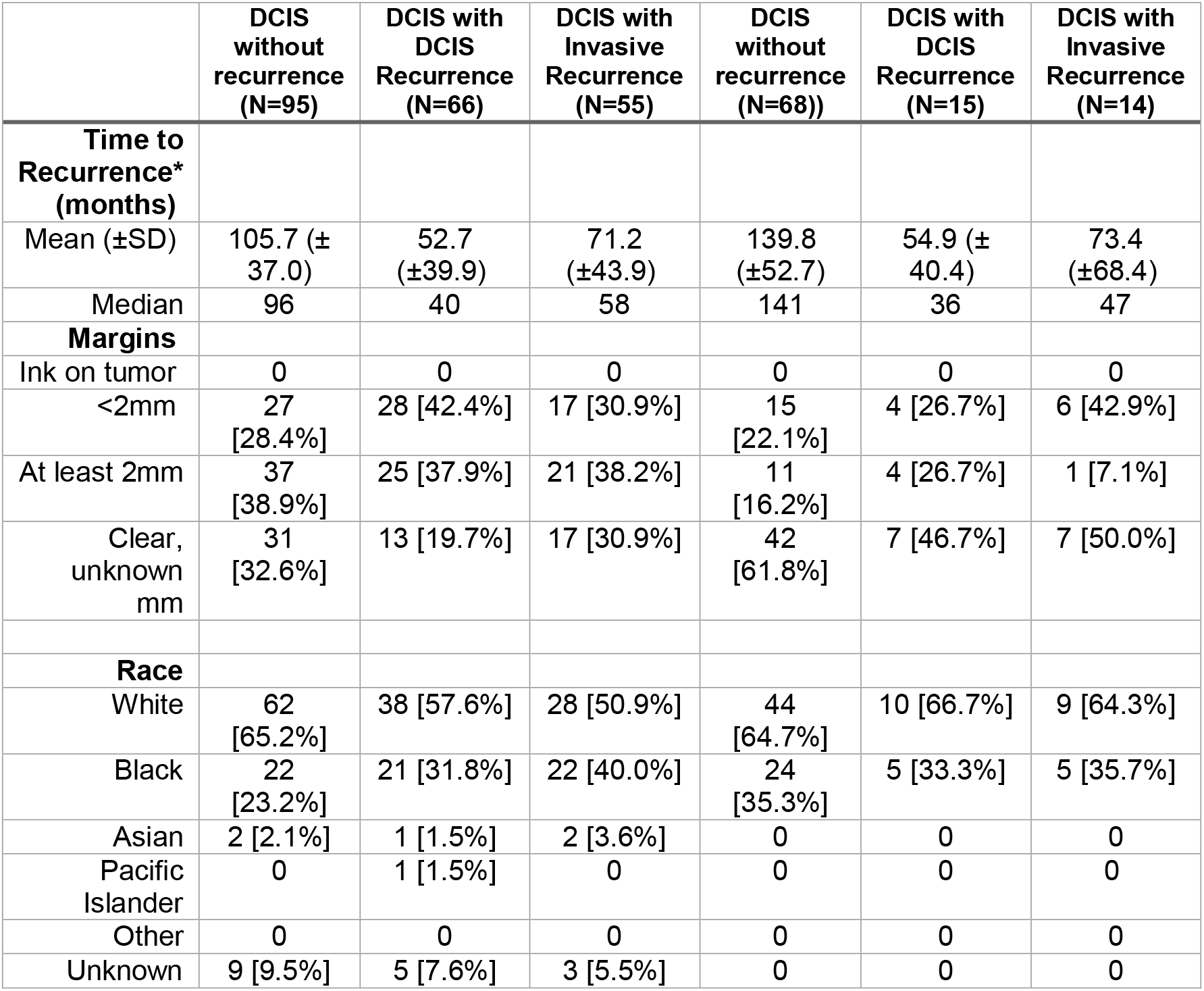
Breast Pre-cancer Atlas Retrospective Patient Cohorts with RNA-seq data and ipsilateral breast event used for outcome analysis. * To end of follow-up for no recurrence.

### Novel 812 gene classifier predicts early recurrence

The TBCRC and RAHBT cohorts were designed to interrogate the biological determinants of recurrence. Specifically, patients with subsequent iBE were matched to patients that did not have any events during long term follow up.

To identify gene expression patterns that correlate with outcome, we analyzed primary DCIS with iBEs within 5 years vs. the remaining samples from the TBCRC cohort, to avoid including non-clonal events that might be more common in later years. We performed RNA-seq expression profiling on RNA from the macrodissected FFPE samples. This analysis identified 812 differentially expressed genes at a false discovery rate of 0.05 (**Figure 2A**, **Table S3**).

**Figure 2.**
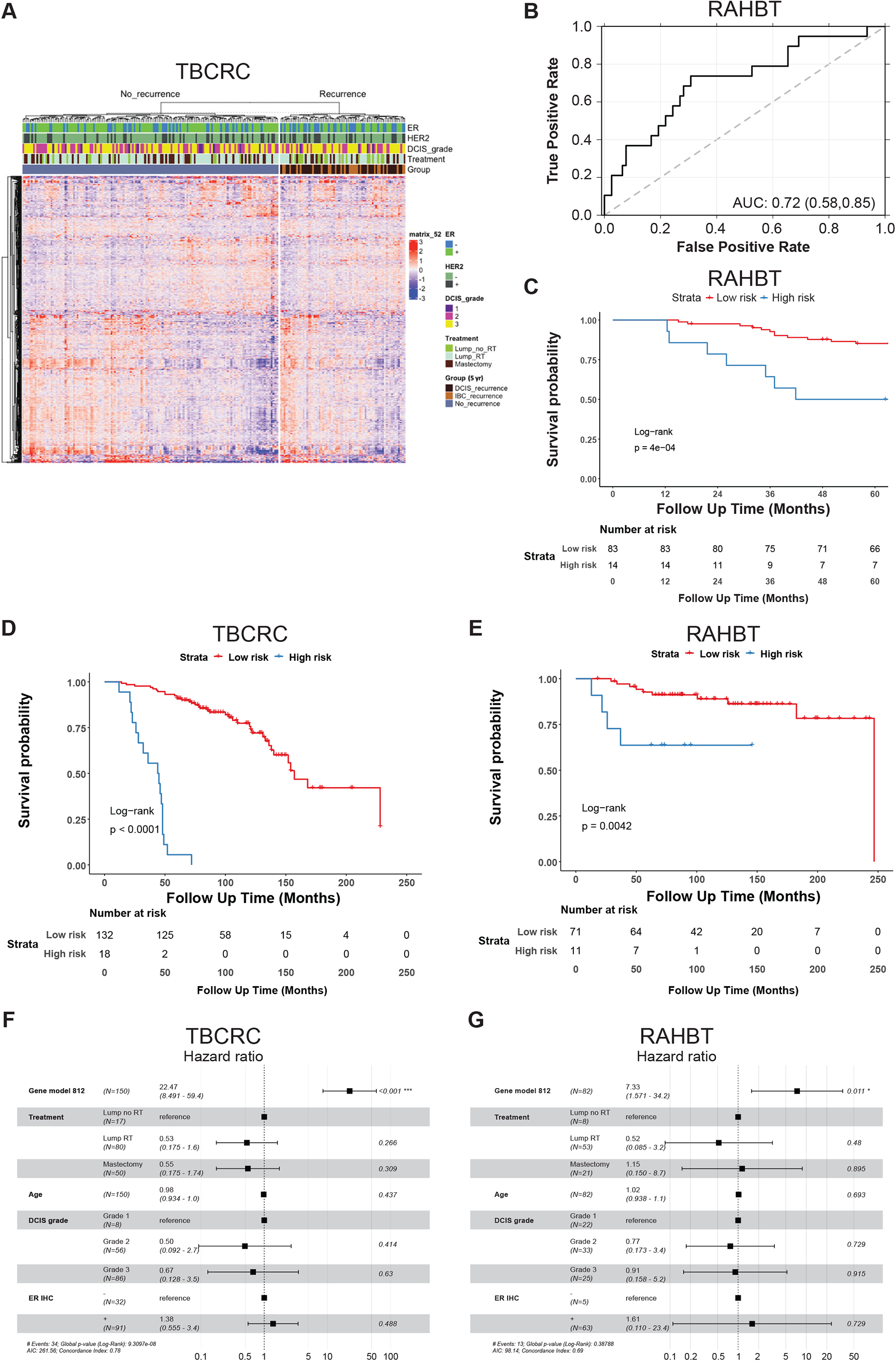
Identification, training and validation of novel 812 gene classifier. **A**) Heatmap of the 812 differentially expressed genes in DESeq2 analysis of cases vs controls (5-year outcome) in TBCRC. Top bars show ER and HER2 status, DCIS grade, treatment, and type of iBE or no iBE. **B**) ROC curve of the 812 gene Random Forest classifier in RAHBT. **C**) Kaplan-Meier plot of time to progression (any iBE, 5-year outcome) stratified by risk groups from the 812 gene classifier in RAHBT. **D-E**) Kaplan-Meier plot of time to invasive progression (full follow-up time) stratified by risk groups from the 812 gene classifier in TBCRC (**D**) and RAHBT (**E**). **C-E**) P-values from log-rank tests. **F-G**) Forest plot of multivariable Cox regression analysis including the 812 gene classifier (high-vs. low-risk groups), treatment, age, DCIS grade, and ER status by IHC for IBC iBEs with full follow-up in TBCRC (**F**) and RAHBT (**G**).

To identify copy number aberrations (CNAs) that correlate with outcome, we performed light pass whole genome sequencing on DNA from macrodissected FFPE samples. We identified 29 recurrent CNAs and performed univariate analysis to determine if any of these had prognostic potential, but found no CNVs that were predictive of recurrence (**Figure S1A**).

Given the absence of significant CNAs, we built our outcome classifier analysis with only the statistically significant differentially expressed genes. Thus, we used the 812 genes to train a Random Forest classifier in the TBCRC cohort. The classifier was then validated in the RAHBT cohort, where the ROC AUC was 0.72 (**Figure 2B**) with Precision 0.86, Recall 0.91, and F1 score 0.88, indicating that the classifier performs well also in the test cohort. The classifier was a significant predictor of any subsequent iBE in both cohorts (RAHBT P=0.0004, **Figure 2C**). Importantly the classifier was also a significant predictor of invasive iBEs over the full follow-up time (TBCRC P <0.0001, RAHBT P = 0.0042, **Figure 2D-E**), demonstrating that it could specifically identify DCIS that progress to the invasive stage.

Next, we examined whether the 812 gene classifier remained an independent predictor of outcome when combined with clinical features. We performed multivariable Cox regression analysis including the 812 gene classifier, treatment, age, clinical ER (IHC based), and DCIS grade (**Figure S1B-C**). Multivariable analysis demonstrates a trend for treatment type and ER status for outcome. However, in this analysis the 812 gene classifier was the only variable that was significant in both cohorts (RAHBT HR = 3.48, (95% CI: 1.14 – 10.6), P = 0.028). Importantly, in multivariable analysis for invasive iBEs only, the classifier showed an even stronger prognostic value in both cohorts, with a hazard ratio of 7.33 in the RAHBT cohort (95% CI: 1.57 – 34.2, P = 0.011, **Figure 2F-G**). Of note, Kaplan-Meier analysis of clinical ER status (IHC-based) demonstrated a trend in the RAHBT cohort (P = 0.053), but not in the TBCRC cohort (P = 0.2, **Figure S1D-E**). Moreover, the 812 gene classifier showed no prognostic value for progression free disease or overall survival for 1064 IBCs from The Cancer Genome Atlas (TCGA,(Liu et al., 2018) (**Figure S1F-I**), suggesting that the classifier is specific for the DCIS stage.

The 812 gene classifier likely represents several distinct biologic processes that promote recurrence and progression to invasion. To further understand the biology and identify pathways involved in recurrence, we performed Gene Set Enrichment Analysis (GSEA) on the differentially expressed genes between cases that recur within 5 years and DCIS that did not in the TBCRC cohort We identified 11 Hallmark pathways significantly associated with early recurrence including those associated with proliferation, immune response, and metabolism (**Figure S1J**).

### DCIS RNA clustering defines expression modules that drive outcome

Since proliferation and metabolism were found to be important pathways in recurrence, we examined whether major phenotype patterns in DCIS drive these pathways. Previous findings have suggested that invasive breast cancer subtypes do not fit well for DCIS (Bergholtz et al., 2020). We hypothesized that a classification scheme based on analysis of DCIS rather than IBC would better address DCIS biology. Thus, to investigate the fundamental biology behind the observed outcome with emphasis on epithelial pathways, we performed unsupervised clustering of RNA sequencing of macrodissected TBCRC samples (n=216) as well as an additional group of RAHBT cases (n=265, **Table S1**) on which laser capture microdissection (LCM) was applied to generate epithelial-enriched samples to evaluate tumor cell expression patterns without the contributions from the surrounding TME (**Figure S2A-F**).

We performed non-negative matrix factorization (NMF) on all protein coding genes (GENCODE v33) with non-zero variance, evaluated the fit of 2-10 clusters and selected a novel three cluster solution based on optimization of silhouette width, cophenetic value, maximizing cluster number and replication in RAHBT (**Figure S2G-H**). The NMF analysis comparing these two datasets found that a 3-cluster solution most reproducibly captured the biologic subgroups. In both the TBCRC and RAHBT LCM cohorts, cluster 1 had significantly higher levels of *ERBB2* and lower levels of *ESR1* compared to clusters 2 and 3 (**Figure 3A, B**), which both had increased *ESR1* expression. We referred to the three clusters as ER_low_, quiescent, and ER_high_ respectively. To characterize the three clusters, we conducted differential abundance analysis comparing each cluster individually to the other two combined (*i.e.,* one-vs-rest). The deregulated pathways in each cluster were highly concordant across TBCRC and RAHBT LCM cohorts, further supporting three transcriptional patterns in DCIS that are driven by the tumor cell compartment (P_ERlow_ = 2.33×10^-2^; P_quiescent_ = 8.37×10^-2^; P_ERhigh_ = 9.20×10^-10^; hypergeometric test; **Figure S2K**).

**Figure 3.**
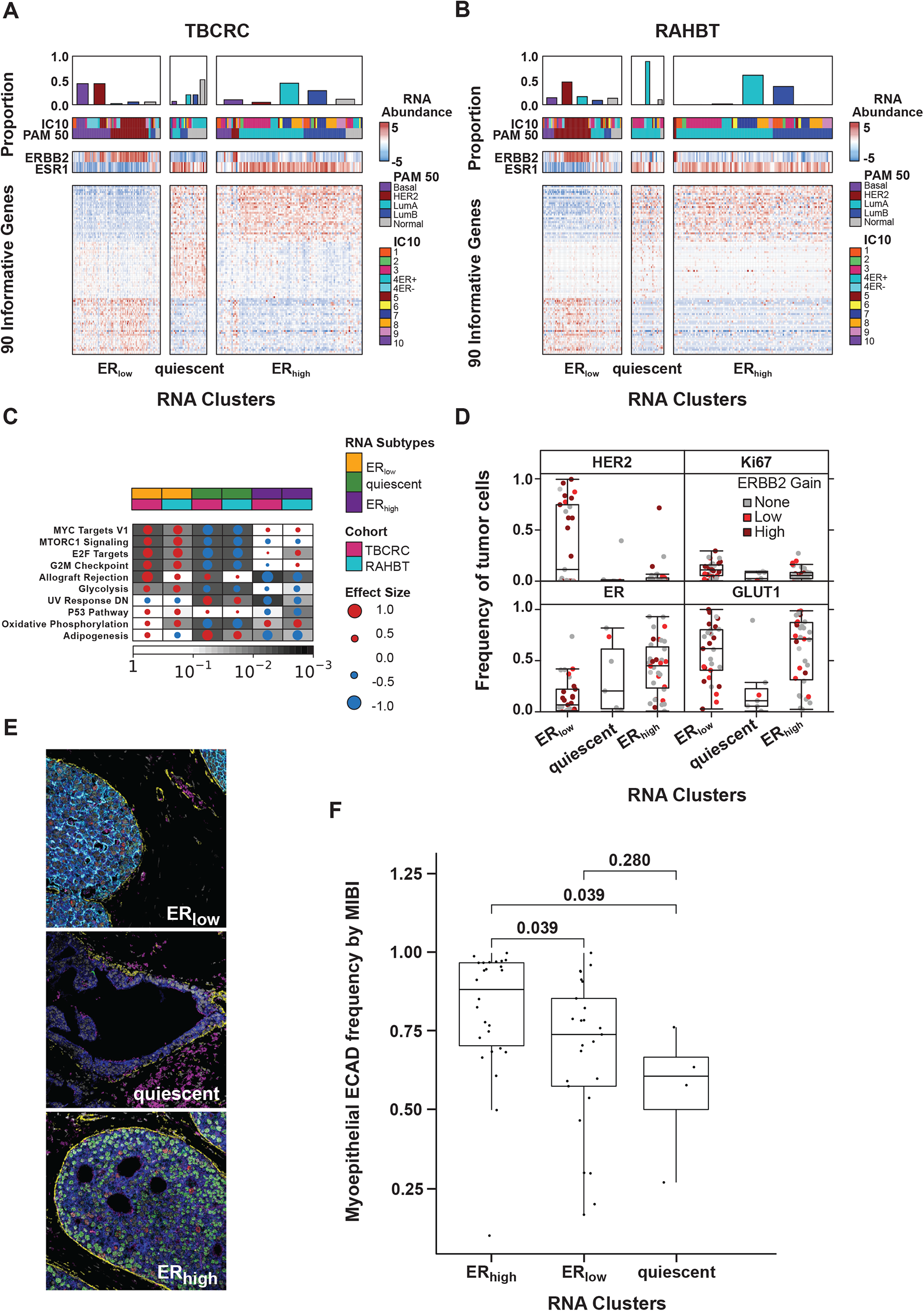
*De novo* transcriptomic DCIS subtypes correlate with outcome pathways. **A)** Unsupervised clustering of DCIS transcriptomes identifies three subtypes: ER-, quiescent and ER+. Heatmap depicts RNA abundance of 90 informative genes, y-axis, contributing to the three subtypes in TBCRC samples, x-axis. Barplot along the top indicates the proportion of PAM50 subtypes within each cluster. Covariates indicate integrative and PAM50 subtypes, along with *ERBB2* and *ESR1* mRNA abundance for each sample. **B)** Heatmap of DCIS subtypes in RAHBT. **C)** GSEA Hallmark pathway analysis of each cluster vs rest for TBCRC and RAHBT LCM showing outcome-associated pathways only. Pathways, y-axis, deregulated in each cluster are highly concordant in TBCRC and RAHBT. Size of the dot and color represents the magnitude and direction of pathway deregulation, *i.e.* blue indicates the pathway is downregulated while red indicates the pathway is upregulated. Background shading indicates the false discovery rate. Covariate along the top indicates the DCIS subtype and cohort. **D)** The ER-subtype is associated with more HER2+ tumor cells, as determined by MIBI, while the ER+ and quiescent clusters are associated with more ER+ tumor cells. The quiescent cluster has a low frequency GLUT1 and Ki67 positive cells. Dot color indicates *ERBB2* genomic amplification level. **E)** Representative MIBI images of the three subtypes in (**B**). White = Nuc; Blue = PanKRT; Yellow = SMA; Pink = GLUT1; Cyan = HER2; Green = ER; Red = Ki67. **F**) Boxplot of myoepithelial ECAD frequency by MIBI shows enrichment in the ER high subtype.

While we observed a differential expression of the estrogen response in the ER_high_ cluster vs ER_low_ cluster, the most striking biologic patterns involved the pathways associated with DCIS recurrence (**Figure 3C, Figure S2K**). Some major patterns were evident. First, pathways including MYC, mTOR signaling, and cell cycle pathways were enriched in the ER_low_ cluster and significantly depleted in the quiescent cluster. Second, the Allograft Rejection, p53 and Adipogenesis pathways were high in the ER_low_ cluster and low in the ER_high_ cluster. Finally, the ER high cluster was depleted for the UV Response Down and enriched for the Oxidative Phosphorylation pathways, both of which were associated with recurrence. Of note, none of the recurrence associated pathways were enriched in the Quiescent cluster. The presence of the Allograft Rejection pathway in the RAHBT LCM epithelial samples, though not significant, suggests that immune cells have infiltrated into the epithelial compartment in the involved samples. Thus, the 3-cluster solution identified pathways associated with early recurrence.

Genomic and transcriptomic-based classifications of IBC (Perou et al., 2000, Curtis et al., 2012) have characterized the spectrum of biologic subtypes in invasive breast cancer, but it remains unclear whether these classification schemes accurately describe the spectrum of the DCIS stage. To answer this question, we applied the PAM50 classification to TBCRC and RAHBT LCM epithelial DCIS samples and evaluated the correlation of each sample to the centroid of its assigned subtype and compared the correlations against those observed in 1109 IBCs from TCGA. The median correlation for basal-like DCIS samples was significantly lower than basal-like IBC samples (P = 8.01×10^-16^; Wilcoxon rank sum test, **Figure S2L**), as previously shown (Bergholtz et al., 2020). Significantly decreased correlation was also observed for luminal A (P = 3.13×10^-3^) and normal-like subtypes (P = 6.21×10^-3^). Projecting the DCIS transcriptome onto two-dimensions using UMAP revealed clear deviations from the PAM50 centroids (**Figure S2M-N**). Moreover, PAM50 failed to robustly predict recurrence in DCIS (**Figure S2O-P**). These data suggest that while established IBC subtypes can be identified in DCIS, they do not fit DCIS as robustly as IBC, and are not prognostic in these premalignant lesions.

In support of the 3-cluster solution, we investigated protein expression by MIBI for a subset of these patients (n=71). The frequency of ER+ tumor cells was significantly higher in the quiescent and ER_high_ subtypes compared to ER_low_ (log_2_FC = 2.73; P= 2.11×10^-5^; Wilcoxon rank sum test) while HER2+ tumor cells were significantly higher in the ER_low_ subtype (log_2_FC = 4.88; P= 3.74×10^-2^; Wilcoxon rank sum test; **Figure 3D**). In general, the frequencies of ER+ and HER2+ tumor cells were well correlated with RNA abundance of *ESR1* and *ERBB2*, respectively (**Figure S2Q-R**). *PGR* levels were similarly upregulated in quiescent and ER_high_ compared to ER_low_ (log_2_FC_quiescent_ = 1.01; P_quiescent_ = 6.28×10^-3^; log_2_FC_ERhigh_ = 1.89; P_ERhigh_ = 4.43×10^-6^; DESeq2; **Figure S2S**). Based on our MIBI data, the quiescent lesions were depleted for Ki67 (log_2_FC = -1.46; P= 8.08×10^-2^; Wilcoxon rank sum test) and GLUT1 (log_2_FC = -2.64; P= 8.47×10^-3^) positive tumor cells, compared to ER_high_ and ER_low_ tumors, suggesting quiescent lesions are less proliferative and less metabolically active (**Figure 3D-E**).

In their analysis of DCIS tumors and TME by MIBI, Risom *et al*. identified high myoepithelial E-cadherin expression as the most discriminative feature for risk of progression (Figure 6A-B in (Risom et al., 2022). To further investigate this in relation to the identified RNA clusters, we compared the distribution of myoepithelial E-cadherin frequency by MIBI in matched RAHBT LCM RNA samples. We found that the ER high cluster had significantly higher myoepithelial E-cadherin frequency by MIBI compared to both the ER low and quiescent clusters (P ≤ 0.025, **Figure 3F**). While most recurrence-associated pathways were enriched in the ER low cluster, this points to a feature associated with recurrence amongst DCIS tumors with high ER expression, and highlights that there are multiple paths to progression in DCIS.

### Amplifications characteristic of high-risk of relapse IBC occur in DCIS

Next, we investigated CNAs in DCIS, and how these contribute to the pathways associated with early recurrence. We analyzed the genomic landscape of DCIS in DNA samples from TBCRC and RAHBT combined (n=228), and identified 29 recurrent CNAs, 13 gains and 18 losses. These occurred in 10.1-52.6% of the DCIS samples (FDR < 0.05; GISTIC2; **Figure 4A**). The identification of these 29 common CNAs was not biased by depth of sequencing, but two CNVs were associated with cohort (1p21.3 and 10p15.3 deletions, **Table S4**). The most frequent alterations were gain of chromosome 1q and 17q, including 17q12 where the *ERBB2/HER2* oncogene is located, and loss of chromosome 17p, 16q, and 11q (**Figure 4A**), confirming prior findings (Yao et al., 2006, Lesurf et al., 2016, Abba et al., 2015, Trinh et al., 2021) and notably reflecting the CNA landscape of IBC (Russnes et al., 2010, Curtis et al., 2012).

**Figure 4.**
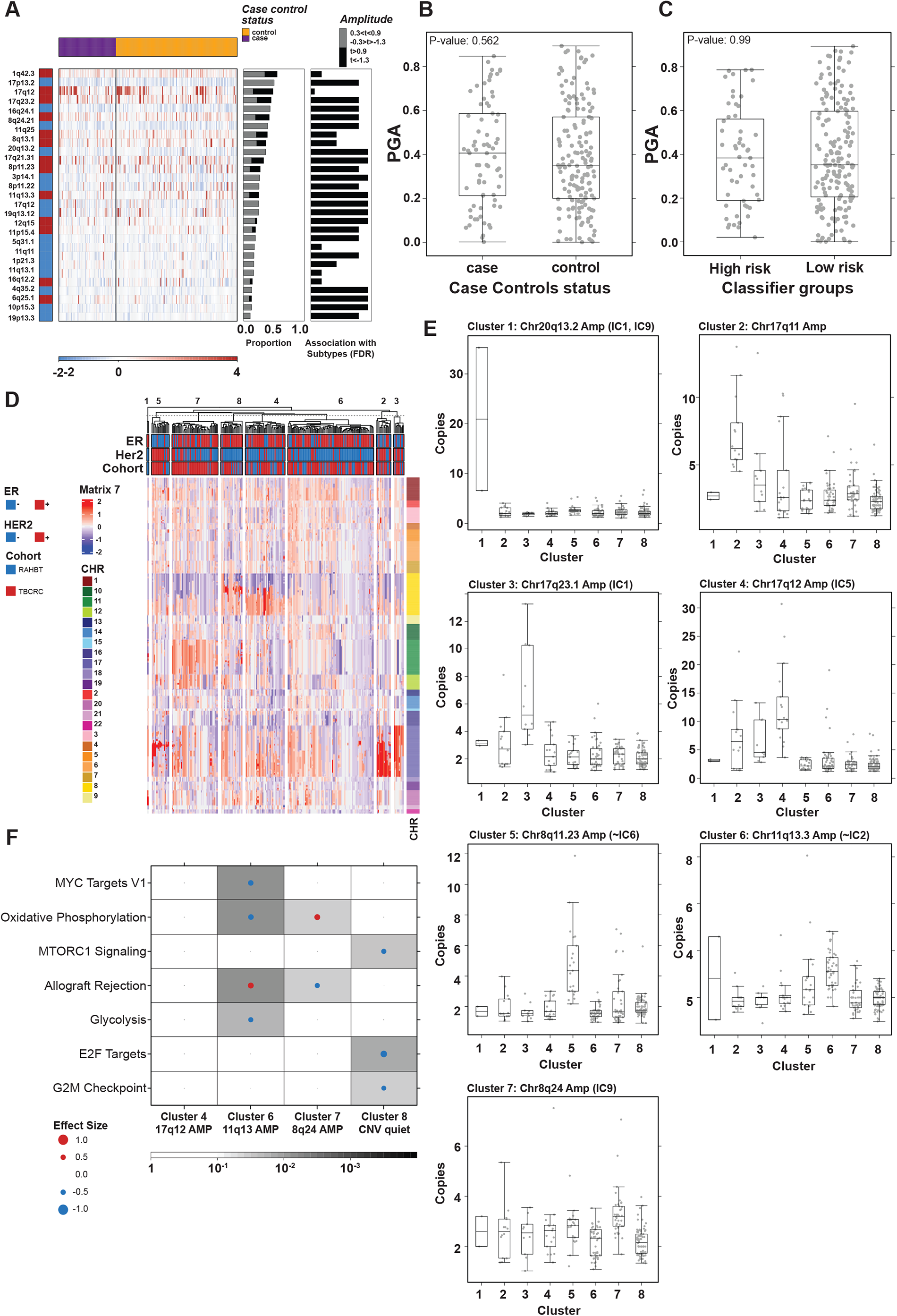
Characteristic invasive breast cancer CNAs present in DCIS. **A)** Twenty-nine cytobands are significantly recurrently altered in DCIS. Heatmap shows log_2_ copy number for each of the recurrent CNAs, y-axis, in each sample, x-axis. Samples are grouped by 5-year outcome groups as indicated by the covariate along the top. The middle barplot shows the proportion of samples with each CNA. Grey and black represent increasing amplitudes. The barplot on the right shows the FDR from a Kruskal-Wallis test of each CNA with the risk groups. No CNAs were significantly associated with outcome groups. **B-C)** No significant association was observed between PGA and 5-year outcome groups (**B**) or classifier risk groups (**C**). P-value from Kruskal-Wallis test. Boxplot represents median, 0.25 and 0.75 quantiles with whiskers at 1.5x interquartile range. **D)** Unsupervised clustering of CNA landscape identifies eight clusters. Cluster1: 20q13 Amp, Cluster2: 17q11 Amp, Cluster3: 17q23 Amp, Cluster4: 17q12 Amp (ERBB2), Cluster5: 8p11 Amp, Cluster6: 11q13 Ap, Cluster7: 8q24 Amp (MYC), Cluster 8: CNV quiet. Heatmap depicts log_2_ copy number of genomic segments, y-axis, in TBCRC and RAHBT samples, x-axis. Covariates indicate ER and HER2 status (RNA derived) for each sample. Covariate along the right shows the chromosome of each segment. **E)** Boxplot shows log_2_ copy number, y-axis, across the eight clusters, x-axis. **F**) GSEA Hallmark pathway analysis of differentially expressed genes in matched RNA samples (DESeq2 each-vs-rest) by DNA cluster for TBCRC and RAHBT, showing outcome-associated pathways only.

Next, we investigated if the distribution of the Proportion of the Genome copy number Altered (PGA) was biased in the 5-year outcome groups or 812 gene classifier risk groups, but found no significant differential distribution (**Figure 4B-C**). PGA was not correlated to sequencing depth, nor predictive of iBEs (**Figure S3A-B**).

Early patterns of alterations may provide insight into the mechanisms by which neoplastic lesions form and progress towards invasion. To identify genomic subtypes emerging in DCIS, we employed unsupervised NMF clustering on CNA segments on TBCRC and RAHBT jointly and identified eight CNA clusters ranging in size from 2-98 samples (**Figure 4D**; **Figure S3C-D**). The clusters were not biased by depth of sequencing (**Figure S3E**). CNA cluster 1 was characterized by amplification of chr20q13.2 (**Figure 4E**). Three clusters were characterized by chr17q amplification (Cluster 2: 17q11, Cluster 3: chr17q23.1, Cluster 4: chr17q12).

Cluster 5 was characterized by amplifications of chr8p11.23, Cluster 6 by chr11q13.3 amplification, and Cluster 7 by amplification of *MYC* on chr8q24. Finally, Cluster 8, the largest group (n=98), represented a CNA quiet subgroup, characterized by the absence or diminished signal of the aforementioned CNAs.

Integrative subgroups (ICs) is a classification scheme developed for IBC based on integration of genomic copy number and expression profiles (Curtis et al., 2012). Intriguingly, despite the eight CNA clusters not being associated with recurrence (**Figure S3F-G**) several of the eight CNA subtypes could be attributed to the presence or absence of CNAs characteristic of the IC subtypes, namely the four high-risk of relapse ER+/HER2-subgroups (IC1,2,6,9) and the HER2-amplified (IC5) subgroup (Rueda et al., 2019) (**Figure 4E**). Of note, these four high-risk integrative subgroups (IC1,2,6,9) account for 25% of ER+/HER2-IBC and the majority of distant relapses (Rueda et al., 2019). Integrative subtypes are prognostic in IBC and improve the prediction of late relapse relative to clinical covariates. Understanding the clinical course of DCIS lesions harboring these high-risk invasive features is highly relevant in refining clinically meaningful risk associated with DCIS progression.

To identify enriched pathways in the eight CNA clusters, we investigated the differential gene expression in matched RNA samples (DESeq2 one-vs-rest) and performed GSEA Hallmark analysis on the resulting gene lists. Here, DNA Clusters 6 (chr11q13 amplification) and 7 (chr8q24 (*MYC*) amplification) were enriched for pathways associated with early recurrence (allograft rejection and oxidative phosphorylation, respectively). In addition, cluster 8, the CNA quiet cluster was depleted for pathways associated with early recurrence (cell cycle and mTORc1 signaling), and cluster 6 was depleted for MYC targets (**Figure 4F, Figure S3H**). The remaining four CNA clusters had no significant pathway enrichments. Thus, we identified a DNA-based cluster solution characterized by amplifications seen in high-risk IBC subtypes (IC10), including 17q12 (*ERBB2*) and 8q24 (*MYC*) amplification, some of which were significantly enriched or depleted for pathways associated with early recurrence.

### The DCIS TME is heterogeneous and reflects distinct immune and fibroblast states

The Hallmark pathways identified represent a diverse set of biologic events and may involve different components of the DCIS ecosystem including the cells within the TME. Accumulating evidence has shown that the tumor microenvironment (TME) is crucial for cancer development and progression (Hinshaw and Shevde, 2019, Gil Del Alcazar et al., 2020). To analyze the DCIS tumor microenvironment (TME), we generated RAHBT LCM stromal samples by dissecting stromal tissue from the DCIS edge (**Figure S2D-F**).

Next, to identify the contribution of epithelial and stromal components to the 812 gene classifier, we performed differential gene expression analysis between stromal (n=196) and epithelial (n=265) primary DCIS RNA samples from the RAHBT LCM cohort. We identified 9748 DE genes (adj. P<0.05) between epithelial and stroma, 5161 of which were predominantly expressed in the epithelial compartment, and 4587 which were mainly expressed in the stroma. An analysis of the 812 genes in the classifier showed that 20% of these significant genes were expressed primarily in the stromal/TME cells, whereas 34% were genes predominantly expressed in epithelium (**Table S3**).

The MIBI method provides an orthogonal view of the TME and gives us protein expression and cell identity, enabling us to both validate and extend our observations from the RNA expression analysis. We took tissue sections from adjacent slides to the same sections used for RNA-seq in order to spatially align the same ducts for both MIBI and RNA. We used the MIBI panel as mentioned above, which consists of 37 metal-conjugated antibodies, that capture 16 different cell types including epithelial, fibroblasts, and immune cell types (Risom et al., 2022). We can compare the cell type distribution as defined by MIBI across the DCIS samples with the inferred cell type distribution from the RNA expression data using CIBERSORTx (CSx, see **Methods, Figure S4A-B**). This approach allowed us to cross-validate findings and extend our observations on cell composition to DCIS samples that do not have MIBI results, including samples from the TBCRC cohort.

In order to define discrete TME phenotypes, we performed a shared nearest neighborhood (SNN) analysis of the stromal RNA expression data and identified four distinct DCIS-associated stromal clusters (**Figure 5A**) and differentially expressed genes (DESeq2 each-vs-rest, **Figure 5B**). Pathway analyses (**Figure 5C, S4C**), MIBI protein expression and cell type distribution (**Figure 5D**), and inferred cell type distribution by CSx (**Figure 5E, Figure S4D-G**) were used to describe the major characteristics of each cluster. Based on these results we termed the clusters Immune, Desmoplastic, Collagen-rich, and Normal. **Figure 5F** shows a representative MIBI image of each cluster, where a strong correlation with fibroblast states and immune cell density is observed.

**Figure 5.**
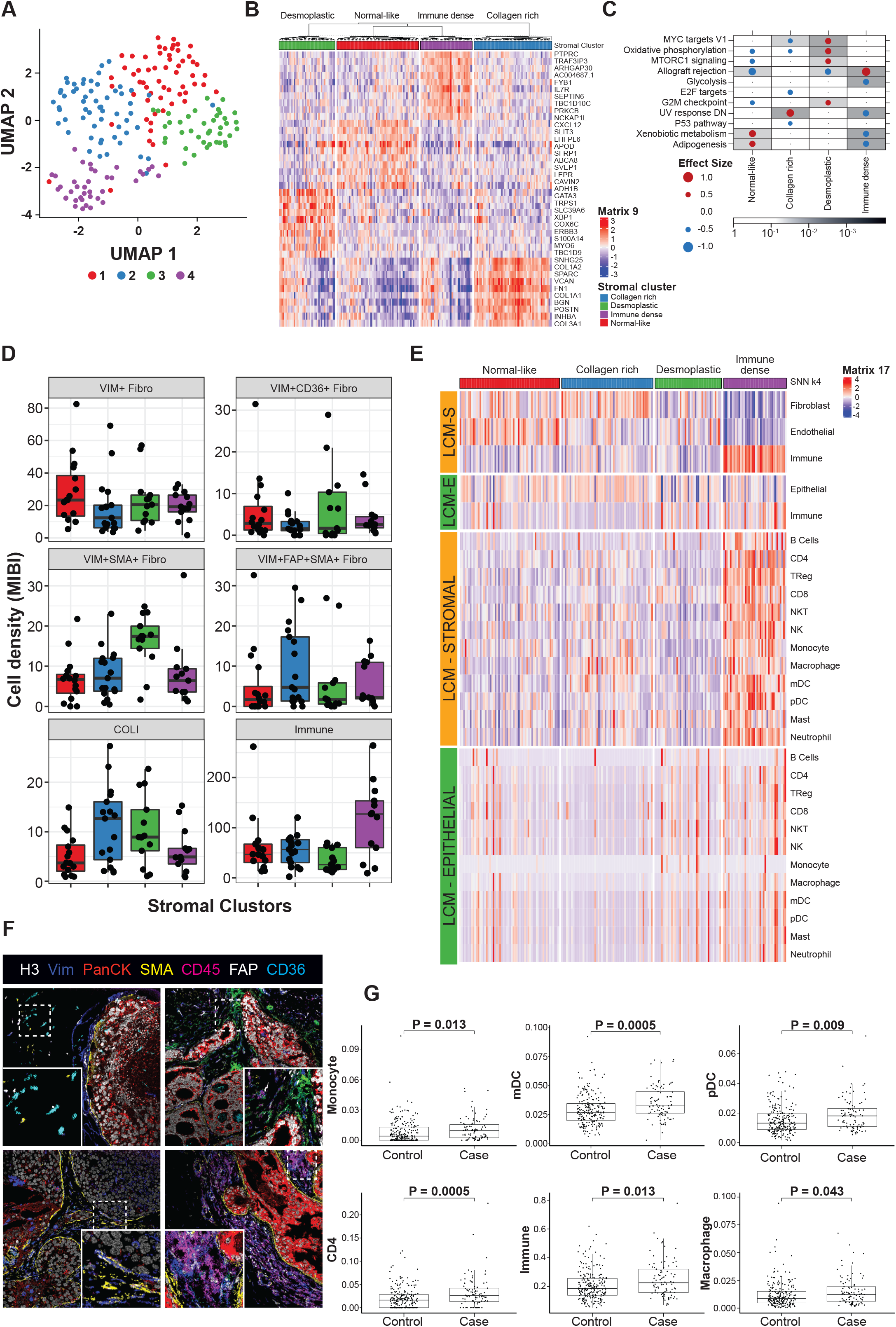
Analysis of the tumor microenvironment. **A)** UMAP projection of DCIS stromal transcriptome colored by the four identified clusters. **B)** Heatmap of the top 10 up-regulated genes for each stromal cluster (DESeq2 each-vs-rest). **C)** GSEA Hallmark pathway analysis of DE genes in each stromal cluster vs rest in RAHBT LCM stromal samples, showing outcome-associated pathways only. **D)** MIBI-estimated cell density within stromal clusters supports CSx findings (total n_MIBI_ = 59, n_CSx_ = 193). **E)** Deconvolution analysis by CSx of epithelial and stromal LCM samples grouped by stromal clusters shows different immune cell and fibroblast abundance in DCIS stromal clusters. **F)** Representative MIBI images of stromal clusters reflecting different fibroblast states and total immune density. Top left: normal-like. Top right: Collagen rich (FAP+). Bottom left: Desmoplastic (SMA+). Bottom right: Immune dense (CD45 high). H3, histone 3; VIM, vimentin; panCK, pan cytokeratin; SMA, smooth muscle actin; FAP, fibroblast activated protein. **G)** CSx inferred cell type distribution between cases with iBEs vs controls (TBCRC and RAHBT combined). Only cell types with significant differences (adj.P<0.05) are shown. mDC: myeloid dendritic cells. pDC: plasmacytoid dendritic cells.

The Immune stromal cluster was the most distinct of the stromal subtypes. It was defined by enrichment for the outcome-associated allograft rejection pathway and other pathways representing immune activation. In addition, MIBI and CSx data demonstrated that total abundance of immune cells was more than twice that of the other clusters, with predominance of lymphoid over myeloid cell populations. Within this cluster, a subgroup of samples was highly enriched for B cells, whereas another subgroup showed an overall balanced immune cell type composition. The Immune cluster also showed association with MIBI-identified T-cell and B-cell enriched neighborhoods (see Risom et al., 2022 for details), as well as myoepithelial- and myeloid-enriched neighborhoods (**Figure S4H**).

The normal-like stromal cluster was enriched for Gene Ontology (GO) pathways involved with ECM organization, complement and coagulation cascades, focal adhesion, and PI3K-AKT signaling. The collagen-rich stromal cluster was characterized by GO pathways associated with collagen metabolism, TGFb signaling, and proteoglycans in cancer, and shared genes involved in cell-substrate and focal adhesion with the ‘normal-like cluster. By predicting frequencies of cell types present in each sample we found that this cluster had the highest fibroblast abundance and total myeloid cells, mostly associated with macrophages and myeloid dendritic cells (mDC). MIBI results showed that this cluster was enriched in collagen and fibroblast associated protein positive (FAP+, VIM+, SMA+) myofibroblasts. The desmoplastic stromal cluster was represented by mammary gland development and fatty acid metabolism, high presence of CD8+ T cells assessed by CSx, and myofibroblasts by MIBI (VIM+, SMA+ myofibroblasts).

From the stromal cluster analysis, it is clear that the immune response is present in a discrete subset of cases. However, analysis of outcome related to stromal subtype demonstrates only a modest outcome difference, and there is no major contribution from the immune subcluster (P = 0.12, log-rank test, **Figure S4I**). We hypothesized that rather than the entire immune response, the outcome differences caused by the immune response can be attributed to a subset of immune cells. To address this, we analyzed CSx inferred cell type distribution in 5-year outcome groups. Using the CSx results allows us to leverage the entire cohort of TBCRC and RAHBT combined. We identified significantly higher levels of CD4+ T cells, myeloid- and plasmacytoid dendritic cells, monocytes, macrophages, and overall immune cells in cases vs. controls (**Figure 5G**). No significant difference was found for the remaining CSx cell types in this analysis (data not shown). Furthermore, in univariate Cox regression analysis we found that several cell types, including CD4 T cells, myeloid- and plasmacytoid dendritic cells, were significant predictors of any iBE at 5 years after treatment (**Table S5**). These differences in outcome groups were overall mirrored by the CSx inferred cell type distribution in the high- and low risk classifier groups (**Figure S4J**). Finally, we investigated the distribution of CSx cell types in cases with invasive and DCIS iBEs as well as controls 5 years after treatment. The results overall reflected the analysis in cases vs. controls, and the largest differences were observed between IBC iBEs and controls (**Figure S4K**).

Taken together, these analyses support the contributions of individual immune cells with high-risk outcomes. However, non-immune cell phenotypes are not well defined by this CSx approach but can still be identified as a biologic response. The stromal signature with the clearest and most favorable outcome stratification was the Desmoplastic cluster (HR = 0.23, P=0.06, **Figure S4J**), despite being enriched for several early recurrence pathways, including proliferative signals (MYC and G2M checkpoint) which are otherwise associated with bad outcome in the epithelial compartment. This highlights the complexity and the differential contribution from the stromal and epithelial compartments.

## DISCUSSION

The aims of the HTAN Breast Pre-Cancer Atlas project are to 1) develop a resource of multi-modal spatially resolved data from breast pre-invasive samples that will facilitate discoveries by the scientific community regarding the natural history of DCIS and predictors of progression to life-threatening IBC; and 2) populate that platform with data from retrospective cohorts of patients with DCIS and demonstrate its use to construct an atlas to test novel predictors of progression. In the current study, we examined two well-annotated, retrospective, longitudinal patient cohorts (TBCRC and RAHBT) with or without a subsequent iBE. Together, these cohorts comprise one of the largest series of matched case control samples to date, allowing greater statistical power to perform the comprehensive genomic studies reported here. A particular strength of the study is the complementary nature of the two cohorts, which have allowed for validation of our findings, as well as the capability to separately study the epithelial and stromal components in the RAHBT LCM samples.

DCIS is a heterogeneous disease with variable prognosis, but has defied attempts to identify molecular factors associated with future progression. Previous studies have evaluated the prognostic value of specific biomarkers associated with outcomes, with conflicting conclusions for virtually all markers tested, including ER, HER2, immune markers such as tumor infiltrating lymphocytes, and stromal characteristics. Many promising leads have not been reproducible due to multiple factors, including lack of endpoint standardization, important differences between cohorts, small sample size, and limited datasets for validation with well annotated long-term outcomes.

Herein, we have developed and validated a novel 812 gene classifier which independently predicted risk of both overall recurrence and invasive progression. This classifier was highly associated with outcome in a multivariable model which included treatment, age, grade, and clinical ER status; the genomic classifier had a HR of 22.5 (95% CI 8.5 to 59.4) in the training set and 7.3 (95% CI 1.6 to 34.2) in the validation set, over four-fold higher than has been previously reported for other prognostic markers for DCIS (Kerlikowske et al., 2010).

Importantly, we found that the genomic classifier was a stronger predictor of 5-year recurrence or progression than previously described factors, including age at diagnosis, tumor grade, ER status, or treatment. The large dataset, with a high number of events, permitted an agnostic analysis of all genome-wide features and was thus less opportunistic than other, more limited studies. Further, since no a priori assumptions were made regarding whether to incorporate the molecular features of invasive cancer, we were able to construct a less biased genomic predictor.

Our novel classifier is characterized by a number of hallmark pathways including several related to cell cycle progression and growth factor signaling (E2F targets, G2M checkpoint, MYC targets, MTORC1 signaling) and metabolism (Glycolysis and Oxidative Phosphorylation). Examination of pathway activation status at the individual tumor level reveals the underlying complexity of the signature (**Figure 6A-C**). High correlation between cell cycle linked E2F and G2M pathways are consistent with a proliferation related signature. However the strongest features of the signature (distinguishing cases from controls) are MYC and MTORC1 signaling which are strongly correlated with each other but less so with the canonical proliferation pathways indicating that proliferation alone is not the central predictor. The three other pathways that are most profoundly different between cases and controls relate to metabolism and immunity. Interestingly, both glycolysis and oxidative phosphorylation are increased in cases suggesting that heightened metabolic activity is associated with risk of progression regardless of whether it is anaerobic. Further, there is significant positive correlation between these two pathways indicating that metabolically active tumors use both pathways (**Figure 6C**). Finally, allograft rejection, a broad immune pathway, is elevated in cases and in general appears to be an independent component of the signature. Overall, there are multiple components to this signature that are elevated in different subsets of the tumors lending additional evidence that simplified predictors fail to capture the heterogeneity of the disease.

**Figure 6.**
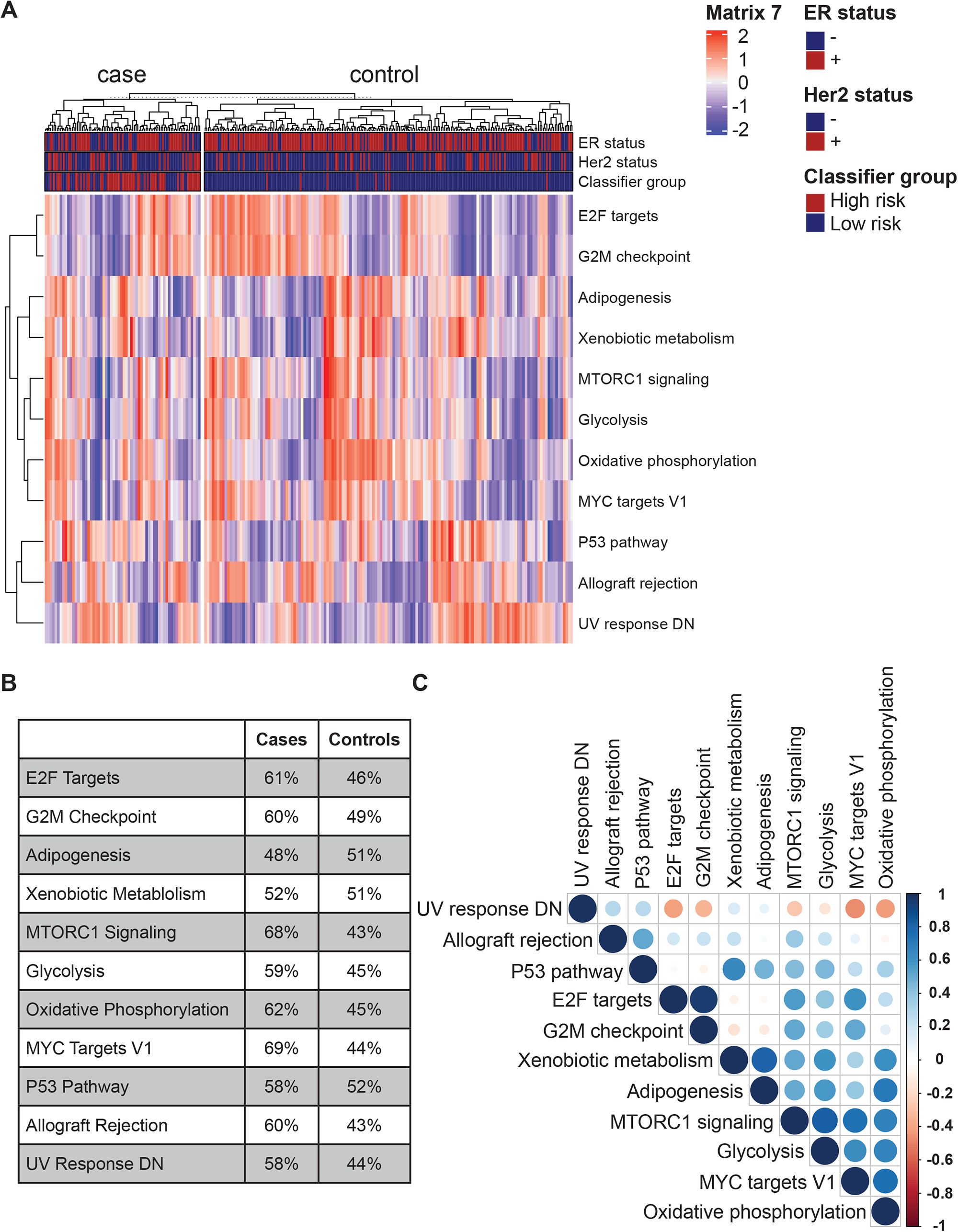
Outcome-associated pathways analyzed in individual samples. **A**) Heatmap of single sample Gene Set Variation Analysis of the 11 Hallmark pathways associated with recurrence. **B**) Percentage of samples in each 5-year outcome group enriched for each pathway i **A**). **C**) Plot showing the correlation between pathways in **A**). Positive correlations are displayed in blue and negative correlations in red. The color intensity and size of the circle are proportional to the correlation coefficients. Bar on the right: Correlation coefficients and corresponding colors.

IBC has been genomically profiled with several approaches, including the PAM50 and IC classification schemes. While DCIS and IBC are part of the same neoplastic process, there are differences in the TME, evolutionary age, and inter-observer variability in diagnostic labeling at different stages of progression. This suggests that a DCIS-specific classification scheme would correlate better with the biologic and clinical features of DCIS. Our analysis revealed clear deviations from the PAM50 subtypes, supporting the conclusion that they are not apt for DCIS characterization, as previously described (Bergholtz et al., 2020, Swanson et al., 2019). Here, we identified three transcriptomic subgroups of DCIS, characterized by ER signaling, proliferation and metabolism, that represent the fundamental genomic organization at this early stage of breast neoplasia. We found that this classification of DCIS more accurately captures the spectrum of DCIS biology than classification schemes derived from IBC, and can be robustly applied across cohorts and transcriptome protocols. The three subgroups identified here may represent the earliest variation in neoplasia transcriptome and may be applicable to earlier stages of neoplasia, such as hyperplasias.

There are several possible explanations for why traditional IBC classifiers do not perform well on DCIS. First, HER2 expression is more common at the DCIS stage than at the IBC stage (Allred et al., 1992), potentially because of its ability to inhibit anoikis (Whelan et al., 2013, Gupta et al., 2019). It is plausible that the more common HER2 expression in DCIS leads to a different transcriptomic distribution compared to IBCs. Many ER-DCIS express HER2 without amplification, in contrast to IBC, where the HER2-specific (amplified) subtype is clearer. Moreover, the cells in DCIS are confined to the epithelial compartment and interact with myoepithelial cells and the basement membrane, and thus are presumably restricted by rules of differentiation that govern normal epithelial cells. This regulation could constrain the transcriptomic variability of the neoplastic cells and in turn the resulting possible subtypes. Finally, the evolutionary age of the neoplasm may influence differences in classification of DCIS and IBC. By comparing WGS data from primary DCIS and IBCs, we found that the same constellation of copy number changes was present in both, consistent with previous studies (Ma et al., 2003, Vincent-Salomon et al., 2008b, Hwang et al., 2004). While DCIS had fewer genomic alterations than IBC, and a larger group of DCIS were classified as genomically quiescent tumors, recurrent genomic events that drive the IC scheme in IBC were evident as early as the DCIS stage.

A unique aspect of our study is the separate profiling of stroma and epithelial components through CSx analysis of LCM-derived stromal gene expression coupled with the in situ MIBI technique. We identified four stromal subtypes characterized by distinct gene pathways, stromal-, and immune cell composition. Moreover, specific stromal patterns were correlated with epithelial expression patterns, and particularly HER2+ and ER-DCIS were associated with a stronger inflammatory response. This could be associated with co-amplification of *ERBB2* (HER2) and a cluster of chemokine encoding genes on the 17q12 chromosomal region (Gil Del Alcazar et al., 2017).

The stroma plays a key role in cancer initiation and progression. In invasive breast cancer, transcriptional analysis of LCM derived samples has identified that stromal gene expression signatures are highly correlated with clinical outcome with features characteristic of angiogenesis, hypoxia, and tumor-associated macrophages (Finak et al., 2008). These early findings have been supported by numerous studies that have further characterized the role of immune surveillance in breast cancer outcome. While the IBC microenvironment has been extensively studied, that of pure DCIS and its role in cancer progression remain less understood. Due to a largely intact basement membrane and myoepithelial cell layer, most DCIS tumor cells are not as exposed to the immune environment as IBC cells (Gil Del Alcazar et al., 2017). Studies have shown that the DCIS TME is characterized by higher immune infiltration than normal breast (Hussein and Hassan, 2006, Gil Del Alcazar et al., 2020), and both tumor infiltrating lymphocytes and macrophage abundance have been associated with high grade DCIS and necrosis (Hendry et al., 2017, Campbell et al., 2017). The transition from DCIS to IBC is complex, with some noting a switch to a less active tumor immune environment (Gil Del Alcazar et al., 2017), and others supporting an increase in CD4+ and FOXP3+ TIL infiltration with invasive progression (Hwang et al., 2004, Kim et al., 2020). Some of these immune alterations appear to be subtype specific, with HER2+ and triple negative DCIS characterized by a higher frequency of cytotoxic CD* GZMB T cells compared with invasive cancer (Gil Del Alcazar et al., 2017). TIL infiltration in DCIS is associated with both activated and immune gene signatures even within the same sample, indicating a complex relationship between immune activation and immune surveillance in the progression from DCIS to invasive cancer (Trinh et al., 2021).

## TECHNICAL CONSIDERATIONS

Generating an atlas of DCIS is similar to the effort of TCGA for IBC. However, there are important differences and challenges in DCIS. First, obtaining DCIS tissue samples is considerably more challenging. In IBC, the tumor is evident by gross exam, and can be easily obtained as fresh, fresh frozen, or archival material. This is not the case for pre-invasive lesions. DCIS can sometimes be recognized radiographically but is only precisely detailed by pathologic examination, making prospective tissue collection a challenge. An additional complication in the study of DCIS is the ambiguous nature of the process. In IBC, the transition from an intraepithelial neoplasia to an invasive neoplasia is definitional. For DCIS, such a clear-cut definition does not exist. DCIS is broadly defined by significant cytologic and architectural changes compared to normal breast architecture by a growth of neoplastic cells in the inter-epithelial compartment.

## LIMITATIONS OF THE STUDY

The Breast Pre-Cancer Atlas presented here provides a foundational advancement in the study of precancerous lesions and will be a valuable resource for years to come. Its utility comes, in part, from the inclusion of two independent and large-scale cohorts that have important and distinct differences. For example, the two cohorts represent subjects from different geographical sites, eligibility criteria for inclusion, median years of diagnosis, and time to recurrence. There were no significant differences in age at diagnosis or treatment across cohorts. However, future observations on a cohort of patients with DCIS who are undergoing watchful waiting would provide outcome results that may be more aligned with emerging treatment strategies of DCIS.

One issue that should be noted is the genetic relationship between the primary DCIS and the subsequent ipsilateral cancer. While we did not specifically assume that every IBE was clonally related to the primary DCIS, identifying a recurrence signature from the DCIS would be more likely if this was true. Recent work (Lips et al., 2021) on a large cohort indicates that 18% of ipsilateral invasive events may be unrelated to the primary DCIS based on mutation and copy number alterations. Non-clonal recurrences were more likely to be in a different breast quadrant and have discordant ER expression whereas time to recurrence and patient age were not significantly associated. While we did not examine the recurrences in the current study to determine clonality, it is likely that a similar fraction would be identified as “unrelated”. We anticipate that further refinement and validation of our recurrence signature will be strengthened by eliminating non-clonal iBEs.

## CONCLUSION

We have identified a novel genomic classifier that predicts both recurrence and invasive progression, using one of the largest, comprehensively annotated case/control data sets of primary DCIS assembled to date. The classifier is comprised of both epithelial and stromal features. Our findings support that progression is a process that requires both invasive propensity among the DCIS cells and stromal permissiveness in the TME. We propose this classifier as the basis for a future clinical test to assess outcomes in patients with primary DCIS in order to guide therapy, such as aggressive treatment (resection, radiation, hormonal therapy) vs less aggressive treatment (watchful waiting or surgery alone) and plan for these data to be a shared resource for the research community.

## ACKNOWLEDGMENTS

R01 CA185138-01 (ESH); U2C CA-17-035 Pre-Cancer Atlas (PCA) Research Centers (ESH, RBW, CM, KP, GAC, KO); UO1 CA214183 (JRM); DOD BC132057 (ESH, CM); BCRF 19-074 (ESH); BCRF 19-028 (GAC) PRECISION CRUK Grand Challenge (JW); R01CA193694 (RBW, GAC), BCRF PPI-18-006 (RBW). AEI RYC2019-026576-I, “LaCaixa” Foundation LCF/PR/PR17/51120011 (JAS). SHS was supported by the Lundbeck Foundation (R288-2018-35) and the Danish Cancer Society (R229-A13616). KEH was supported by a CIHR Banting Postdoctoral Fellowship. TBCRC 038 was conducted by the TBCRC, which receives major funding support from The Breast Cancer Research Foundation and Susan G. Komen. Some results in this paper are based upon data generated by the TCGA Research Network: https://www.cancer.gov/tcga.

## AUTHOR CONTRIBUTIONS

Conceptualization: ESH, RBW, CM, GAC, JRM, CC, and KP. Investigation: SHS, BRG, KEH, JAS, TR, SVe, AK, LC, DJV, KD, DYD, DZ, JL, SJ, SS, AH, and RBW. Resources: LAS, TH, BH, FC, KG, MK, SW, AD, TK, PM, JN, JL, JTs, AMS, AT, GG, CZ, MM, JW, SXZ, and SVa, Data curation: LK and JTa; Writing – Original Draft: SHS, BRG, KEH, JRM, ESH, and RBW. Writing – Review & Editing: All co-authors. Funding Acquisition: ESH, RBW, GAC, and CM.

Project Administration: RB, LA, and CS. Supervision: CC, RT, RMA, KO, KP, CM, JRM, GAC, ESH, and RBW.

## Supplementary Figure Legends

**Figure S1: Outcome analysis. Supplemental to Figure 2**.

**A**) The 29 recurrent CNAs were not robustly associated with progression. Hazard ratios from CoxPH modeling. Vertical dotted line represents HR = 1. Covariate on the right indicates if CNA is gain (red) or loss (blue). **B-C**) Forest plot of multivariable Cox regression analysis including the 812 gene classifier (high- vs. low-risk groups), treatment, age, DCIS grade, and ER status by IHC for any iBE with full follow-up in TBCRC (**B**) and RAHBT (**C**). **D-E**) Kaplan-Meier plot of time to progression (any iBE, full follow-up time) stratified by clinical ER status (IHC based) in TBCRC (**D**) and RAHBT (**E**). P-values from log-rank tests. **F**) ROC curve of the 812 gene Random Forest classifier tested towards progression-free survival in TCGA IBC samples. **G**) Kaplan-Meier plot of time to progression in TCGA IBC samples. P-value from log-rank test. **H**) ROC curve of the 812 gene Random Forest classifier tested towards overall survival in TCGA IBC samples. **I**) Kaplan-Meier plot of time to death in TCGA IBC samples. P-value from log-rank test. **J**) GSEA Hallmark pathway analysis of the differentially expressed genes between cases with any iBE at 5 years after treatment vs the rest in TBCRC. Only significant pathways shown (adj.P<0.05). Pathways are sorted by effect size.

**Figure S2:LCM dissection of DCIS epithelium and associated stroma in RAHBT, and characterization of RNA Subtypes, supplemental to Figure 3**.

**A)** Marked DCIS epithelium (blue) prior to dissection. **B**) Dissected DCIS epithelium on cap. **C**) Remaining tissue on slide after LCM dissection of DCIS epithelium. **D**) Marked stroma (yellow) adjacent to dissected DCIS epithelium (blue, panel A-C) prior to dissection. **E**) Dissected stroma on cap. **F**) Remaining tissue on slide after LCM dissection of DCIS epithelium and adjacent stroma. All images were taken at 2X magnification. **G**) NMF diagnostics supported three clusters in DCIS. Scatterplots show cophenetic and silhouette values with increasing numbers of clusters in TBCRC and RAHBT. **H**) Three subtypes are highly concordant across cohorts. Mosaic plot shows concordance of de novo clustering in RAHBT *vs* clusters determined from centroids identified in TBCRC. Blue indicates an enrichment while red indicates a depletion. **I-J)** UMAP projection of DCIS transcriptome colored by *de novo* RNA clusters in TBCRC (**I**) and RAHBT (**J**). **K)** GSEA Hallmark pathway analysis of each cluster vs rest for TBCRC and RAHBT LCM in full. **L**) Invasive breast cancer intrinsic subtypes do not fit DCIS. Boxplot shows Spearman *ᑭ* of DCIS and IBC samples with PAM50 centroids. The median correlations (Spearman’s *ᑭ*) were significantly lower for basal-like, Luminal A, and Normal-like IBC samples (Basal-like median *ᑭ*_IBC_ = 0.75; median *ᑭ*_DCIS_= 0.38; P_Bonferroni_ = 8.01×10^-16^; Luminal A median *ᑭ*_IBC_ = 0.60; median *ᑭ*_DCIS_= 0.50; P_Bonferroni_ = 3.13×10^-3^, Normal-like median *ᑭ*_IBC_ = 0.60; median p_DCIS_= 0.49; P_Bonferroni_ = 6.21×10^-3^). P-values from Wilcoxon rank sum test. Dots are colored by PAM50 subtype and the covariate along the top indicates if the sample is IBC or DCIS. Boxplot represents median, 0.25 and 0.75 quantiles with whiskers at 1.5x interquartile range. **M-N)** UMAP projection of DCIS transcriptome in TBCRC (**M**) and RAHBT LCM (**N**) colored by PAM50 subtype. Large circles represent the PAM50 subtype centroids. **O-P**) PAM50 do not robustly predict progression in DCIS. Kaplan-Meier plots of time to progression in TBCRC (**O**) and RAHBT (**P**). **Q-R**) *ESR1* (ER) and *ERBB2* (HER2) mRNA abundance are highly correlated with protein levels of ER (**Q**) and HER2 (**R**), respectively, as measured by MIBI. **S**) *PGR* mRNA abundance was highest in ER_high_ cluster.

**Figure S3:Characterizing the CNA landscape of DCIS, supplemental to Figure 4**.

**A**) PGA is not correlated with the number of mapped reads. **B**) PGA (median dichotomized) is not associated with progression. **C**) Silhouette plot from NMF unsupervised clustering of the CNA landscape of DCIS in TBCRC and RAHBT combined. **D**) Consensus matrix from NMF unsupervised clustering of the CNA landscape of DCIS in TBCRC and RAHBT combined. **E**) DNA clusters are not biased by depth of sequencing. **F-G**) Kaplan-Meier plot of time to progression stratified by the eight CNA clusters in TBCRC (**F**) and RAHBT (**G**). **H**) GSEA Hallmark pathway analysis of DE genes by DNA cluster in matched RNA samples (each cluster vs rest) for TBCRC and RAHBT in full. Clusters with no significant pathway enrichment or depletion are not included.

**Figure S4:Analysis of the tumor microenvironment, supplemental to Figure 5**.

**A**) Signature matrix created with CSx, with 12 different immune cell types. **B**) Protein validation of CSx signature matrix by MIBI. Correlogram showing MIBI-estimated cell types vs CSx-estimated cell types in RAHBT LCM samples. **C**) GSEA Hallmark pathway analysis of DE genes in each stromal cluster vs rest in RAHBT LCM stromal samples in full. **D**) Percentage of fibroblasts, endothelial and total immune cells present in each stromal cluster estimated by CSx. **E**) Abundance of total myeloid, lymphoid and granulocyte cells, represented as percentage of total immune cells. **F-G**) Abundance of 12 immune cell types, represented as percentage of total immune cells, by stromal clusters. **H**) Cell neighborhood frequencies from MIBI by stromal clusters in RAHBT LCM. Box plots (**D-H**): Red: Normal-like. Blue: Collage rich. Green: Desmoplastic. Purple: Immune dense. *: adj.P <0.05; **: adj.P < 0.01; ***: adj.P<0.001; ****:adj.P < 0.0001; ns: adj.P >0.05. **I**) Kaplan-Meier plot of time to progression (any iBE, full follow-up time) stratified by stromal clusters in RAHBT LCM. **J**) CSx inferred cell type distribution between 812 gene classifier risk groups (TBCRC and RAHBT combined). Only cell types with significant differences (adj.P<0.05) are shown. **K**) CSx inferred cell type distribution between cases with IBC iBEs, DCIS iBEs, and controls (TBCRC and RAHBT combined). Only cell types with significant differences (adj.P<0.05) are shown. **J, K**) * adj.P < 0.05. ** adj.P ≤ 0.1. *** adj.P ≤ 0.001; ns: adj.P >0.05. mDC: myeloid dendritic cells. pDC: plasmacytoid dendritic cells. NK: Natural killer cells.

## STAR Methods

### 1. Cohort collection and sample acquisition

#### RAHBT Cohort

The Resource of Archival Breast Tissue (RAHBT) is a data/tissue resource established by Drs. Allred and Colditz in 2008 focused on premalignant or benign breast disease. Uniform coding of premalignant lesions assures greater consistency and use of research. Follow-up through hospital record linkages documents subsequent breast lesions including IBC. The entire study population includes women ages 18 and older with documented cases of premalignant breast disease (including carcinoma in situ). The study was approved by the Washington University in St. Louis Institutional Review Board (IRB ID #: 201707090).

Women were identified as eligible through seven primary sources: Washington University School of Medicine Departmental databases (Surgery, Radiation Oncology, Pathology, and Radiology), and the Siteman Oncology Services Database (local tumor registry), the St. Louis Breast Tissue Repository, and the Women’s Health Repository. We reviewed all records, excluded women with cancer prior to qualifying premalignant lesions and identified 1831 unique women with DCIS or DCIS and subsequent recurrence. A common data set with pathologic details, risk factor data, treatment, and unique identifiers was created and used to follow these women for subsequent breast lesions. Centralized pathology review confirmed 174 cases of DCIS with recurrent lesions. For each case (with subsequent ipsilateral or contralateral breast events) we matched two controls who remained free from subsequent breast events based on race, year of diagnosis (+/-5 years), age at diagnosis (+/-5 years), and type of definitive surgery (mastectomy or lumpectomy). For each DCIS diagnosis we retrieved slides and blocks for pathology review, secured a whole slide image of each sample, marked for TMA cores, and prepared for laboratory processing. A total of 172 cases and 338 controls were cored for TMAs. Breast pathology review was completed by Drs. Allred, Warrick, DeSchryver, and Veis.

To define an external validation data set that used identical eligibility criteria to TBCR 038 including year of initial DCIS diagnosis, we identified an additional set of cases from RAHBT and used comparable laboratory procedures for RNA-seq.

For RAHBT, 97 patients were analyzed by RNA-seq (**Table 1**). The median age at diagnosis was 53, and median year of diagnosis 2006. Time to recurrence with ipsilateral IBC was 36 months, and to diagnosis of ipsilateral DCIS 46.9 months. For women in the cohort with no iBEs, median follow up extended to 141 months. Treatment of initial DCIS ranged from lumpectomy with radiation (66.0%), and no radiation (10.3%) and mastectomy (23.7%). This subset of the RAHBT cohort was composed of 35.1% African American women.

For RAHBT LCM, 265 patients were analyzed by RNA-seq (**Table S1**). The median age at diagnosis was 53, and median year of diagnosis 2002. Time to recurrence with ipsilateral IBC was 80 months, and to diagnosis of ipsilateral DCIS 50 months. For women in the cohort with no iBEs, median follow up extended to 111 months. Treatment of initial DCIS ranged from lumpectomy with radiation (52%), and no radiation (18%) and mastectomy (28%). This subset of the RAHBT cohort was composed of 25% African American women.

#### TBCRC 038 Cohort

TBCRC 038 is a retrospective multi-center study activated at 12 participating TBCRC (Translational Breast Cancer Consortium) sites, which identified women treated for ductal carcinoma in situ (DCIS) at one of the enrolling institutions between 01/01/1998 and 02/29/2016. The TBCRC and the Department of Defense (DOD) approved this study for the collection of archival tissues. Duke served as the initiating and central site for all data, samples, assays, and analysis. The study was approved by the Duke Health Institutional Review Board (Protocol ID: Pro00068646) as well as the IRB at each participating institution. Individual sites reviewed medical records to identify patients eligible for the study.

Study eligibility criteria included: Women aged 40-75 years at diagnosis of DCIS without invasion; no prior treatment for breast cancer; and definitive surgical excision with no ink on tumor margins and treated with mastectomy, lumpectomy with radiation, or lumpectomy.

Cases (patients with subsequent iBEs) were matched 1:1 to controls with at least 5 years of follow up without subsequent iBEs. Matching was based on year of diagnosis (+/-5 years), age at diagnosis (+/-5 years), and DCIS nuclear grade (high grade vs. non-high grade). All cases consisted of initial diagnosis of pure DCIS, with ipsilateral recurrence occurring no less than 12 months from date of primary diagnosis. Clinical data, including treatment data, were collected at each site, and standardized data points were entered into a web-based portal. Tumor tissue was collected from FFPE blocks and cut into 5um sections. All slides were scanned and reviewed centrally by a breast pathologist (AH) to confirm the diagnosis. Tumor tissue marked by the pathologist was macrodissected for bulk analysis assays.

The 216 patients from the TBCRC cohort analyzed by RNA-seq (**Table 1**) includes 95 women without iBE after 5 or more years, 66 with DCIS iBEs, and 55 with IBC iBEs. Median time to IBC iBE for this subset was 58 months and 40 months to DCIS iBE. 30% of this subset were African American.

### 2. Wet lab methods

#### a. TMA construction

Qualified DCIS or subsequent lesion slides were assembled for pathology review. The research breast pathologist marked the slides for best area to core (1mm) for the carcinoma in situ and later event. The TMAs were designed such that cases/controls were assigned randomly on the map. The Beecher Tissue Arrayer was used to take a core from the patient donor block and place it in the designated area of the recipient TMA block. Slides were then cut for research purposes, and stained H&E and unstained slides were prepared. The TMAs were stored in the St. Louis Breast Tissue Registry Lab at room temperature.

#### b. Slide cutting

A TMA cutting breakdown was established to include slides for laser capture microdissection (LCM PEN membrane glass slides) sequencing, multiplex protein (MIBI high-purity gold-coated slides) staining and charged glass slides for FISH analysis of the RAHBT TMAs. The order of the slides for the different assays was as follows:

Slide 1-3: FISH/routine IHC – 4 um slices on charged slides

Slide 4-6: RNA/DNA sequencing – 7 um slices on LCM membrane glass slides

Slide 7: MIBI analysis – 4 um slices on gold coated slides

Slide 8-10: FISH/routine IHC – 4 um slices on charged slides

Slide 11-13: RNA/DNA sequencing – 7 um slices on LCM membrane slides

Slide 14: MIBI analysis – 4 um slices on gold coated slides

Slide 15-17: FISH/routine IHC – 4 um slices on charged slides

Slide 18 H&E stained.

#### c. Digital H&E generation (scanners)

At Washington University School of Medicine, the H&E original slide and TMA slide for RAHBT was imaged (20x) by Aperio AT2 (Leica). ImageScope provides the software for viewing the slides. Images are stored on secure servers in the Dept of Pathology, Washington University School of Medicine.

#### d. Pathologic analysis and masking

For the TBCRC cohort, whole slide images of the H&E slide made from the block sourced for DNA and RNA was reviewed and scored for grade, presence of necrosis and architecture by a breast pathologist (AH). For the RAHBT cohort, H&E images from the TMAs were used to score for grade, presence of necrosis and architecture by four breast pathologists (DJV, AH, SS, RBW). Areas of DCIS and normal tissue from the RAHBT TMAs were annotated and masked for LCM by two breast pathologists (SS and RBW).

#### e. LCM

Consecutive sections of tissue microarray blocks were cut and mounted on PEN membrane slides. Slides were dissected immediately after staining on an Arcturus XT LCM System based on the masked areas. Epithelial and stromal sections were dissected separately (**Figure S1**). Each sample adhere to a CapSure HS LCM Cap (Thermo Fisher #LCM0215). After LCM, the cap was sealed in an 0.5 mL tube (Thermo Fisher #N8010611) and stored at −80°C until library preparation. The matching epithelial regions in consecutive slides were dissected for corresponding DNA libraries.

#### f. smart-3seq

Sequencing libraries were prepared according to the Smart-3SEQ method (Foley et al., 2019) starting from dissected FFPE tissue on an Arcturus LCM HS Cap, except for the unique P5 index and universal P7 primers. Three control samples were added to each library preparation batch and sequence batch to allow batch effect analysis. Libraries were pooled together according to qPCR measurements and prepared according to the manufacturer’s instructions with a 1% spike-in of the PhiX control library (Illumina #FC-110-3002) and sequenced on an Illumina NextSeq 500 instrument with a High Output v2.5 reagent kit (Illumina # 20024906),

#### g. DNA-seq

Genomic DNA was isolated from LCM FFPE cells using PicoPure DNA Extraction kit (Thermo Fisher Scientific # KIT0103). 50ul lysis buffer with Proteinase K were added to each sample and incubated at 65°C overnight. After inactivating proteinase K, the genomic DNA was cleaned up with AMPure XP beads at 3:1 ratio (Beckman Coulter# A63880) and eluted in the 10mM Tris-HCl (pH8.0).

DNA Libraries were constructed with KAPA HyperPlus Kit (Kapa Biosystems #07962428001). Barcode adapters were used for multiplexed sequencing of libraries with SeqCap Adapter Kit A (Kapa Biosystems #7141530001). DNA libraries were amplified by 19 PCR cycles. AMPure XP beads were used for the size selection and cleaning up. DNA libraries were eluted in the 30 μL 10mM Tris-HCl (pH8.0).

Library size distribution was assessed on an Agilent 2100 Bioanalyzer using the DNA 1000 assay and the concentration was measured by Qubit® dsDNA HS Assay Kit (Thermo Fisher Scientific # Q32851). For each lane, 12 samples were pooled and sequenced by Novogene (Sacramento, CA, US) on the Illumina HiSeq Platform, collecting 110G per 275M reads output of paired-end reads of 150 bp length.

#### h. MIBI

For full details of the MIBI methods, see the companion paper by Risom et al. Briefly, antibodies were conjugated to isotopic metal reporters. Tissues were sectioned (5μm section thickness) from tissue blocks on gold and tantalum-sputtered microscope slides. Imaging was performed using a MIBI-TOF instrument with a Hyperion ion source.

### 3. Data processing

#### a. RNA-seq processing

RNA sequencing data was processed with 3SEQtools (https://github.com/jwfoley/3SEQtools). Single-end Illumina FASTQ files were generated from NextSeq BCL files with bcl2fastq (v2.20.0.422) and then aligned to reference hg38 with STAR aligner (v2.7.3a). Samples that did not meet a minimum threshold of uniquely aligned reads were filtered out. The samples in this study averaged 1.11 million uniquely aligned reads. Gene expression matrices of raw and normalized read counts were produced from BAM files with featureCounts (v1.6.4) of the Subread package (v2.4.2) and GENCODE Release 33.

#### b. DNA-seq processing

Low-pass WGS data were preprocessed using the Nextflow-base pipeline Sarek (Garcia et al., 2020) v2.6.1 with BWA v0.7.17 for sequence alignment to the reference genome GRCh38/hg38 and GATK (McKenna et al., 2010) v4.1.7.0 to mark duplicates and calibration. The recalibrated reads were further processed and filtered for mappability, GC content using the R/Bioconductor quantitative DNA-sequencing (QDNAseq) v1.22.0 with R v3.6.0. For QDNAseq, 50-kb bins were generated from (http://doi.org/10.5281/zenodo.4274556). We kept only autosomal sequences after filtering due to low-depth mappability and GC correction. We used the QDNAseq corrected output and segmented for CN analysis using the circular binary segmentation (CBS) algorithm from DNAcopy R/Bioconductor package v1.60.0. Copy number aberrations were called using CGHcall v2.48.0 (van de Wiel et al., 2007). The R/Bioconductor package ACE v1.4.0 (Poell et al., 2019) was used to estimate purity and ploidy. Proportion of the genome copy number altered (PGA) was calculated based on CNAs with |log_2_ ratio| > 0.3 based on the following:

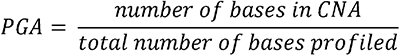

#### c. MIBI

Multiplexed image sets were extracted, slide background-subtracted, denoised, and aggregate filtered. Nuclear segmentation was performed using an adapted version of the DeepCell CNN architecture. Single cell data was extracted for all cell objects and area normalized. The FlowSOM R package v1.22.0 (Van Gassen et al., 2015) was used to assign each cell to one of five major cell lineages (tumor, myoepithelial, fibroblast, endothelial, immune). Immune cells were subclustered to delineate B cells, CD4+ T cells, CD8+ T cells, monocytes, MonoDC cells, DC cells, macrophages, neutrophils, mast cells, double-negative CD4−CD8−T cells, and HLADR+ APC cells. Tumor and fibroblast cells were similarly sub clustered to reveal phenotypic subsets. A total of 16 cell populations were quantified and analyzed. For full details of the MIBI methods, see the companion paper by Risom et al.

### 4. Analyses

#### a. ER, HER2 status

Clinical ER status (by IHC) was available for 83.3% (180 of 216) of the TBCRC cohort and 83.5% (81 of 97) of the RAHBT cohort.

Additionally, we called ER and HER2 positivity based on mRNA abundance levels of *ESR1* and *ERBB2*, respectively. We applied a Guassian mixture model with two components using the mclust R package (v5.4.7).

#### b. PAM50 and IC10

PAM50 subtypes were called using the genefu v2.22.1 (Gendoo et al., 2016) R package. We compared the PAM50 subtypes called by genefu against subtypes called adjusting for the expected proportion of ER+ samples, as implemented in (Bergholtz et al., 2020). We found both methods to be highly concordant (>96% concordance). We compared the correlation of DCIS and IBC samples to the PAM50 centroids within the genefu R package using Spearman’s correlation. We also compared the silhouette widths based on Euclidean distances of the PAM50 subtypes to the de novo DCIS subtypes using the cluster R package (v2.1.1). IC10 subtypes were called using the iC10 (v1.5) R package.

#### c. Differential abundance analyses

Differential gene expression analysis was performed using the R package DESeq2 v1.30.1 (Love et al., 2014) with default options. P-values were adjusted for multiple testing using the

Benjamini-Hochberg method. FDR<0.05 was considered significant for all DESeq2 analyses. Reads matrices were VST normalized for downstream analyses.

#### d. Unsupervised clustering: Non-negative matrix factorization

We identified RNA and CNA based clusters by non-negative matrix factorization using the NMF R package v0.23.0 (Brunet et al., 2004). Each NMF rank was run 30 times to evaluate cluster stability. We comprehensively evaluated 2-10 clusters for each data type and evaluated cluster fit by cophenetic and silhouette values. RNA clusters were first discovered in TBCRC and replicated in RAHBT. We evaluated replication by quantifying the concordance of de novo clusters identified in RAHBT *vs* clusters determined from centroids identified in TBCRC. CNA clusters were discovered in TBCRC and RAHBT jointly and compared against clusters identified in TBCRC and RAHBT individually to ensure robustness.

#### e. Identification of recurrent CNAs (GISTIC)

Recurrent CNAs were identified from purity-adjusted segment CNA calls from QDNASeq for 228 DCIS samples using GISTIC2 v2.0.23 (Mermel et al., 2011). GISTIC2 was run with the following parameters: -ta 0.3 -td 0.3 -qvt 0.05 -brlen 0.98 -conf 0.95 -armpeel 1 -res 0.01 -rx 0. To ensure CNAs were not biased by sequencing depth, recurrent CNAs significantly associated (FDR < 0.05) with the number of uniquely mapped reads were filtered out. Associations were quantified by Mann-Whitney test. The number of uniquely mapped reads was determined from samtools flagstat (v1.9).

#### f. CIBERSORTx

Using single-cell RNA-seq datasets, a breast specific signature matrix was built to resolve proportions of tumor, fibroblasts, endothelial and immune cells from bulk RNA-seq data (Azizi et al., 2018). scRNAseq data was downloaded from Gene Expression Omnibus database (GEO data repository accession numbers GSE114727, GSE114725). Normalized counts were obtained using Seurat R package (v3.2.0). The resultant signature matrix contained 3484 genes and allowed to resolve different immune cell types, including B, CD8 T, CD4 T, NKT, NK, mast cells, neutrophils, monocytes, macrophages and dendritic cells. The signature matrix was first *in-silico* validated. In order to test the accuracy of the signature matrix, a set of samples (1/10 of each type) from the same scRNAseq dataset was reserved to build a synthetic matrix of bulk RNA-seq data. By mixing different proportions of single cell transcripts, the synthetic bulk was used to predict cell type proportions and subsequently correlated with the true proportions used to build the synthetic mix. Pearson’s coefficient was >0.75 in all the cases, and most >0.9 (**Figure S5C**). The aforementioned matrix was used to deconvolve the LCM RNA-seq samples and to compare CSx-estimated cell abundance with MIBI-identified cell types. Cell abundance between groups was compared by Wilcoxon test followed by Benjamini-Hochberg correction for multiple testing.

#### g. Shared Nearest Neighbor clustering

LCM stromal samples from RAHBT were classified using the Shared Nearest Neighbor (SNN) clustering method implemented in the Seurat R package (v3.2.0). Data was normalized by negative binomial regression (sctransform R package, v0.3.2, variable.feature.n = “all.genes”). The first 15 principal components were used to identify the clusters and 16 different resolutions were compared, selecting resolution 0.75 and four clusters as the final solution. Positive markers were selected at a minimum fraction of 0.25 and the resultant gene list was used to further characterize each cluster by gene ontology and KEGG pathway analysis, implemented in clusterProfiler R package (version 3.18.1).

#### h. Statistical analyses

We used Mann-Whitney U test to compare continuous distributions between two groups, as specified in the text. We used the Kruskal-Wallis test to compare continuous values between three groups. All statistical analyses were implemented in the R statistical language (v3.6.1). P-values were corrected for multiple hypothesis testing *via* Bonferroni (when <10 independent tests) or Benjamini & Hochberg (when >10 independent tests).

#### i. Pathway & Gene Set Enrichment Analyses

Gene set enrichment analyses were performed using fgsea R package (v1.12.0) based on the MSigDB Hallmark pathways v7.4, (Subramanian et al., 2005). All genes from differential abundance analyses were included, and were ranked by their signed adjusted P-values. Pathways were considered enriched if adjusted P-values < 0.05. We evaluated pathway concordance across the DCIS subtypes using a hypergeometric test.

Single sample gene set variation analysis was performed using the GSVA R package (Hanzelmann et al., 2013) (v1.38.2) using default parameters.

#### j. Data visualization

Boxplots, heatmaps, scatterplots and barplots were generated using the BoutrosLab.plotting.general R package v6.0.3 (P’ng et al., 2019), or the R packages ggplot2 (v3.3.3, boxplots), corrplot (v0.84, scatterplots), and ComplexHeatmap (v.2.6.2, heatmaps). UMAPs were generated using the umap (v0.2.7.0) R package with the number of genes indicated in the text. Mosaic plots were generated using the vcd (v1.4.8) R package.

#### k. Outcome analysis

Associations with time to event were quantified using Cox Proportional Hazard model correcting for treatment as indicated in the text. To standardize follow-up across TBCRC and RAHBT, we censored the follow-up time at 250 months, the maximum follow-up time in TBCRC. Kaplan-Meier plots as implemented in the R packages survival (v3.2.10) and survminer (v0.4.9) were used to visualize outcome differences.

The 812 gene classifier was built using the cforest implementation of Random Forest in the Caret (v6.0-91) R package using default parameters. The TBRCR cohort was used as the training cohort and the model was tested on the RAHBT cohort. Hyperparameters were tuned on the training cohort using four-fold cross validation. The mtry parameters 5, 20, 50, 100, 200, 500, and 800 were tested and the optimal mtry selected was 5. Accuracy of the classifier was assessed using ROC curve, Precision, Recall, and F1 score.

Breast cancer data (BRCA) from TCGA was downloaded from https://www.cancer.gov/tcga. A total of 1064 samples with available follow-up information was used to test the 812 gene classifier towards progression-free survival and overall survival as defined in the TCGA-BRCA metadata.

RNA for the TCGA samples was normalized using the same protocols as the DCIS RNA-sequencing (TBCRC and RAHBT cohorts, above). The accuracy of the classifier in the TCGA cohort was assessed using ROC curve, Precision, Recall, and F1 score.

## 5. Data and Code Availability

All custom code used to analyze data will be made available through a Github repository.

The datasets generated during this study will be made available on the Human Tumor Atlas Network public repository.

De-identified images, including whole slide images of the H&E slide made from the block sourced for DNA and RNA, will be available on the Human Tumor Atlas Network public repository.

Further information and requests for resources and reagents should be directed to and will be fulfilled by the Lead Contact, Robert B West.

