## Supplemental information for "DCIS genomic signatures define biology and clinical outcome: Human Tumor Atlas Network (HTAN) analysis of TBCRC 038 and RAHBT cohorts"

### Figure S1

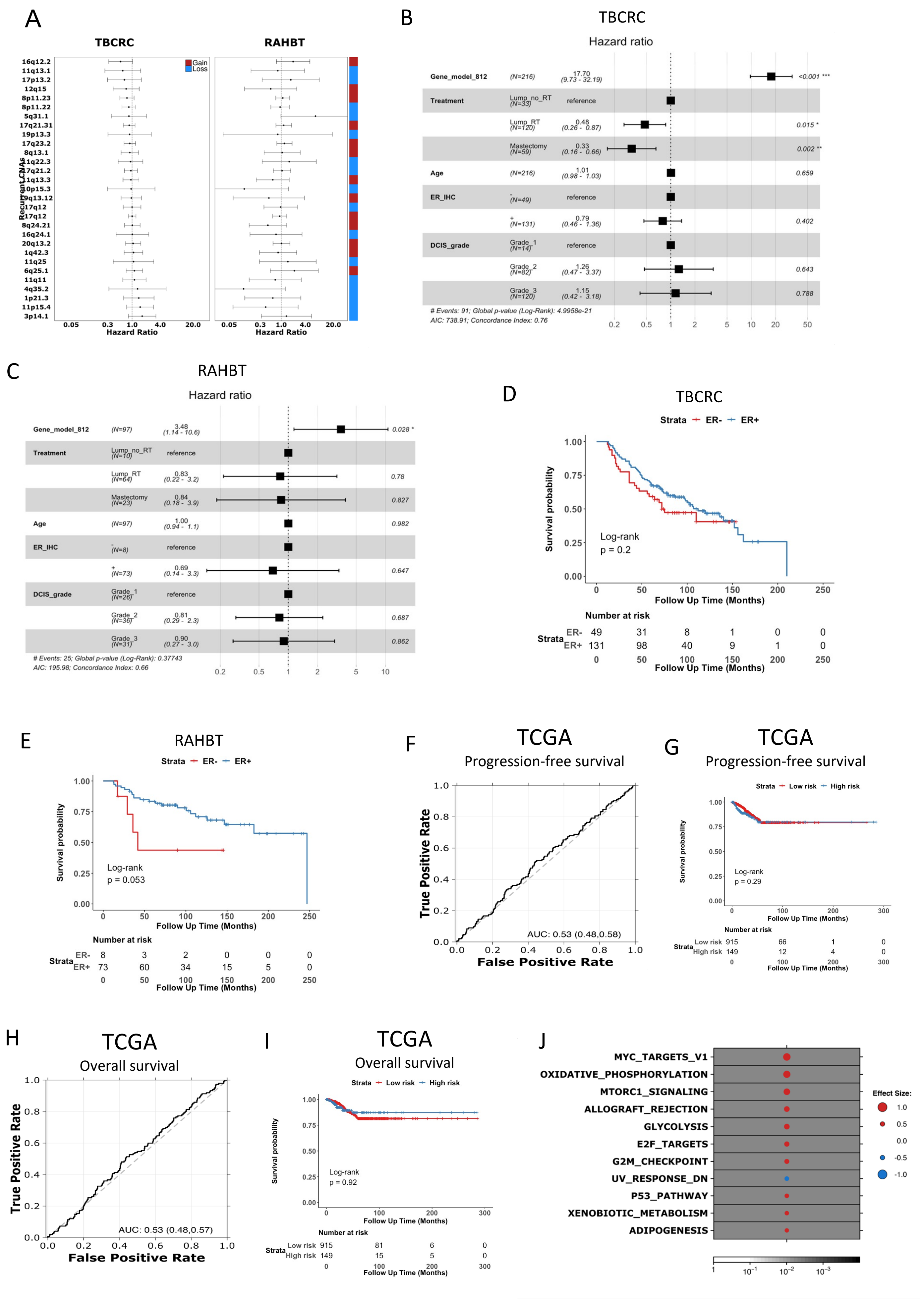

A
TBCRC
RAHBT
Gain
Loss
16q12.2
11q13.1
17p13.2
12q15
8p11.23
8p11.22
5q31.1
17q21.31
19p13.3
17q23.2
8q13.1
1q22.3
17q21.2
11q13.3
10p15.3
19q13.12
17q12
17q12
8q24.21
16q24.1
20q13.2
1q42.3
11q25
6q25.1
11q11
4q35.2
1p21.3
11p15.4
3p14.1
Recurrent CNAs
0.05 0.3 1.0 4.0 20.0
Hazard Ratio
B
TBCRC
Hazard ratio
Gene_model_812 (N=216) 17.70 (9.73 - 32.19) <0.001 ***
Treatment Lump_no_RT (N=33) reference
Lump_RT (N=120) 0.48 (0.26 - 0.87) 0.015 *
Mastectomy (N=59) 0.33 (0.16 - 0.66) 0.002 **
Age (N=216) 1.01 (0.98 - 1.03) 0.659
ER_IHC - (N=49) reference
+ (N=131) 0.79 (0.46 - 1.36) 0.402
DCIS_grade Grade_1 (N=14) reference
Grade_2 (N=82) 1.26 (0.47 - 3.37) 0.643
Grade_3 (N=120) 1.15 (0.42 - 3.18) 0.788
### Events: 91; Global p-value (Log-Rank): 4.9958e-21
AIC: 738.91; Concordance Index: 0.76
0.2 0.5 1 2 5 10 20 50
C
RAHBT
Hazard ratio
Gene_model_812 (N=97) 3.48 (1.14 - 10.6) 0.028 *
Treatment Lump_no_RT (N=10) reference
Lump_RT (N=64) 0.83 (0.22 - 3.2) 0.78
Mastectomy (N=23) 0.84 (0.18 - 3.9) 0.827
Age (N=97) 1.00 (0.94 - 1.1) 0.982
ER_IHC - (N=8) reference
+ (N=73) 0.69 (0.14 - 3.3) 0.647
DCIS_grade Grade_1 (N=26) reference
Grade_2 (N=36) 0.81 (0.29 - 2.3) 0.687
Grade_3 (N=31) 0.90 (0.27 - 3.0) 0.862
### Events: 25; Global p-value (Log-Rank): 0.37743
AIC: 195.98; Concordance Index: 0.66
0.2 0.5 1 2 5 10
D
TBCRC
Strata ER- ER+
Survival probability
Log-rank
p = 0.2
Follow Up Time (Months)
Number at risk
Strata ER- 49 31 8 1 0 0
ER+ 131 98 40 9 1 0
0 50 100 150 200 250
E
RAHBT
Strata ER- ER+
Survival probability
Log-rank
p = 0.053
Follow Up Time (Months)
Number at risk
Strata ER- 8 3 2 0 0 0
ER+ 73 60 34 15 5 0
0 50 100 150 200 250
F
TCGA
Progression-free survival
True Positive Rate
False Positive Rate
AUC: 0.53 (0.48,0.58)
G
TCGA
Progression-free survival
Strata Low risk High risk
Survival probability
Log-rank
p = 0.29
Follow Up Time (Months)
Number at risk
Strata Low risk 915 66 1 0
High risk 149 12 4 0
0 100 200 300
H
TCGA
Overall survival
True Positive Rate
False Positive Rate
AUC: 0.53 (0.48,0.57)
I
TCGA
Overall survival
Strata Low risk High risk
Survival probability
Log-rank
p = 0.92
Follow Up Time (Months)
Number at risk
Strata Low risk 915 81 6 0
High risk 149 15 5 0
0 100 200 300
J
MYC_TARGETS_V1
OXIDATIVE_PHOSPHORYLATION
MTORC1_SIGNALING
ALLOGRAFT_REJECTION
GLYCOLYSIS
E2F_TARGETS
G2M_CHECKPOINT
UV_RESPONSE_DN
P53_PATHWAY
XENOBIOTIC_METABOLISM
ADIPOGENESIS
Effect Size:
1.0
0.5
0.0
-0.5
-1.0
1 10^-1 10^-2 10^-3

### Figure S2

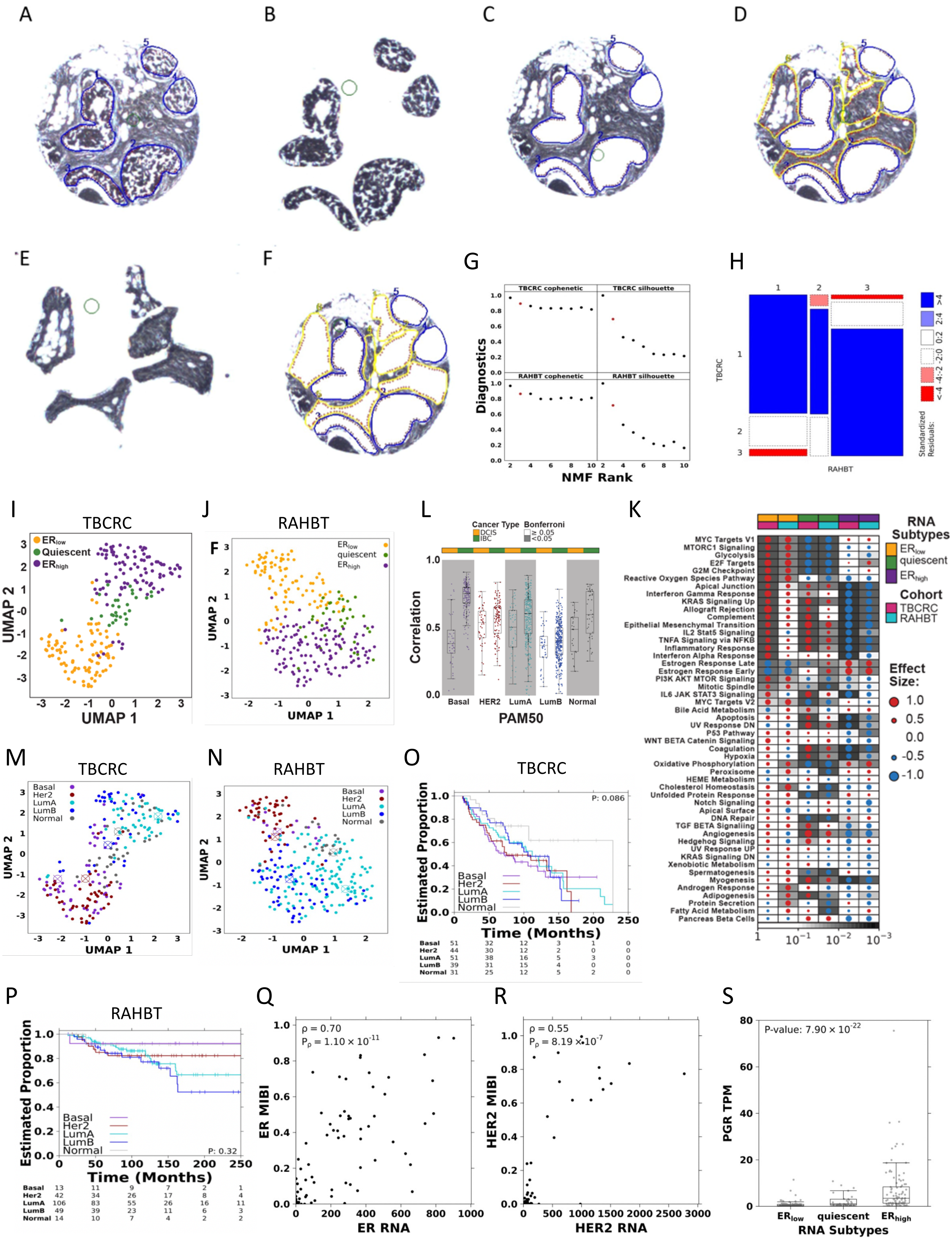

A
B
C
D
E
F
G
TBCRC cophenetic
TBCRC silhouette
RAHBT cophenetic
RAHBT silhouette
Diagnostics
NMF Rank
H
TBCRC
RAHBT
Standardized Residuals:
<-4 -4:-2 -2:0 0:2 2:4 >4
I
TBCRC
$ER_{low}$
Quiescent
$ER_{high}$
UMAP 2
UMAP 1
J
RAHBT
$ER_{low}$
quiescent
$ER_{high}$
UMAP 2
UMAP 1
L
Cancer Type
DCIS
IBC
Bonferroni
≥ 0.05
<0.05
Correlation
Basal
HER2
LumA
LumB
Normal
PAM50
K
RNA Subtypes
$ER_{low}$
quiescent
$ER_{high}$
Cohort
TBCRC
RAHBT
Effect Size:
1.0
0.5
0.0
-0.5
-1.0
MYC Targets V1
MTORC1 Signaling
Glycolysis
E2F Targets
G2M Checkpoint
Reactive Oxygen Species Pathway
Apical Junction
Interferon Gamma Response
KRAS Signaling Up
Allograft Rejection
Complement
Epithelial Mesenchymal Transition
IL2 Stat5 Signaling
TNFA Signaling via NFKB
Inflammatory Response
Interferon Alpha Response
Estrogen Response Late
Estrogen Response Early
PI3K AKT MTOR Signaling
Mitotic Spindle
IL6 JAK STAT3 Signaling
MYC Targets V2
Bile Acid Metabolism
Apoptosis
UV Response DN
P53 Pathway
WNT BETA Catenin Signaling
Coagulation
Hypoxia
Oxidative Phosphorylation
Peroxisome
HEME Metabolism
Cholesterol Homeostasis
Unfolded Protein Response
Notch Signaling
Apical Surface
DNA Repair
TGF BETA Signaling
Angiogenesis
Hedgehog Signaling
UV Response UP
KRAS Signaling DN
Xenobiotic Metabolism
Spermatogenesis
Myogenesis
Androgen Response
Adipogenesis
Protein Secretion
Fatty Acid Metabolism
Pancreas Beta Cells
1
$10^{-1}$
$10^{-2}$
$10^{-3}$
M
TBCRC
Basal
Her2
LumA
LumB
Normal
UMAP 2
UMAP 1
N
RAHBT
Basal
Her2
LumA
LumB
Normal
UMAP 2
UMAP 1
O
TBCRC
P: 0.086
Estimated Proportion
Basal
Her2
LumA
LumB
Normal
Time (Months)
Basal 51 32 12 3 1 0
Her2 44 30 12 2 0 0
LumA 51 38 16 5 3 0
LumB 39 31 15 4 0 0
Normal 31 25 12 5 2 0
P
RAHBT
Estimated Proportion
Basal
Her2
LumA
LumB
Normal
P: 0.32
Time (Months)
Basal 13 11 9 7 2 1
Her2 42 34 26 17 8 4
LumA 106 83 55 26 16 11
LumB 49 39 23 11 6 3
Normal 14 10 7 4 2 2
Q
$\rho$ = 0.70
$P_{\rho}$ = 1.10 × $10^{-11}$
ER MIBI
ER RNA
R
$\rho$ = 0.55
$P_{\rho}$ = 8.19 × $10^{-7}$
HER2 MIBI
HER2 RNA
S
P-value: 7.90 × $10^{-22}$
PGR TPM
$ER_{low}$
quiescent
$ER_{high}$
RNA Subtypes

### Figure S3

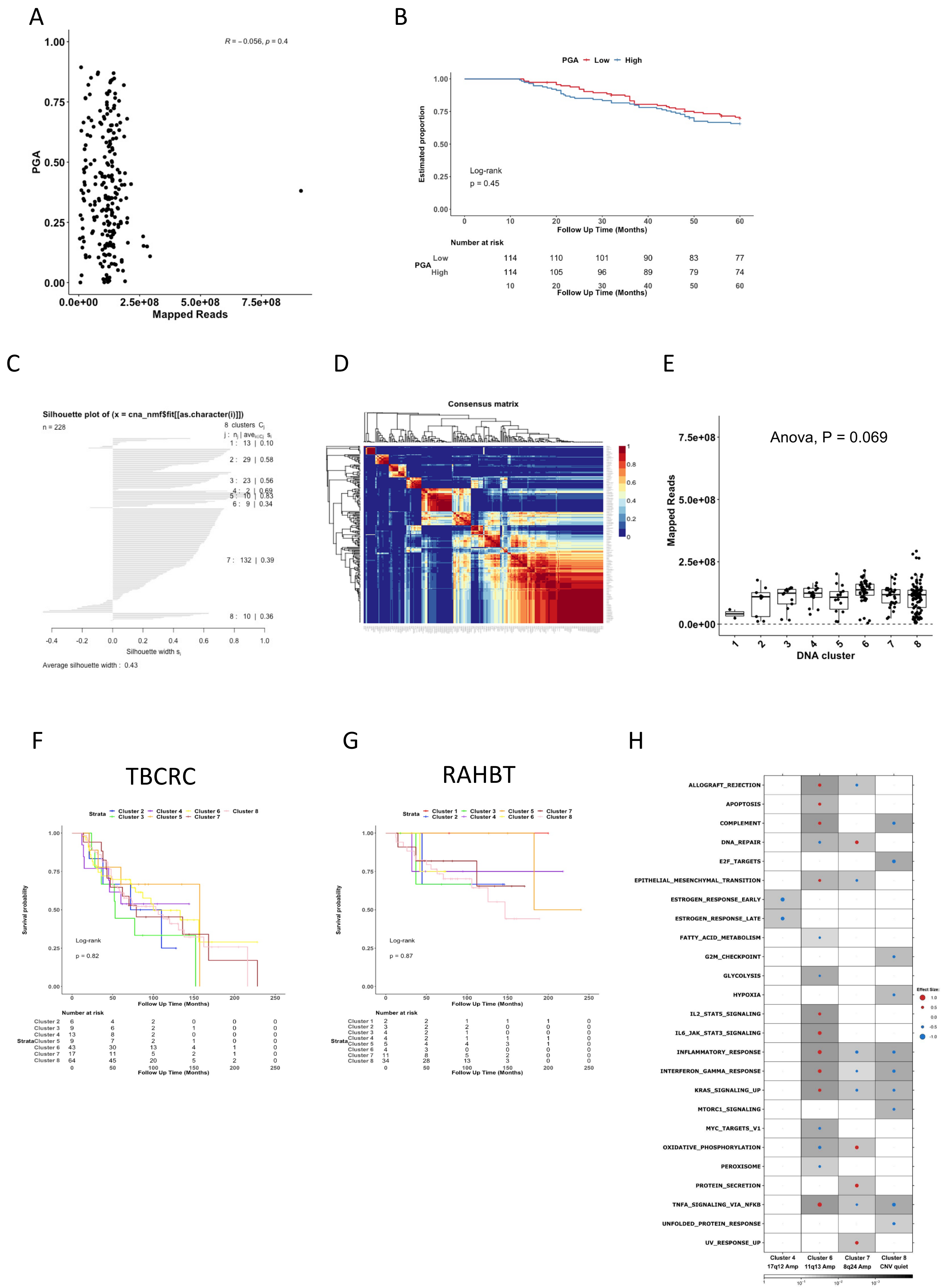

A
B
C
D
E
F
G
H
R = −0.056, p = 0.4
PGA
Mapped Reads
PGA Low High
Estimated proportion
Log-rank
p = 0.45
Follow Up Time (Months)
Number at risk
Silhouette plot of (x = cna_nmf$fit[[as.character(i)]])
n = 228
Silhouette width
Average silhouette width : 0.43
Consensus matrix
Anova, P = 0.069
DNA cluster
TBCRC
RAHBT
Survival probability
Log-rank
p = 0.82
p = 0.87
ALLOGRAFT_REJECTION
APOPTOSIS
COMPLEMENT
DNA_REPAIR
E2F_TARGETS
EPITHELIAL_MESENCHYMAL_TRANSITION
ESTROGEN_RESPONSE_EARLY
ESTROGEN_RESPONSE_LATE
FATTY_ACID_METABOLISM
G2M_CHECKPOINT
GLYCOLYSIS
HYPOXIA
IL2_STAT5_SIGNALING
IL6_JAK_STAT3_SIGNALING
INFLAMMATORY_RESPONSE
INTERFERON_GAMMA_RESPONSE
KRAS_SIGNALING_UP
MTORC1_SIGNALING
MYC_TARGETS_V1
OXIDATIVE_PHOSPHORYLATION
PEROXISOME
PROTEIN_SECRETION
TNFA_SIGNALING_VIA_NFKB
UNFOLDED_PROTEIN_RESPONSE
UV_RESPONSE_UP
Cluster 4 17q12 Amp
Cluster 6 11q13 Amp
Cluster 7 8q24 Amp
Cluster 8 CNV quiet

### Figure S4

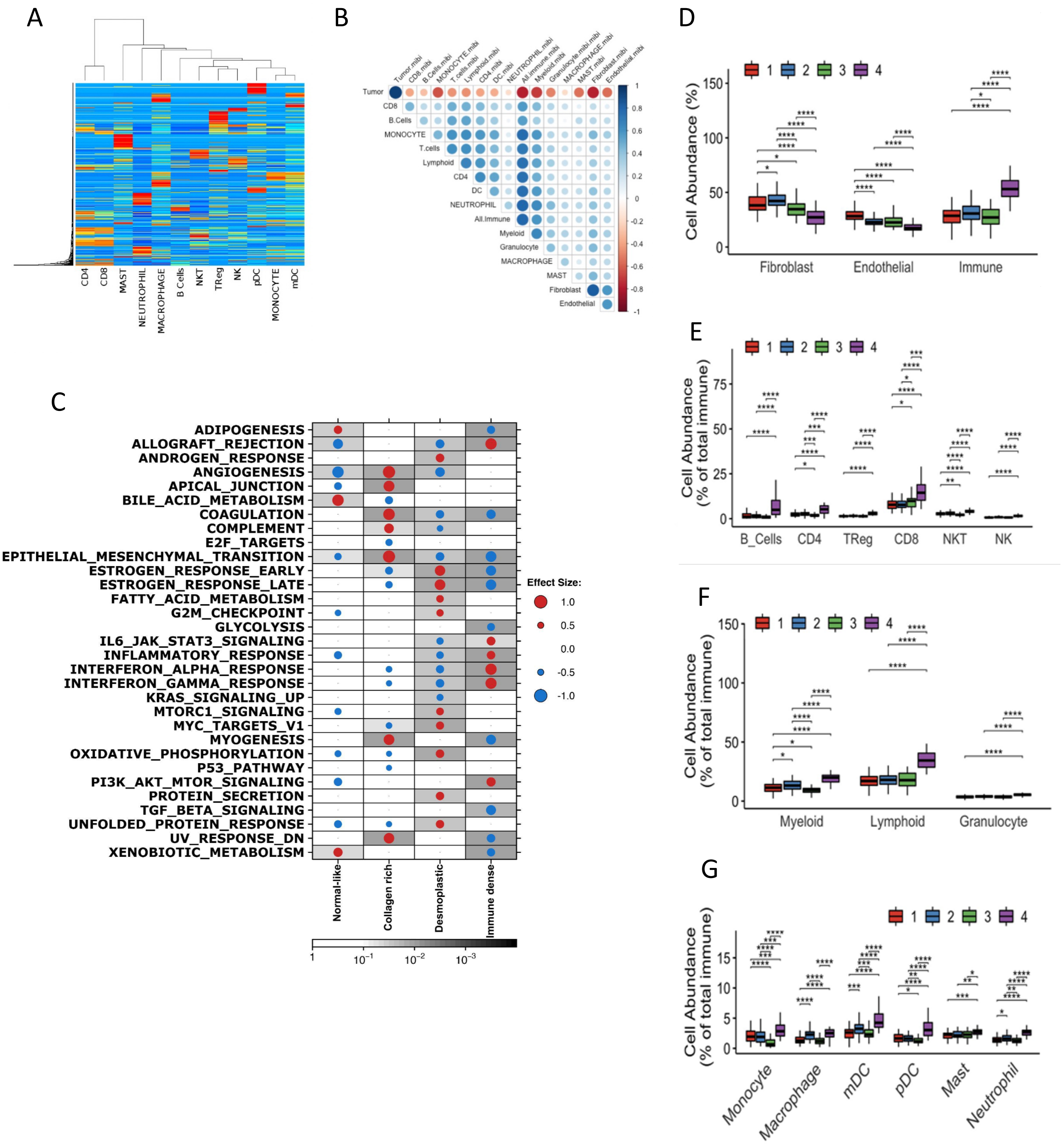
A
CD4
CD8
MAST
NEUTROPHIL
MACROPHAGE
B Cells
NKT
TReg
NK
pDC
MONOCYTE
mDC
B
Tumor
CD8
B.Cells
MONOCYTE
T.cells
Lymphoid
CD4
DC
NEUTROPHIL
All.Immune
Myeloid
Granulocyte
MACROPHAGE
MAST
Fibroblast
Endothelial
C
ADIPOGENESIS
ALLOGRAFT_REJECTION
ANDROGEN_RESPONSE
ANGIOGENESIS
APICAL_JUNCTION
BILE_ACID_METABOLISM
COAGULATION
COMPLEMENT
E2F_TARGETS
EPITHELIAL_MESENCHYMAL_TRANSITION
ESTROGEN_RESPONSE_EARLY
ESTROGEN_RESPONSE_LATE
FATTY_ACID_METABOLISM
G2M_CHECKPOINT
GLYCOLYSIS
IL6_JAK_STAT3_SIGNALING
INFLAMMATORY_RESPONSE
INTERFERON_ALPHA_RESPONSE
INTERFERON_GAMMA_RESPONSE
KRAS_SIGNALING_UP
MTORC1_SIGNALING
MYC_TARGETS_V1
MYOGENESIS
OXIDATIVE_PHOSPHORYLATION
P53_PATHWAY
PI3K_AKT_MTOR_SIGNALING
PROTEIN_SECRETION
TGF_BETA_SIGNALING
UNFOLDED_PROTEIN_RESPONSE
UV_RESPONSE_DN
XENOBIOTIC_METABOLISM
Normal-like
Collagen rich
Desmoplastic
Immune dense
Effect Size:
1.0
0.5
0.0
-0.5
-1.0
D
Cell Abundance (%)
Fibroblast
Endothelial
Immune
E
Cell Abundance (% of total immune)
B_Cells
CD4
TReg
CD8
NKT
NK
F
Cell Abundance (% of total immune)
Myeloid
Lymphoid
Granulocyte
G
Cell Abundance (% of total immune)
Monocyte
Macrophage
mDC
pDC
Mast
Neutrophil

### Figure S4, cont.

H

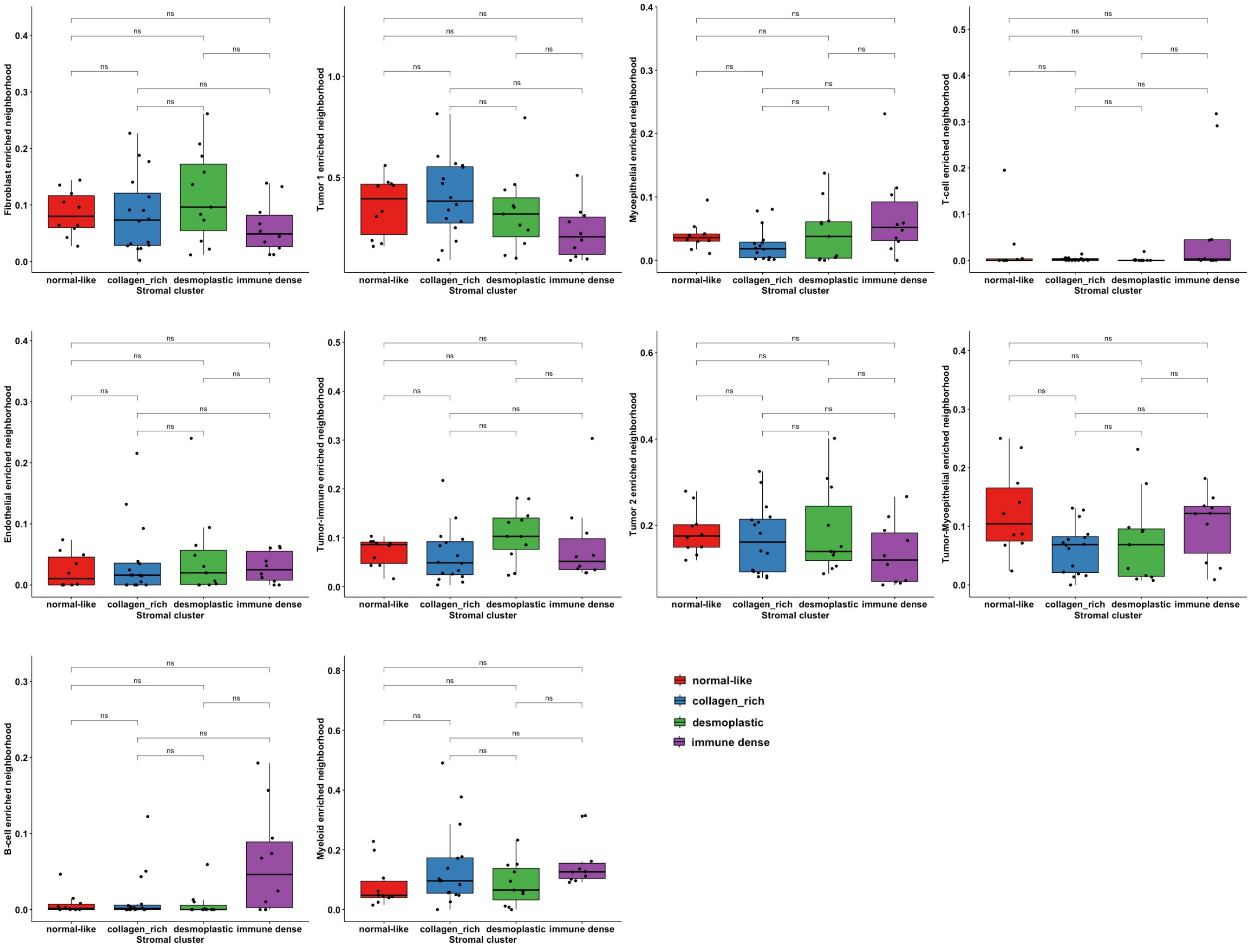

I

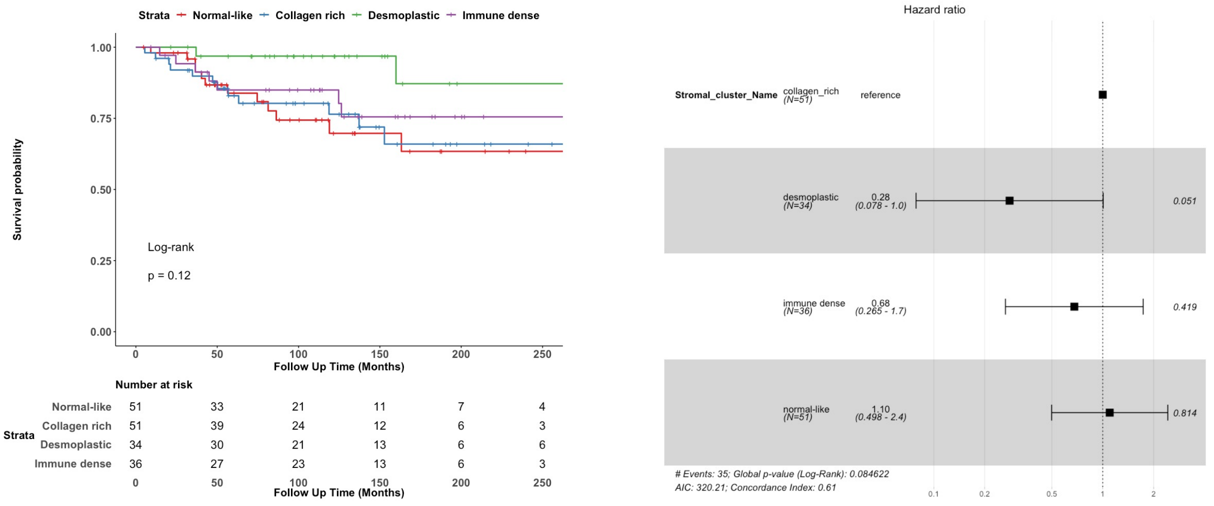

### Figure S4, cont.

J

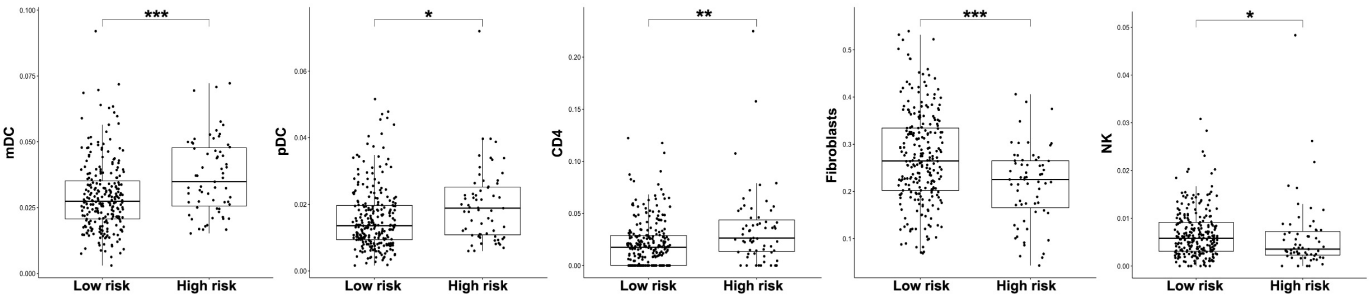

K

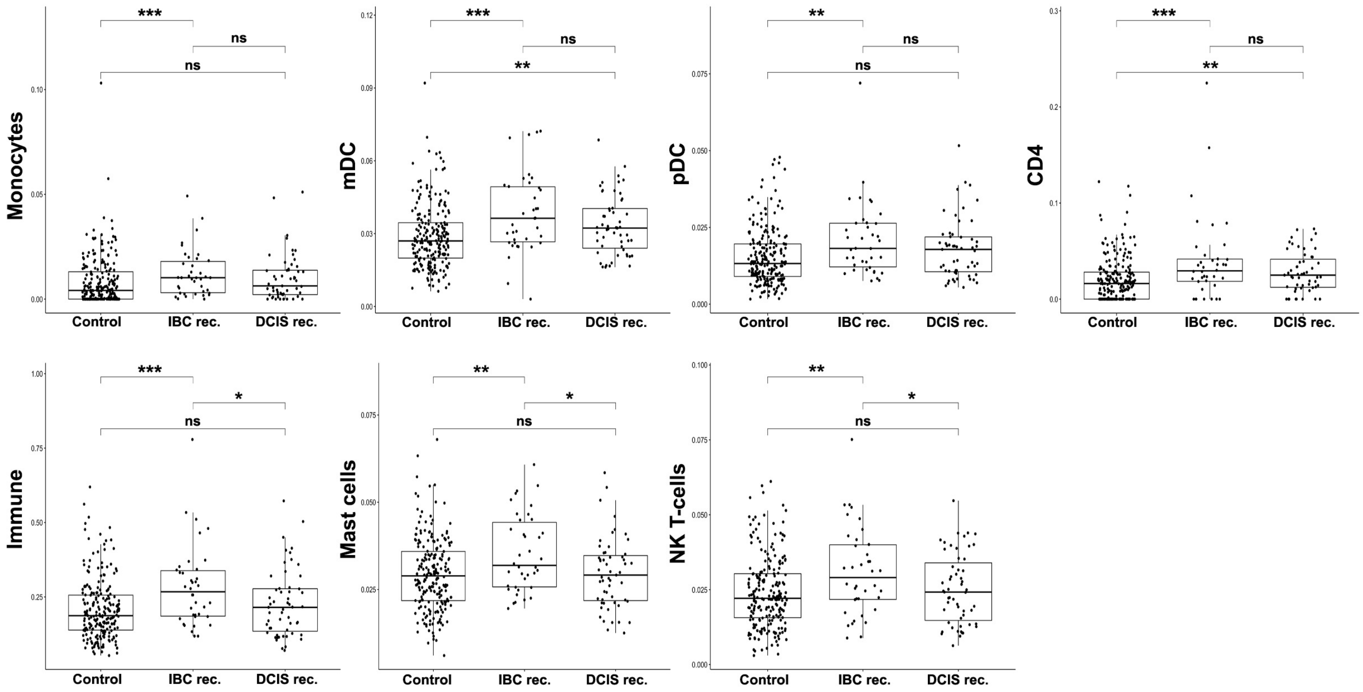

**Table S1:** Breast Pre-cancer Atlas Retrospective RAHBT Cohort for LCM. Supplemental to Figure 1 and Table 1. *To end of follow-up for no recurrence

| | RAHBT | | | | | |
|---|---|---|---|---|---|---|
| | **DCIS without recurrence (N=184)** | **DCIS with Ipsilateral DCIS Recurrence (N=17)** | **DCIS with Ipsilateral Invasive Recurrence (N=29)** | **DCIS with Contralateral DCIS (N=19)** | **DCIS with Contralateral Invasive Disease (N=16)** | **RAHBT Total (N=265)** |
| Year of Diagnosis | | | | | | |
| Median | 2002 | 2005 | 2000 | 2002 | 1991 | 2002 |
| Age at Diagnosis | | | | | | |
| Median | 53 | 57 | 48 | 57 | 54 | 53 |
| Mean (±SD) | 55.6 (±11.4) | 61.1 (±12.8) | 49.9 (±10.3) | 55.9 (±9.9) | 58.2 (±12.2) | 55.5 (±11.5) |
| Grade | | | | | | |
| 1 | 51 [27.7%] | 3 [17.6%] | 8 [27.6%] | 7 [36.8%] | 4 [25.0%] | 73 [27.5%] |
| 2 | 65 [35.3%] | 7 [41.2%] | 15 [51.7%] | 8 [42.1%] | 7 [43.8%] | 102 [38.5%] |
| 3 | 65 [35.3%] | 6 [35.3%] | 4 [13.8%] | 4 [21.1%] | 5 [31.3%] | 84 [31.7%] |
| Missing | 2 [1.1%] | 1 [5.9%] | 2 [6.9%] | 0 | 0 | 6 [2.3%] |
| **Pathologic Tumor Size** | | | | | | |
| Median | NA | NA | NA | NA | NA | NA |
| Mean (±SD) | NA | NA | NA | NA | NA | NA |
| **Marker Status** | | | | | | |
| ER(+) | 123 [66.8%] | 11 [64.7%] | 24 [82.8%] | 17 [89.5%] | 14 [87.5%] | 189 [71.3%] |
| ER(-) | 61 [33.2%] | 6 [35.3%] | 5 [17.2%] | 2 [10.5%] | 2 [12.5%] | 76 [28.7%] |
| ER(+) Dx before 2000 | 46 [25.0%] | 2 [11.8%] | 10 [34.5%] | 7 [36.8%] | 9 [56.2%] | 74 [27.9%] |
| ER(+) Dx 2000 & after | 77 [41.8%]] | 9 [52.9%] | 14 [48.3%] | 10 [52.6%] | 5 [31.2%] | 67 [25.3%] |
| ER(-) Dx before 2000 | 29 [15.8% | 3 [17.6%] | 4 [13.8%] | 2 [10.5%] | 1 [6.3%] | 87 [32.8%] |
| ER(-) Dx 2000 & after | 32 [17.4%] | 3 [17.6%] | 1 [3.4%] | 0 | 1 [6.3%] | 37 [14.0%] |
| **Treatment** | | | | | | |
| Lumpectomy w Radiation | 91 [49.5%] | 12 [70.6%] | 18 [62.1%] | 8 [42.1%] | 9 [50.0%] | 17 [51.7%] |
| Lumpectomy no Radiation | 34 [18.5%] | 5 [29.4%] | 7 [24.1%] | 1 [5.3%] | 0 | 47 [17.7%] |
| Lumpectomy Radiation Unknown | 3 [1.6%] | 0 | 1 [3.4%] | 1 [5.3%] | 1 [6.3%] | 6 [2.3%] |
| Mastectomy | 56 [30.4%] | 0 | 3 [10.3%] | 9 [47.4%] | 7 [43.8%] | 75 [28.3%] |
| **Time to Recurrence* (months)** | | | | | | |

| | | | | | | |
|---|---|---|---|---|---|---|
| Median | 111* | 49 | 80 | 81 | 56 | 62.3 |
| Mean (±SD) | 127.1 (±84.4) | 61.5 (±43.6) | 93.2 (±74.2) | 107.3 (±89.1) | 71.3 (±56.3) | 85.5 (±70.6) |
| **Margins** | | | | | | |
| Ink on tumor | 9 [4.9%] | 2 [11.8%] | 2 [6.9%] | 3 [15.8%] | 2 [12.5%] | 17 [6.4%] |
| <2mm | 24 [13.0%] | 3 [17.6%] | 3 [10.3%] | 3 [15.8%] | 3 [18.8%] | 36 [13.6%] |
| At least 2mm | 27 [14.7%] | 4 23.5%] | 2 [6.9%] | 1 [5.3%] | 1 [6.3%] | 38 [14.3%] |
| Clear, unknown mm | 81 [44.0%] | 8 [47.1%] | 17 [58.6%] | 10 [52.6%] | 4 [25.0%] | 118 [44.5%] |
| Missing | 43 [23.4%] | 0 | 5 [17.2%] | 2 [10.5%] | 6 [37.5%] | 56 [21.1%] |
| **Race** | | | | | | |
| White | 138 [75.0%] | 12 [70.6%] | 22 [75.9%] | 15 [78.9%] | 10 [62.5%] | 197 [74.3%] |
| Black | 45 [24.5%] | 5 [29.4%] | 7 [24.1%] | 3 [15.8%] | 6 [37.5%] | 66 [24.9%] |
| Asian | 0 | 0 | 0 | 0 | 0 | 0 |
| Pacific Islander | 0 | 0 | 0 | 1 [5.3%] | 0 | 1 [0.4%] |
| Other | 0 | 0 | 0 | 0 | 0 | 0 |
| Unknown | 1 [0.5%] | 0 | 0 | 0 | 0 | 1 [0.4%] |
| Black | 45 [24.5%] | 5 [29.4%] | 7 [24.1%] | 3 [15.8%] | 6 [37.5%] | 66 [24.9%] |
| Asian | 0 | 0 | 0 | 0 | 0 | 0 |
| Pacific Islander | 0 | 0 | 0 | 1 [5.3%] | 0 | 1 [0.4%] |
| Other | 0 | 0 | 0 | 0 | 0 | 0 |
| Unknown | 1 [0.5%] | 0 | 0 | 0 | 0 | 1 [0.4%] |

**Table S2:** Breast Pre-cancer Atlas Multi-scale Characterization Assays. Supplemental to Figure 1.

| **Assay** | **Scale** | **Type of Data** | **Integration and validation with other assays** | **Analyzed in RAHBT** | **Analyzed in RAHBT LCM** | **Analyzed in TBCRC** |
|---|---|---|---|---|---|---|
| RNA-seq (Single duct, tumor microenvironment) | Duct, organ, normal tissue | Whole transcriptome gene expression profiling per single duct | Gene expression and prediction of cell type composition (CibersortX) confirmed by MIBI (single cell) | Figure 2B, C, E, G<br>Figure 4F<br>Figure 6 | Figure 3B, C<br>Figure 5A-C, E, G | Figure 2A, D, F<br>Figure 3A, C<br>Figure 4F<br>Figure 6 |
| Low-pass whole genome DNA-seq | Duct and adjacent normal | CNV profiling per single duct | Analysis of CNV supported by RNA-seq (single duct) and MIBI (single cell) | Figure 4A-E | NA | Figure 4A-E |
| Multiplex IHC (MIBI) | Cell | 1. Cell type<br>2. Proteomic analysis | Analysis of protein expression and cell type supported by RNA-seq of ducts (CibersortX) | NA | Figure 3D, E, F<br>Figure 5D, F | NA |

**Table S3:** 812 differentially expressed genes from DESeq2 analysis iBEs within 5 years vs. the rest in TBCRC. Supplemental to Figure 2. Log2FoldChange > 0 : Up in iBEs within 5 years. Compartment column indicates if the respective gene was significantly differentially expressed (FDR<0.05) in the epithelial or stromal compartment by DESeq2 analysis of stromal vs epithelial RAHBT LCM samples.

| | baseMean | log2FoldChange | lfcSE | stat | P-value | FDR | Compartment |
|---|---|---|---|---|---|---|---|
| FRS2 | 166.7259 | -0.9103 | 0.1562 | 5.8289 | 0.0000 | 0.0001 | Epithelial |
| SLC30A2 | 9.5025 | -1.3093 | 0.2365 | 5.5371 | 0.0000 | 0.0002 | NA |
| ARPC4 | 115.8311 | 0.3632 | 0.0660 | -5.5070 | 0.0000 | 0.0002 | Stromal |
| RPS10 | 186.6479 | 0.4468 | 0.0795 | -5.6194 | 0.0000 | 0.0002 | NA |
| POLE3 | 103.3779 | 0.2992 | 0.0547 | -5.4733 | 0.0000 | 0.0002 | Epithelial |
| ZYG11B | 153.6384 | -0.2592 | 0.0487 | 5.3191 | 0.0000 | 0.0002 | Stromal |
| LEPR | 81.7666 | -0.7825 | 0.1457 | 5.3701 | 0.0000 | 0.0002 | Stromal |
| SULT1C2P1 | 2.9400 | 2.8289 | 0.5364 | -5.2740 | 0.0000 | 0.0002 | NA |
| SCN7A | 15.5456 | -0.8815 | 0.1678 | 5.2521 | 0.0000 | 0.0002 | Stromal |
| SDCBPP2 | 9.6769 | -1.5592 | 0.2964 | 5.2599 | 0.0000 | 0.0002 | NA |
| MRPL45 | 52.6197 | 0.8739 | 0.1636 | -5.3418 | 0.0000 | 0.0002 | Epithelial |
| MTCO1P40 | 73.0610 | 0.7268 | 0.1380 | -5.2654 | 0.0000 | 0.0002 | Epithelial |
| MT-CO2 | 611.4592 | 0.8407 | 0.1593 | -5.2771 | 0.0000 | 0.0002 | Epithelial |
| TXN | 324.1005 | 0.3545 | 0.0678 | -5.2266 | 0.0000 | 0.0002 | Epithelial |
| AP002360.2 | 9.1223 | 0.6803 | 0.1310 | -5.1918 | 0.0000 | 0.0003 | NA |
| PPP1R14B-AS1 | 7.2024 | 1.1226 | 0.2171 | -5.1719 | 0.0000 | 0.0003 | Epithelial |
| NAA10 | 62.5264 | 0.3525 | 0.0682 | -5.1652 | 0.0000 | 0.0003 | NA |
| STUM | 10.8517 | -1.2684 | 0.2487 | 5.1001 | 0.0000 | 0.0003 | NA |
| BRK1 | 160.7369 | 0.3812 | 0.0746 | -5.1116 | 0.0000 | 0.0003 | NA |
| TPH1 | 10.7104 | -1.2124 | 0.2374 | 5.1069 | 0.0000 | 0.0003 | NA |
| RPL19 | 1759.1388 | 0.8692 | 0.1710 | -5.0833 | 0.0000 | 0.0003 | Epithelial |
| RPL24 | 576.3087 | 0.3157 | 0.0624 | -5.0620 | 0.0000 | 0.0004 | Stromal |
| PTMA | 476.3741 | 0.5529 | 0.1100 | -5.0277 | 0.0000 | 0.0004 | NA |
| EIF3K | 221.1603 | 0.2932 | 0.0583 | -5.0279 | 0.0000 | 0.0004 | NA |
| RPS2 | 311.2376 | 0.4876 | 0.0971 | -5.0191 | 0.0000 | 0.0004 | Epithelial |
| S100A11 | 614.9017 | 0.4828 | 0.0966 | -4.9995 | 0.0000 | 0.0004 | Epithelial |
| MT-ATP6 | 3469.4586 | 0.5933 | 0.1197 | -4.9581 | 0.0000 | 0.0005 | Epithelial |
| FMO4 | 11.5632 | 0.9049 | 0.1831 | -4.9426 | 0.0000 | 0.0005 | NA |

| | | | | | | | |
|---|---|---|---|---|---|---|---|
| VDAC1 | 80.7150 | 0.4241 | 0.0860 | -4.9294 | 0.0000 | 0.0006 | Epithelial |
| CYP4Z1 | 45.0136 | 1.8857 | 0.3836 | -4.9160 | 0.0000 | 0.0006 | Epithelial |
| HOXC4 | 36.0919 | -0.5941 | 0.1209 | 4.9149 | 0.0000 | 0.0006 | NA |
| SET | 220.6596 | 0.3299 | 0.0674 | -4.8920 | 0.0000 | 0.0006 | Epithelial |
| LINC02611 | 18.2469 | 1.3154 | 0.2697 | -4.8766 | 0.0000 | 0.0006 | NA |
| COX5A | 43.4693 | 0.5103 | 0.1048 | -4.8688 | 0.0000 | 0.0006 | NA |
| RPS19 | 463.8981 | 0.5656 | 0.1163 | -4.8639 | 0.0000 | 0.0006 | Stromal |
| SNORA35B | 14.1749 | -0.8234 | 0.1692 | 4.8674 | 0.0000 | 0.0006 | Stromal |
| GALNT5 | 48.9741 | -1.1435 | 0.2362 | 4.8409 | 0.0000 | 0.0007 | Epithelial |
| RN7SL151P | 16.7452 | -0.7361 | 0.1529 | 4.8142 | 0.0000 | 0.0008 | NA |
| STK38L | 53.9741 | 0.4852 | 0.1010 | -4.8027 | 0.0000 | 0.0008 | Stromal |
| TFPI2 | 58.0207 | -1.5729 | 0.3282 | 4.7926 | 0.0000 | 0.0008 | Epithelial |
| DPAGT1 | 48.0511 | 0.3700 | 0.0775 | -4.7731 | 0.0000 | 0.0009 | Epithelial |
| TDRD12 | 4.8818 | -2.0909 | 0.4385 | 4.7678 | 0.0000 | 0.0009 | NA |
| RPS7 | 72.5643 | 0.6730 | 0.1420 | -4.7386 | 0.0000 | 0.0009 | NA |
| FANCM | 28.3416 | -0.4328 | 0.0912 | 4.7444 | 0.0000 | 0.0009 | NA |
| TK1 | 42.7004 | 0.7441 | 0.1568 | -4.7457 | 0.0000 | 0.0009 | Epithelial |
| UBE3AP2 | 9.0407 | -1.2024 | 0.2538 | 4.7383 | 0.0000 | 0.0009 | Stromal |
| GLYATL2 | 40.2562 | 1.9428 | 0.4118 | -4.7178 | 0.0000 | 0.0010 | Epithelial |
| RPL3 | 112.8046 | 0.7246 | 0.1546 | -4.6863 | 0.0000 | 0.0011 | NA |
| SMDT1 | 71.6589 | 0.3735 | 0.0796 | -4.6893 | 0.0000 | 0.0011 | Epithelial |
| RPS27 | 406.7587 | 0.6804 | 0.1456 | -4.6726 | 0.0000 | 0.0012 | NA |
| RPL13A | 583.5606 | 0.4922 | 0.1055 | -4.6643 | 0.0000 | 0.0012 | NA |
| NUTF2 | 80.6522 | 0.4020 | 0.0866 | -4.6403 | 0.0000 | 0.0013 | Epithelial |
| HDGF | 537.1875 | 0.2749 | 0.0594 | -4.6248 | 0.0000 | 0.0014 | Epithelial |
| ALG10B | 30.6788 | -0.4327 | 0.0936 | 4.6229 | 0.0000 | 0.0014 | NA |
| CHGA | 4.2989 | -2.4899 | 0.5401 | 4.6099 | 0.0000 | 0.0014 | NA |
| TAGLN2 | 667.8300 | 0.3986 | 0.0866 | -4.6038 | 0.0000 | 0.0015 | Epithelial |
| RPL7A | 418.3559 | 0.4178 | 0.0910 | -4.5933 | 0.0000 | 0.0015 | NA |
| RPL18 | 1325.6953 | 0.2930 | 0.0638 | -4.5955 | 0.0000 | 0.0015 | NA |
| ENO1 | 584.1607 | 0.4004 | 0.0874 | -4.5810 | 0.0000 | 0.0015 | Epithelial |
| S100A7 | 317.6505 | 2.5077 | 0.5472 | -4.5824 | 0.0000 | 0.0015 | Epithelial |

| | | | | | | | |
|---|---|---|---|---|---|---|---|
| LIFR | 114.6394 | -0.4420 | 0.0964 | 4.5846 | 0.0000 | 0.0015 | Stromal |
| SNORA79B | 23.0955 | -0.5493 | 0.1205 | 4.5570 | 0.0000 | 0.0017 | NA |
| RPS27A | 424.8563 | 0.5321 | 0.1169 | -4.5535 | 0.0000 | 0.0017 | Stromal |
| RPL23 | 1848.7490 | 0.7410 | 0.1629 | -4.5480 | 0.0000 | 0.0017 | Epithelial |
| ATP5MG | 189.8699 | 0.3173 | 0.0699 | -4.5394 | 0.0000 | 0.0017 | NA |
| KANSL1 | 200.9551 | -0.2638 | 0.0583 | 4.5265 | 0.0000 | 0.0018 | Stromal |
| MT-CYB | 3096.6212 | 0.4939 | 0.1092 | -4.5220 | 0.0000 | 0.0018 | Epithelial |
| ST13 | 194.8926 | 0.3250 | 0.0720 | -4.5102 | 0.0000 | 0.0019 | NA |
| C1orf116 | 18.6089 | 0.7786 | 0.1735 | -4.4880 | 0.0000 | 0.0020 | Epithelial |
| PSMD7 | 161.3428 | 0.3342 | 0.0745 | -4.4893 | 0.0000 | 0.0020 | NA |
| RPL35A | 890.5315 | 0.3082 | 0.0690 | -4.4694 | 0.0000 | 0.0022 | NA |
| TTC28 | 51.4364 | -0.3625 | 0.0812 | 4.4662 | 0.0000 | 0.0022 | Stromal |
| DNPH1 | 50.7834 | 0.4114 | 0.0924 | -4.4547 | 0.0000 | 0.0022 | Epithelial |
| RBM20 | 27.8790 | -1.6059 | 0.3601 | 4.4590 | 0.0000 | 0.0022 | Epithelial |
| RPL4 | 161.8864 | 0.3800 | 0.0853 | -4.4538 | 0.0000 | 0.0022 | Stromal |
| ABCC13 | 4.0090 | -1.7983 | 0.4039 | 4.4528 | 0.0000 | 0.0022 | Epithelial |
| TOMM40 | 46.6127 | 0.3369 | 0.0757 | -4.4483 | 0.0000 | 0.0022 | Epithelial |
| NDUFB11 | 99.0407 | 0.2781 | 0.0626 | -4.4455 | 0.0000 | 0.0022 | NA |
| PGK1 | 273.8511 | 0.3304 | 0.0744 | -4.4433 | 0.0000 | 0.0022 | Epithelial |
| TNRC6A | 328.2238 | -0.2043 | 0.0461 | 4.4337 | 0.0000 | 0.0023 | NA |
| RPL18A | 28.9285 | 0.8002 | 0.1808 | -4.4252 | 0.0000 | 0.0023 | NA |
| NDUFS5 | 145.9924 | 0.3530 | 0.0799 | -4.4203 | 0.0000 | 0.0024 | Epithelial |
| JPT1 | 126.1400 | 0.5764 | 0.1307 | -4.4096 | 0.0000 | 0.0025 | Epithelial |
| IGSF10 | 28.8993 | -0.6280 | 0.1426 | 4.4040 | 0.0000 | 0.0025 | Stromal |
| PHB2 | 183.9273 | 0.3159 | 0.0717 | -4.4032 | 0.0000 | 0.0025 | Epithelial |
| PLP1 | 7.8462 | -0.9068 | 0.2061 | 4.3994 | 0.0000 | 0.0025 | Stromal |
| CPLANE1 | 193.1142 | -0.3243 | 0.0738 | 4.3946 | 0.0000 | 0.0025 | Epithelial |
| FO393411.1 | 10.9132 | 0.8037 | 0.1833 | -4.3850 | 0.0000 | 0.0026 | NA |
| RPL32 | 1407.4865 | 0.2923 | 0.0667 | -4.3813 | 0.0000 | 0.0026 | Stromal |
| COX4I1 | 269.2329 | 0.3598 | 0.0825 | -4.3584 | 0.0000 | 0.0029 | NA |
| NCL | 877.1905 | 0.2058 | 0.0473 | -4.3545 | 0.0000 | 0.0029 | NA |
| KYNU | 37.1913 | 0.9441 | 0.2174 | -4.3427 | 0.0000 | 0.0029 | NA |

| | | | | | | | |
|---|---|---|---|---|---|---|---|
| MRPL51 | 124.4353 | 0.3170 | 0.0729 | -4.3487 | 0.0000 | 0.0029 | Epithelial |
| GAPDH | 1354.1132 | 0.3685 | 0.0849 | -4.3406 | 0.0000 | 0.0029 | Epithelial |
| LIPG | 5.1049 | 0.9250 | 0.2131 | -4.3398 | 0.0000 | 0.0029 | Stromal |
| SCGB1B2P | 9.5136 | -1.6117 | 0.3710 | 4.3440 | 0.0000 | 0.0029 | NA |
| C19orf48 | 31.5983 | 0.4550 | 0.1049 | -4.3383 | 0.0000 | 0.0029 | Epithelial |
| MT-CO1 | 633.6585 | 0.4873 | 0.1123 | -4.3381 | 0.0000 | 0.0029 | Epithelial |
| AL078622.1 | 3.8600 | 1.4241 | 0.3285 | -4.3348 | 0.0000 | 0.0029 | NA |
| EEF1B2 | 309.8869 | 0.3042 | 0.0702 | -4.3316 | 0.0000 | 0.0029 | Stromal |
| FAU | 535.6733 | 0.3132 | 0.0725 | -4.3221 | 0.0000 | 0.0030 | Stromal |
| MYT1L | 3.0995 | -1.2573 | 0.2921 | 4.3042 | 0.0000 | 0.0032 | NA |
| SNU13 | 142.9673 | 0.2887 | 0.0670 | -4.3058 | 0.0000 | 0.0032 | NA |
| AC104984.4 | 52.8122 | -0.6241 | 0.1451 | 4.3006 | 0.0000 | 0.0032 | NA |
| RPS29 | 1416.6224 | 0.3674 | 0.0858 | -4.2838 | 0.0000 | 0.0034 | Stromal |
| RPLP1 | 2700.8877 | 0.3998 | 0.0933 | -4.2847 | 0.0000 | 0.0034 | NA |
| MTRNR2L8 | 17.5107 | -0.8186 | 0.1914 | 4.2768 | 0.0000 | 0.0035 | Epithelial |
| CNOT2 | 184.0274 | -0.2344 | 0.0551 | 4.2574 | 0.0000 | 0.0038 | NA |
| CD55 | 77.2275 | 0.4205 | 0.0989 | -4.2501 | 0.0000 | 0.0038 | NA |
| NDRG1 | 220.3773 | 0.5430 | 0.1277 | -4.2511 | 0.0000 | 0.0038 | Stromal |
| MMP1 | 2.6952 | 1.5527 | 0.3651 | -4.2533 | 0.0000 | 0.0038 | Stromal |
| RPL35 | 1143.5910 | 0.2425 | 0.0571 | -4.2452 | 0.0000 | 0.0038 | NA |
| RPS12 | 1223.2005 | 0.4056 | 0.0956 | -4.2424 | 0.0000 | 0.0039 | Stromal |
| S100P | 92.1684 | 1.3833 | 0.3266 | -4.2358 | 0.0000 | 0.0039 | Epithelial |
| ISG20 | 39.9208 | 0.5928 | 0.1400 | -4.2352 | 0.0000 | 0.0039 | Stromal |
| RPS16 | 858.0049 | 0.3284 | 0.0775 | -4.2373 | 0.0000 | 0.0039 | Stromal |
| FDPS | 148.0351 | 0.3522 | 0.0833 | -4.2264 | 0.0000 | 0.0040 | Epithelial |
| GALE | 29.8475 | 0.4863 | 0.1155 | -4.2109 | 0.0000 | 0.0042 | Epithelial |
| TOGARAM1 | 76.6170 | -0.2760 | 0.0655 | 4.2128 | 0.0000 | 0.0042 | NA |
| NBR1 | 176.1716 | -0.2829 | 0.0672 | 4.2108 | 0.0000 | 0.0042 | NA |
| HMGA1 | 157.8932 | 0.4291 | 0.1020 | -4.2065 | 0.0000 | 0.0042 | Epithelial |
| NUP107 | 78.2503 | -0.3125 | 0.0745 | 4.1958 | 0.0000 | 0.0044 | NA |
| NHLRC2 | 78.6647 | -0.2195 | 0.0525 | 4.1767 | 0.0000 | 0.0047 | NA |
| RPL38 | 1193.5428 | 0.3215 | 0.0770 | -4.1748 | 0.0000 | 0.0047 | NA |

| | | | | | | | |
|---|---|---|---|---|---|---|---|
| PI3 | 6.6804 | 1.6641 | 0.3991 | -4.1700 | 0.0000 | 0.0048 | NA |
| MT-ND2 | 3821.4932 | 0.4804 | 0.1153 | -4.1677 | 0.0000 | 0.0048 | Epithelial |
| MRPS21 | 131.4244 | 0.3357 | 0.0807 | -4.1618 | 0.0000 | 0.0049 | Epithelial |
| NONO | 288.4391 | 0.1728 | 0.0415 | -4.1594 | 0.0000 | 0.0049 | Epithelial |
| EXTL2 | 50.1677 | -0.2809 | 0.0678 | 4.1426 | 0.0000 | 0.0052 | Stromal |
| IGFBP6 | 101.2148 | -0.5104 | 0.1233 | 4.1399 | 0.0000 | 0.0052 | Stromal |
| MRPS11 | 57.5934 | 0.3343 | 0.0808 | -4.1373 | 0.0000 | 0.0052 | NA |
| ABCA10 | 53.2498 | -0.6209 | 0.1501 | 4.1366 | 0.0000 | 0.0052 | Stromal |
| SMIM26 | 23.3890 | 0.5016 | 0.1210 | -4.1436 | 0.0000 | 0.0052 | NA |
| TMEM47 | 75.6516 | -0.6470 | 0.1564 | 4.1368 | 0.0000 | 0.0052 | NA |
| SEC61B | 136.7901 | 0.3720 | 0.0900 | -4.1335 | 0.0000 | 0.0052 | NA |
| SEC13 | 114.6709 | 0.2734 | 0.0662 | -4.1293 | 0.0000 | 0.0053 | NA |
| ASS1 | 53.7406 | 0.6461 | 0.1566 | -4.1254 | 0.0000 | 0.0053 | Epithelial |
| YBX1 | 98.0247 | 0.4810 | 0.1168 | -4.1198 | 0.0000 | 0.0054 | NA |
| LTF | 314.4192 | 1.1584 | 0.2811 | -4.1206 | 0.0000 | 0.0054 | Epithelial |
| FH | 41.7578 | 0.4064 | 0.0988 | -4.1151 | 0.0000 | 0.0055 | Epithelial |
| FAR2 | 12.5825 | 0.7846 | 0.1909 | -4.1090 | 0.0000 | 0.0056 | NA |
| RPL12 | 819.5702 | 0.3993 | 0.0973 | -4.1025 | 0.0000 | 0.0057 | NA |
| SDAD1 | 90.9892 | 0.2556 | 0.0623 | -4.0997 | 0.0000 | 0.0057 | NA |
| C5orf46 | 5.6317 | 1.0519 | 0.2567 | -4.0981 | 0.0000 | 0.0057 | NA |
| NIBAN2 | 123.8899 | 0.2598 | 0.0634 | -4.0959 | 0.0000 | 0.0057 | NA |
| RPL23A | 259.6116 | 0.7049 | 0.1721 | -4.0963 | 0.0000 | 0.0057 | NA |
| RPS3 | 1510.5243 | 0.3452 | 0.0843 | -4.0931 | 0.0000 | 0.0057 | NA |
| RN7SKP104 | 19.5474 | -0.6489 | 0.1587 | 4.0877 | 0.0000 | 0.0058 | NA |
| IRAK1 | 148.9408 | 0.2995 | 0.0733 | -4.0876 | 0.0000 | 0.0058 | NA |
| MYO15B | 127.5696 | -0.4012 | 0.0983 | 4.0825 | 0.0000 | 0.0059 | Stromal |
| NOA1 | 25.2287 | 0.3125 | 0.0767 | -4.0716 | 0.0000 | 0.0061 | NA |
| RPL9 | 309.0343 | 0.3698 | 0.0909 | -4.0696 | 0.0000 | 0.0061 | Stromal |
| NDUFB3 | 44.4324 | 0.3507 | 0.0863 | -4.0622 | 0.0000 | 0.0062 | Epithelial |
| AMPH | 14.2340 | -0.8552 | 0.2104 | 4.0648 | 0.0000 | 0.0062 | NA |
| TIMM8B | 74.8903 | 0.3296 | 0.0811 | -4.0627 | 0.0000 | 0.0062 | NA |
| ADIPOR1 | 95.8011 | 0.3492 | 0.0860 | -4.0591 | 0.0000 | 0.0062 | Epithelial |

| | | | | | | | |
|---|---|---|---|---|---|---|---|
| EFNA5 | 58.5736 | 0.6491 | 0.1600 | -4.0563 | 0.0000 | 0.0063 | Epithelial |
| PSMB4 | 165.9808 | 0.2752 | 0.0679 | -4.0541 | 0.0001 | 0.0063 | Epithelial |
| RPL37A | 2411.0963 | 0.2885 | 0.0713 | -4.0469 | 0.0001 | 0.0063 | NA |
| MDM1 | 24.8060 | -0.3696 | 0.0913 | 4.0463 | 0.0001 | 0.0063 | NA |
| RN7SKP118 | 4.6725 | -0.7677 | 0.1897 | 4.0472 | 0.0001 | 0.0063 | NA |
| POLDIP3 | 34.5322 | 0.3045 | 0.0752 | -4.0468 | 0.0001 | 0.0063 | NA |
| AL354861.2 | 4.7135 | -0.5462 | 0.1351 | 4.0441 | 0.0001 | 0.0064 | NA |
| CYCS | 132.4572 | 0.2650 | 0.0656 | -4.0379 | 0.0001 | 0.0064 | Epithelial |
| CHST1 | 48.9111 | 1.1075 | 0.2744 | -4.0368 | 0.0001 | 0.0064 | Epithelial |
| COX7B | 47.5459 | 0.3598 | 0.0891 | -4.0367 | 0.0001 | 0.0064 | Epithelial |
| RPL29 | 166.0592 | 0.3969 | 0.0985 | -4.0298 | 0.0001 | 0.0066 | NA |
| ANP32B | 294.6327 | 0.2124 | 0.0527 | -4.0272 | 0.0001 | 0.0066 | Stromal |
| SFSWAP | 145.2728 | -0.2199 | 0.0547 | 4.0228 | 0.0001 | 0.0067 | NA |
| MON2 | 125.1904 | -0.2369 | 0.0590 | 4.0134 | 0.0001 | 0.0069 | NA |
| RNU1-82P | 3.1328 | -0.7760 | 0.1935 | 4.0106 | 0.0001 | 0.0070 | NA |
| PDZD7 | 3.2627 | -0.8745 | 0.2187 | 3.9976 | 0.0001 | 0.0073 | NA |
| RPS8P6 | 9.8689 | 0.3796 | 0.0950 | -3.9952 | 0.0001 | 0.0074 | Epithelial |
| S100A8 | 195.3024 | 1.3180 | 0.3304 | -3.9889 | 0.0001 | 0.0074 | Epithelial |
| DANCR | 68.1714 | 0.3599 | 0.0903 | -3.9847 | 0.0001 | 0.0074 | Epithelial |
| ZNF862 | 21.2554 | -0.3614 | 0.0907 | 3.9838 | 0.0001 | 0.0074 | NA |
| HOXC6 | 69.9432 | -0.3921 | 0.0984 | 3.9863 | 0.0001 | 0.0074 | NA |
| RPL21 | 77.7876 | 0.6168 | 0.1548 | -3.9842 | 0.0001 | 0.0074 | NA |
| RPL26 | 590.7807 | 0.4073 | 0.1022 | -3.9851 | 0.0001 | 0.0074 | Stromal |
| PSMB3 | 225.4197 | 0.6772 | 0.1699 | -3.9853 | 0.0001 | 0.0074 | Epithelial |
| TTC37 | 181.2177 | -0.2065 | 0.0519 | 3.9786 | 0.0001 | 0.0076 | NA |
| SEM1 | 53.0279 | 0.4108 | 0.1033 | -3.9754 | 0.0001 | 0.0076 | NA |
| PSMA7 | 402.3231 | 0.3117 | 0.0784 | -3.9747 | 0.0001 | 0.0076 | Epithelial |
| RNU2-64P | 27.8676 | 0.6992 | 0.1762 | -3.9677 | 0.0001 | 0.0078 | NA |
| RPL34 | 1340.3513 | 0.2856 | 0.0720 | -3.9660 | 0.0001 | 0.0078 | Stromal |
| MPST | 48.1154 | 0.3537 | 0.0893 | -3.9625 | 0.0001 | 0.0079 | Epithelial |
| AL133355.1 | 1.5128 | -2.0641 | 0.5217 | 3.9565 | 0.0001 | 0.0080 | NA |
| RPS15A | 201.8343 | 0.2544 | 0.0643 | -3.9563 | 0.0001 | 0.0080 | NA |

| | | | | | | | |
|---|---|---|---|---|---|---|---|
| SRCIN1 | 28.1996 | 0.9273 | 0.2347 | -3.9512 | 0.0001 | 0.0081 | Epithelial |
| RN7SL183P | 2.1394 | -0.8376 | 0.2121 | 3.9498 | 0.0001 | 0.0081 | NA |
| MRPL3 | 61.6986 | 0.2576 | 0.0653 | -3.9479 | 0.0001 | 0.0081 | Epithelial |
| TES | 134.8704 | 0.3230 | 0.0818 | -3.9468 | 0.0001 | 0.0081 | Stromal |
| TKT | 277.1517 | 0.3205 | 0.0813 | -3.9404 | 0.0001 | 0.0083 | Epithelial |
| NDUFB8 | 64.9274 | 0.3364 | 0.0854 | -3.9407 | 0.0001 | 0.0083 | NA |
| DLC1 | 133.0172 | -0.4050 | 0.1029 | 3.9356 | 0.0001 | 0.0084 | Stromal |
| AC124068.2 | 3.6993 | -0.8502 | 0.2161 | 3.9351 | 0.0001 | 0.0084 | NA |
| XRCC6 | 148.3686 | 0.2318 | 0.0589 | -3.9334 | 0.0001 | 0.0084 | Epithelial |
| PSMB7 | 137.8070 | 0.2182 | 0.0555 | -3.9312 | 0.0001 | 0.0084 | NA |
| RUNDC3B | 7.6424 | -0.5685 | 0.1448 | 3.9276 | 0.0001 | 0.0085 | Stromal |
| YWHAH | 189.9630 | 0.2664 | 0.0678 | -3.9286 | 0.0001 | 0.0085 | Stromal |
| LSM3 | 79.5947 | 0.2520 | 0.0642 | -3.9257 | 0.0001 | 0.0085 | NA |
| SNORA13 | 51.5161 | -0.5076 | 0.1294 | 3.9243 | 0.0001 | 0.0085 | NA |
| RPS21 | 1535.7095 | 0.3325 | 0.0848 | -3.9186 | 0.0001 | 0.0087 | Epithelial |
| GPATCH8 | 156.6037 | -0.2359 | 0.0603 | 3.9159 | 0.0001 | 0.0087 | Stromal |
| AL391056.1 | 1.5619 | -1.1678 | 0.2983 | 3.9144 | 0.0001 | 0.0087 | NA |
| HELLPAR | 31.4562 | -0.4695 | 0.1201 | 3.9111 | 0.0001 | 0.0088 | Stromal |
| HOXC5 | 2.0586 | -1.2031 | 0.3085 | 3.9004 | 0.0001 | 0.0092 | NA |
| GPAM | 65.7249 | -0.4966 | 0.1274 | 3.8977 | 0.0001 | 0.0092 | Stromal |
| IGIP | 37.2713 | -0.3202 | 0.0822 | 3.8951 | 0.0001 | 0.0093 | Stromal |
| ADH1B | 111.5543 | -0.6726 | 0.1731 | 3.8846 | 0.0001 | 0.0096 | Stromal |
| USP30 | 18.7925 | -0.3309 | 0.0853 | 3.8809 | 0.0001 | 0.0097 | Epithelial |
| TOMM22 | 57.9923 | 0.3284 | 0.0846 | -3.8804 | 0.0001 | 0.0097 | NA |
| ACTG1 | 1799.7067 | 0.2253 | 0.0581 | -3.8772 | 0.0001 | 0.0098 | NA |
| NPM1 | 137.2848 | 0.2925 | 0.0755 | -3.8742 | 0.0001 | 0.0099 | NA |
| OST4 | 232.6950 | 0.2709 | 0.0700 | -3.8685 | 0.0001 | 0.0100 | NA |
| RNU1-89P | 2.6856 | -0.7711 | 0.1993 | 3.8692 | 0.0001 | 0.0100 | NA |
| RN7SL8P | 11.2300 | -0.6568 | 0.1697 | 3.8705 | 0.0001 | 0.0100 | NA |
| MAGI2-AS3 | 66.0857 | -0.3800 | 0.0983 | 3.8658 | 0.0001 | 0.0100 | Stromal |
| ATF4 | 290.8105 | 0.3083 | 0.0798 | -3.8631 | 0.0001 | 0.0101 | NA |
| NPEPPS | 254.1017 | -0.2611 | 0.0676 | 3.8615 | 0.0001 | 0.0101 | Epithelial |

| | | | | | | | |
|---|---|---|---|---|---|---|---|
| C2orf27A_1 | 15.9979 | -0.6479 | 0.1680 | 3.8573 | 0.0001 | 0.0102 | Epithelial |
| CHCHD10 | 148.6283 | 0.3696 | 0.0958 | -3.8566 | 0.0001 | 0.0102 | Stromal |
| SSB | 174.5414 | 0.1695 | 0.0440 | -3.8535 | 0.0001 | 0.0103 | NA |
| LMLN | 66.2284 | -0.2968 | 0.0770 | 3.8545 | 0.0001 | 0.0103 | Epithelial |
| EPGN | 2.7792 | 1.4111 | 0.3665 | -3.8498 | 0.0001 | 0.0103 | NA |
| RSL24D1 | 76.0448 | 0.2682 | 0.0697 | -3.8503 | 0.0001 | 0.0103 | NA |
| DNAJC8 | 92.5259 | 0.2165 | 0.0563 | -3.8443 | 0.0001 | 0.0105 | Stromal |
| PQBP1 | 53.2964 | 0.2548 | 0.0663 | -3.8436 | 0.0001 | 0.0105 | NA |
| RFC4 | 22.6743 | 0.3599 | 0.0937 | -3.8406 | 0.0001 | 0.0106 | Epithelial |
| PGD | 59.0110 | 0.3964 | 0.1033 | -3.8388 | 0.0001 | 0.0106 | Epithelial |
| ATP5MD | 130.2745 | 0.3247 | 0.0846 | -3.8361 | 0.0001 | 0.0107 | Epithelial |
| ANKRD30B | 151.5538 | -1.1317 | 0.2954 | 3.8313 | 0.0001 | 0.0108 | Epithelial |
| TSHZ1 | 92.8341 | -0.2344 | 0.0612 | 3.8322 | 0.0001 | 0.0108 | Stromal |
| MIR320E | 3.9902 | -1.0044 | 0.2628 | 3.8219 | 0.0001 | 0.0112 | Stromal |
| PHF21A | 83.7804 | -0.2056 | 0.0538 | 3.8205 | 0.0001 | 0.0112 | NA |
| YDJC | 38.8244 | 0.3196 | 0.0837 | -3.8179 | 0.0001 | 0.0112 | Epithelial |
| TSPAN15 | 53.2259 | 0.4957 | 0.1299 | -3.8158 | 0.0001 | 0.0113 | Epithelial |
| SP1 | 150.5837 | -0.1832 | 0.0480 | 3.8132 | 0.0001 | 0.0114 | NA |
| EIF2S3 | 343.8412 | 0.1856 | 0.0487 | -3.8100 | 0.0001 | 0.0115 | Epithelial |
| RPS25 | 319.4263 | 0.3445 | 0.0905 | -3.8051 | 0.0001 | 0.0116 | Stromal |
| MPHOSPH6 | 89.7900 | 0.5828 | 0.1531 | -3.8059 | 0.0001 | 0.0116 | Epithelial |
| RPL37 | 1081.8818 | 0.3530 | 0.0928 | -3.8027 | 0.0001 | 0.0116 | NA |
| PPIB | 390.6639 | 0.2711 | 0.0713 | -3.8036 | 0.0001 | 0.0116 | Stromal |
| HSPE1 | 92.8636 | 0.3921 | 0.1032 | -3.7998 | 0.0001 | 0.0117 | Epithelial |
| CARTPT | 18.5166 | 2.6350 | 0.6939 | -3.7974 | 0.0001 | 0.0117 | Epithelial |
| SNORD124 | 19.2914 | -0.7209 | 0.1898 | 3.7977 | 0.0001 | 0.0117 | NA |
| GASK1B | 213.6539 | -0.4953 | 0.1305 | 3.7946 | 0.0001 | 0.0118 | NA |
| PFDN2 | 88.6498 | 0.3052 | 0.0805 | -3.7923 | 0.0001 | 0.0119 | NA |
| DUBR | 22.9058 | -0.4992 | 0.1318 | 3.7885 | 0.0002 | 0.0120 | NA |
| AC009283.1 | 27.8807 | 0.9488 | 0.2505 | -3.7875 | 0.0002 | 0.0120 | Epithelial |
| NDUFB7 | 162.3880 | 0.2362 | 0.0624 | -3.7860 | 0.0002 | 0.0120 | Epithelial |
| EIF4G1 | 391.1727 | 0.1791 | 0.0473 | -3.7828 | 0.0002 | 0.0121 | Epithelial |

| | | | | | | | |
|---|---|---|---|---|---|---|---|
| STK36 | 52.1085 | -0.3190 | 0.0844 | 3.7811 | 0.0002 | 0.0122 | Epithelial |
| RPS11 | 1898.7726 | 0.2667 | 0.0706 | -3.7782 | 0.0002 | 0.0123 | Stromal |
| CRTC2 | 26.9653 | 0.2960 | 0.0784 | -3.7766 | 0.0002 | 0.0123 | NA |
| RPL31 | 2163.4680 | 0.2399 | 0.0636 | -3.7690 | 0.0002 | 0.0126 | NA |
| RPS18 | 145.3164 | 0.6948 | 0.1846 | -3.7642 | 0.0002 | 0.0128 | NA |
| PEX5L | 6.0398 | -1.3348 | 0.3550 | 3.7605 | 0.0002 | 0.0130 | NA |
| ZNF331 | 52.1755 | -0.3480 | 0.0926 | 3.7595 | 0.0002 | 0.0130 | NA |
| RN7SL277P | 2.3939 | -0.7965 | 0.2120 | 3.7563 | 0.0002 | 0.0131 | NA |
| GNMT | 11.9556 | 0.9418 | 0.2510 | -3.7530 | 0.0002 | 0.0131 | Epithelial |
| MPZL3 | 13.9105 | 0.4399 | 0.1172 | -3.7539 | 0.0002 | 0.0131 | NA |
| PRPF39 | 60.4316 | -0.2757 | 0.0734 | 3.7536 | 0.0002 | 0.0131 | NA |
| MAP4K5 | 121.8216 | -0.2183 | 0.0582 | 3.7531 | 0.0002 | 0.0131 | Stromal |
| KCNS3 | 19.2770 | 0.5242 | 0.1400 | -3.7441 | 0.0002 | 0.0135 | NA |
| SNORD67 | 14.3998 | -0.4702 | 0.1257 | 3.7392 | 0.0002 | 0.0136 | NA |
| TUBA1C | 74.1684 | 0.4109 | 0.1099 | -3.7386 | 0.0002 | 0.0136 | Epithelial |
| AC103702.1 | 8.0264 | -1.0572 | 0.2826 | 3.7411 | 0.0002 | 0.0136 | NA |
| PITPNB | 70.2844 | 0.2656 | 0.0710 | -3.7391 | 0.0002 | 0.0136 | Stromal |
| RPS26 | 51.2479 | 0.5964 | 0.1596 | -3.7376 | 0.0002 | 0.0136 | NA |
| PPT2-EGFL8 | 5.7858 | -0.6341 | 0.1699 | 3.7326 | 0.0002 | 0.0137 | NA |
| NDUFA6 | 123.6170 | 0.2872 | 0.0769 | -3.7333 | 0.0002 | 0.0137 | Epithelial |
| LRRC8D | 18.6202 | 0.3653 | 0.0981 | -3.7243 | 0.0002 | 0.0139 | NA |
| RN7SL752P | 11.5498 | -0.6391 | 0.1714 | 3.7280 | 0.0002 | 0.0139 | NA |
| SNORA20 | 19.2818 | -0.5037 | 0.1352 | 3.7242 | 0.0002 | 0.0139 | NA |
| ATP5MF | 182.4768 | 0.2135 | 0.0573 | -3.7272 | 0.0002 | 0.0139 | Epithelial |
| RPLP0 | 645.0857 | 0.3616 | 0.0971 | -3.7245 | 0.0002 | 0.0139 | NA |
| FASN | 446.4476 | 0.5658 | 0.1519 | -3.7254 | 0.0002 | 0.0139 | Epithelial |
| EXOSC5 | 27.6071 | 0.3334 | 0.0895 | -3.7235 | 0.0002 | 0.0139 | Epithelial |
| C2orf50 | 7.0972 | -0.9094 | 0.2444 | 3.7210 | 0.0002 | 0.0139 | NA |
| LINC02055 | 3.0199 | -1.1593 | 0.3116 | 3.7208 | 0.0002 | 0.0139 | NA |
| RAPGEF3 | 42.6829 | -0.3575 | 0.0961 | 3.7216 | 0.0002 | 0.0139 | Stromal |
| SURF2 | 25.7062 | 0.2755 | 0.0741 | -3.7197 | 0.0002 | 0.0139 | Epithelial |
| LDHA | 81.0751 | 0.3668 | 0.0987 | -3.7179 | 0.0002 | 0.0139 | NA |

| | | | | | | | |
|---|---|---|---|---|---|---|---|
| PTRH2 | 43.8323 | 0.4839 | 0.1302 | -3.7181 | 0.0002 | 0.0139 | Epithelial |
| MTRNR2L6 | 28.1875 | -0.4320 | 0.1162 | 3.7169 | 0.0002 | 0.0139 | NA |
| BTNL9 | 59.4652 | -0.4962 | 0.1337 | 3.7110 | 0.0002 | 0.0142 | Stromal |
| RNF170 | 55.3759 | -0.3188 | 0.0860 | 3.7078 | 0.0002 | 0.0142 | NA |
| HOXC8 | 58.2411 | -0.4816 | 0.1299 | 3.7085 | 0.0002 | 0.0142 | NA |
| ANXA2 | 313.0419 | 0.3210 | 0.0865 | -3.7098 | 0.0002 | 0.0142 | NA |
| EBP | 48.5206 | 0.3283 | 0.0885 | -3.7083 | 0.0002 | 0.0142 | Epithelial |
| DYNC2H1 | 98.1934 | -0.3325 | 0.0897 | 3.7047 | 0.0002 | 0.0143 | Stromal |
| NTRK3 | 15.0842 | -0.6135 | 0.1657 | 3.7017 | 0.0002 | 0.0144 | Stromal |
| TTC1 | 101.4034 | 0.2108 | 0.0570 | -3.6996 | 0.0002 | 0.0145 | NA |
| BAG4 | 45.9235 | -0.4433 | 0.1199 | 3.6969 | 0.0002 | 0.0145 | Epithelial |
| HECTD2 | 35.1453 | -0.4308 | 0.1165 | 3.6980 | 0.0002 | 0.0145 | NA |
| MIF | 1.2749 | 1.1345 | 0.3069 | -3.6970 | 0.0002 | 0.0145 | NA |
| TRPC1 | 17.4067 | -0.4304 | 0.1165 | 3.6953 | 0.0002 | 0.0145 | Stromal |
| ABCA6 | 75.7506 | -0.4696 | 0.1273 | 3.6904 | 0.0002 | 0.0147 | Stromal |
| UBA52 | 610.1536 | 0.2446 | 0.0663 | -3.6910 | 0.0002 | 0.0147 | NA |
| LDHB | 125.0073 | 0.4971 | 0.1349 | -3.6858 | 0.0002 | 0.0149 | Stromal |
| KRR1 | 190.5768 | -0.2011 | 0.0546 | 3.6846 | 0.0002 | 0.0149 | Epithelial |
| DHX33 | 70.6983 | -0.2293 | 0.0622 | 3.6854 | 0.0002 | 0.0149 | NA |
| SLC25A5 | 90.3456 | 0.4516 | 0.1225 | -3.6864 | 0.0002 | 0.0149 | Epithelial |
| POLN | 17.3296 | -0.5721 | 0.1554 | 3.6826 | 0.0002 | 0.0149 | Epithelial |
| AC106037.1 | 1.7150 | -1.0507 | 0.2853 | 3.6833 | 0.0002 | 0.0149 | NA |
| SNORA75 | 82.6076 | -0.5721 | 0.1554 | 3.6815 | 0.0002 | 0.0149 | Epithelial |
| SMARCE1 | 67.0011 | -0.4998 | 0.1358 | 3.6793 | 0.0002 | 0.0149 | Epithelial |
| ECH1 | 123.0274 | 0.2912 | 0.0791 | -3.6800 | 0.0002 | 0.0149 | NA |
| HSPD1 | 203.5319 | 0.2725 | 0.0741 | -3.6782 | 0.0002 | 0.0149 | Epithelial |
| HMGCS1 | 90.4131 | 0.4169 | 0.1134 | -3.6753 | 0.0002 | 0.0150 | Epithelial |
| TMEM250 | 52.0267 | 0.2359 | 0.0642 | -3.6756 | 0.0002 | 0.0150 | NA |
| AC005670.3 | 28.1117 | -0.3235 | 0.0880 | 3.6761 | 0.0002 | 0.0150 | Stromal |
| RALGAPA1 | 84.4045 | -0.2491 | 0.0678 | 3.6732 | 0.0002 | 0.0151 | NA |
| GDAP1 | 46.8261 | -0.7848 | 0.2138 | 3.6716 | 0.0002 | 0.0151 | Epithelial |
| RPL23AP42 | 9.6966 | 0.7471 | 0.2036 | -3.6696 | 0.0002 | 0.0151 | NA |

| | | | | | | | |
|---|---|---|---|---|---|---|---|
| STK24 | 88.9784 | 0.2070 | 0.0564 | -3.6695 | 0.0002 | 0.0151 | Stromal |
| UBXN4 | 472.0169 | 0.1464 | 0.0400 | -3.6636 | 0.0002 | 0.0152 | Epithelial |
| MMADHC | 71.0778 | 0.2179 | 0.0594 | -3.6654 | 0.0002 | 0.0152 | NA |
| TDRD3 | 71.4517 | -0.2598 | 0.0709 | 3.6643 | 0.0002 | 0.0152 | Stromal |
| MPP2 | 11.9645 | -0.7737 | 0.2112 | 3.6638 | 0.0002 | 0.0152 | Epithelial |
| IL1RAPL1 | 1.7168 | -0.9888 | 0.2699 | 3.6641 | 0.0002 | 0.0152 | NA |
| PDAP1 | 32.8366 | 0.3397 | 0.0928 | -3.6620 | 0.0003 | 0.0153 | NA |
| PABPC1L | 50.9472 | -0.4318 | 0.1180 | 3.6592 | 0.0003 | 0.0154 | Epithelial |
| PSMB6 | 95.9690 | 0.2636 | 0.0721 | -3.6566 | 0.0003 | 0.0155 | Epithelial |
| LRRC37B | 6.5809 | 0.7331 | 0.2007 | -3.6530 | 0.0003 | 0.0157 | NA |
| MTHFD2 | 150.0399 | 0.3414 | 0.0935 | -3.6510 | 0.0003 | 0.0158 | Epithelial |
| NACA | 183.0502 | 0.2431 | 0.0666 | -3.6497 | 0.0003 | 0.0158 | NA |
| YWHAE | 316.3479 | 0.2937 | 0.0805 | -3.6487 | 0.0003 | 0.0158 | Epithelial |
| EDF1 | 383.6365 | 0.1861 | 0.0510 | -3.6474 | 0.0003 | 0.0158 | Epithelial |
| GABBR2 | 2.7327 | 1.3440 | 0.3689 | -3.6435 | 0.0003 | 0.0160 | NA |
| RN7SL832P | 8.0411 | -0.5117 | 0.1406 | 3.6400 | 0.0003 | 0.0161 | Stromal |
| RPL14 | 143.4067 | 0.2616 | 0.0719 | -3.6407 | 0.0003 | 0.0161 | NA |
| FABP4 | 169.0150 | -0.7346 | 0.2018 | 3.6409 | 0.0003 | 0.0161 | Stromal |
| BOLA3 | 26.6999 | 0.2998 | 0.0825 | -3.6340 | 0.0003 | 0.0163 | Epithelial |
| LINC00882 | 8.3753 | -0.5897 | 0.1622 | 3.6353 | 0.0003 | 0.0163 | NA |
| NSD3 | 305.9557 | -0.3824 | 0.1053 | 3.6330 | 0.0003 | 0.0163 | Epithelial |
| ZW10 | 21.5654 | 0.3048 | 0.0839 | -3.6328 | 0.0003 | 0.0163 | NA |
| RUNDC3A | 2.7419 | -1.5516 | 0.4269 | 3.6349 | 0.0003 | 0.0163 | NA |
| UBE2L3 | 75.3491 | 0.2635 | 0.0725 | -3.6359 | 0.0003 | 0.0163 | Epithelial |
| PABPC1 | 2791.5554 | 0.3458 | 0.0953 | -3.6307 | 0.0003 | 0.0163 | Epithelial |
| AVPR1A | 11.6101 | -0.5344 | 0.1472 | 3.6310 | 0.0003 | 0.0163 | Stromal |
| RPL36 | 567.5612 | 0.3812 | 0.1050 | -3.6293 | 0.0003 | 0.0163 | NA |
| CD163L1 | 14.0522 | 0.5722 | 0.1578 | -3.6263 | 0.0003 | 0.0165 | NA |
| IRGQ | 73.3257 | 0.2279 | 0.0628 | -3.6256 | 0.0003 | 0.0165 | Epithelial |
| SNHG5 | 9226.9532 | -0.4118 | 0.1137 | 3.6217 | 0.0003 | 0.0166 | Stromal |
| COMT | 81.7900 | 0.3657 | 0.1010 | -3.6218 | 0.0003 | 0.0166 | Epithelial |
| PRDX4 | 62.0714 | 0.3154 | 0.0871 | -3.6209 | 0.0003 | 0.0166 | NA |

| | | | | | | | |
|---|---|---|---|---|---|---|---|
| ATP5PO | 143.9241 | 0.2388 | 0.0660 | -3.6198 | 0.0003 | 0.0167 | NA |
| CLIC1 | 304.5624 | 0.1909 | 0.0529 | -3.6110 | 0.0003 | 0.0172 | NA |
| FADS2 | 58.2913 | 0.8232 | 0.2280 | -3.6106 | 0.0003 | 0.0172 | Epithelial |
| UFC1 | 188.4449 | 0.3014 | 0.0836 | -3.6074 | 0.0003 | 0.0173 | Epithelial |
| EZH1 | 70.9775 | -0.2599 | 0.0721 | 3.6069 | 0.0003 | 0.0173 | Stromal |
| EDN2 | 2.8718 | 1.2316 | 0.3416 | -3.6054 | 0.0003 | 0.0173 | NA |
| ABCA9 | 59.8059 | -0.4916 | 0.1363 | 3.6058 | 0.0003 | 0.0173 | Stromal |
| SOX2-OT | 8.4246 | -0.6734 | 0.1869 | 3.6022 | 0.0003 | 0.0174 | NA |
| RPS24 | 1643.4599 | 0.2675 | 0.0743 | -3.6020 | 0.0003 | 0.0174 | NA |
| UVSSA | 69.2945 | -0.3038 | 0.0844 | 3.5998 | 0.0003 | 0.0175 | Epithelial |
| RPS8 | 1418.8359 | 0.3567 | 0.0992 | -3.5947 | 0.0003 | 0.0177 | Stromal |
| ZFHX4 | 103.0384 | -0.4200 | 0.1168 | 3.5947 | 0.0003 | 0.0177 | Stromal |
| SRPRA | 126.5176 | 0.2419 | 0.0673 | -3.5931 | 0.0003 | 0.0177 | NA |
| UBALD2 | 102.4898 | 0.3290 | 0.0915 | -3.5951 | 0.0003 | 0.0177 | NA |
| BIRC5 | 33.5345 | 0.5915 | 0.1646 | -3.5939 | 0.0003 | 0.0177 | Epithelial |
| PSMG1 | 37.4765 | 0.2836 | 0.0789 | -3.5929 | 0.0003 | 0.0177 | Epithelial |
| MID1IP1 | 45.0926 | 0.3154 | 0.0877 | -3.5941 | 0.0003 | 0.0177 | NA |
| IDH2 | 176.7698 | 0.4095 | 0.1140 | -3.5919 | 0.0003 | 0.0177 | Epithelial |
| OXCT1 | 26.8535 | 0.4402 | 0.1227 | -3.5875 | 0.0003 | 0.0178 | NA |
| HYMAI | 7.7609 | -0.7333 | 0.2044 | 3.5874 | 0.0003 | 0.0178 | Stromal |
| SERF2 | 467.3476 | 0.3300 | 0.0920 | -3.5875 | 0.0003 | 0.0178 | Epithelial |
| IGBP1 | 67.5333 | 0.2407 | 0.0671 | -3.5866 | 0.0003 | 0.0178 | NA |
| DKC1 | 140.8681 | 0.2416 | 0.0674 | -3.5863 | 0.0003 | 0.0178 | Epithelial |
| SPCS2 | 14.6954 | 0.5223 | 0.1457 | -3.5852 | 0.0003 | 0.0178 | NA |
| TRPM7 | 139.2131 | -0.2014 | 0.0562 | 3.5848 | 0.0003 | 0.0178 | Stromal |
| SRD5A3 | 58.7737 | 0.4564 | 0.1274 | -3.5823 | 0.0003 | 0.0180 | Epithelial |
| EIF3D | 170.8675 | 0.2263 | 0.0632 | -3.5816 | 0.0003 | 0.0180 | NA |
| PSD3 | 115.8670 | -0.6910 | 0.1931 | 3.5773 | 0.0003 | 0.0182 | Epithelial |
| Z82217.1 | 4.2143 | -0.6607 | 0.1847 | 3.5763 | 0.0003 | 0.0182 | NA |
| SYP | 5.7808 | -0.8035 | 0.2250 | 3.5712 | 0.0004 | 0.0185 | NA |
| MRPL9 | 67.0801 | 0.2109 | 0.0591 | -3.5693 | 0.0004 | 0.0186 | NA |
| RPL15 | 1039.2036 | 0.2341 | 0.0656 | -3.5665 | 0.0004 | 0.0187 | Stromal |

| | | | | | | | |
|---|---|---|---|---|---|---|---|
| KRT8 | 125.7984 | 0.4565 | 0.1280 | -3.5666 | 0.0004 | 0.0187 | Epithelial |
| EIF5A | 77.8317 | 0.4260 | 0.1195 | -3.5650 | 0.0004 | 0.0187 | NA |
| MBD2 | 180.6362 | 0.1732 | 0.0486 | -3.5649 | 0.0004 | 0.0187 | Epithelial |
| REV3L | 184.5734 | -0.2466 | 0.0692 | 3.5640 | 0.0004 | 0.0188 | Stromal |
| ATP5MPL | 218.1185 | 0.2022 | 0.0567 | -3.5625 | 0.0004 | 0.0188 | Epithelial |
| AC022007.1 | 7.2901 | 0.5921 | 0.1665 | -3.5568 | 0.0004 | 0.0190 | Epithelial |
| RACK1 | 1781.0099 | 0.2181 | 0.0613 | -3.5559 | 0.0004 | 0.0190 | Stromal |
| AUXG01000058.1 | 26.8904 | -0.4485 | 0.1260 | 3.5591 | 0.0004 | 0.0190 | NA |
| SAPCD2 | 34.0289 | 0.5327 | 0.1498 | -3.5572 | 0.0004 | 0.0190 | Epithelial |
| TMTC2 | 46.0243 | -0.4194 | 0.1179 | 3.5563 | 0.0004 | 0.0190 | Stromal |
| SNRPD2 | 246.3727 | 0.1962 | 0.0552 | -3.5568 | 0.0004 | 0.0190 | Epithelial |
| GALNT14 | 6.6549 | 0.8153 | 0.2293 | -3.5552 | 0.0004 | 0.0190 | Epithelial |
| MIEF1 | 57.1078 | 0.2467 | 0.0695 | -3.5502 | 0.0004 | 0.0193 | NA |
| ACTB | 216.6451 | 0.2614 | 0.0737 | -3.5481 | 0.0004 | 0.0193 | Stromal |
| SLC31A1 | 77.0154 | 0.3344 | 0.0942 | -3.5487 | 0.0004 | 0.0193 | Epithelial |
| HDC | 14.0639 | -0.8233 | 0.2320 | 3.5489 | 0.0004 | 0.0193 | Stromal |
| COX6B1 | 331.3832 | 0.2345 | 0.0661 | -3.5470 | 0.0004 | 0.0194 | Epithelial |
| URM1 | 79.3383 | 0.1939 | 0.0547 | -3.5455 | 0.0004 | 0.0194 | NA |
| E2F2 | 4.7378 | 0.6592 | 0.1860 | -3.5440 | 0.0004 | 0.0195 | Epithelial |
| POLD1 | 21.4023 | 0.3075 | 0.0868 | -3.5424 | 0.0004 | 0.0196 | NA |
| RPL30 | 579.4401 | 0.3016 | 0.0852 | -3.5414 | 0.0004 | 0.0196 | NA |
| CSTB | 190.7836 | 0.3920 | 0.1109 | -3.5351 | 0.0004 | 0.0200 | Epithelial |
| ITGB6 | 76.5047 | 0.6551 | 0.1854 | -3.5342 | 0.0004 | 0.0200 | Epithelial |
| PCAT6 | 8.0406 | 0.5929 | 0.1678 | -3.5328 | 0.0004 | 0.0201 | NA |
| CEP126 | 70.5061 | -0.4472 | 0.1267 | 3.5302 | 0.0004 | 0.0202 | Stromal |
| MT2A | 267.3290 | 0.5046 | 0.1430 | -3.5300 | 0.0004 | 0.0202 | NA |
| RPL10A | 175.1183 | 0.4176 | 0.1184 | -3.5261 | 0.0004 | 0.0204 | NA |
| UHRF1BP1L | 70.1374 | -0.2079 | 0.0590 | 3.5261 | 0.0004 | 0.0204 | NA |
| CD48 | 16.4704 | 0.7369 | 0.2090 | -3.5254 | 0.0004 | 0.0204 | Stromal |
| LAMA2 | 79.9367 | -0.3690 | 0.1047 | 3.5247 | 0.0004 | 0.0204 | Stromal |
| HSP90AA1 | 1519.0752 | 0.2877 | 0.0816 | -3.5241 | 0.0004 | 0.0204 | Epithelial |
| SIAH2-AS1 | 7.3244 | -1.0668 | 0.3032 | 3.5184 | 0.0004 | 0.0206 | Epithelial |

| | | | | | | | |
|---|---|---|---|---|---|---|---|
| RN7SL838P | 18.9337 | -0.5960 | 0.1694 | 3.5184 | 0.0004 | 0.0206 | NA |
| AC004223.2 | 4.4292 | -0.8046 | 0.2287 | 3.5187 | 0.0004 | 0.0206 | NA |
| MYL12A | 252.3594 | 0.2709 | 0.0770 | -3.5178 | 0.0004 | 0.0206 | Stromal |
| CLDN14 | 1.4793 | 1.0866 | 0.3089 | -3.5181 | 0.0004 | 0.0206 | NA |
| AL391832.2 | 7.1015 | 0.6034 | 0.1718 | -3.5117 | 0.0004 | 0.0210 | Epithelial |
| CFAP70 | 56.4245 | -0.4771 | 0.1359 | 3.5111 | 0.0004 | 0.0210 | Epithelial |
| PDGFD | 42.9744 | -0.4255 | 0.1212 | 3.5116 | 0.0004 | 0.0210 | Stromal |
| DNAH9 | 1.7871 | -1.2553 | 0.3575 | 3.5111 | 0.0004 | 0.0210 | NA |
| RPL11 | 1607.0729 | 0.1998 | 0.0570 | -3.5027 | 0.0005 | 0.0210 | Stromal |
| UQCRH | 229.0603 | 0.2736 | 0.0781 | -3.5027 | 0.0005 | 0.0210 | Epithelial |
| S100A7A | 10.1513 | 2.5175 | 0.7184 | -3.5043 | 0.0005 | 0.0210 | Epithelial |
| ILF2 | 134.8824 | 0.2307 | 0.0658 | -3.5078 | 0.0005 | 0.0210 | Epithelial |
| CAPN13 | 24.3169 | 0.7130 | 0.2036 | -3.5025 | 0.0005 | 0.0210 | Epithelial |
| CMSS1 | 48.9005 | 0.3070 | 0.0877 | -3.5027 | 0.0005 | 0.0210 | NA |
| AC109347.2 | 2.1030 | 0.8332 | 0.2378 | -3.5044 | 0.0005 | 0.0210 | NA |
| ST14 | 96.0440 | 0.3801 | 0.1084 | -3.5078 | 0.0005 | 0.0210 | Epithelial |
| CHAC1 | 4.0519 | 0.8981 | 0.2564 | -3.5032 | 0.0005 | 0.0210 | NA |
| CIAPIN1 | 19.4517 | 0.3510 | 0.1001 | -3.5067 | 0.0005 | 0.0210 | NA |
| EEF2 | 1944.0799 | 0.2050 | 0.0585 | -3.5068 | 0.0005 | 0.0210 | NA |
| PSMD8 | 244.2775 | 0.2654 | 0.0757 | -3.5054 | 0.0005 | 0.0210 | Epithelial |
| STARD7 | 97.3247 | 0.1875 | 0.0536 | -3.5010 | 0.0005 | 0.0211 | NA |
| HSD17B10 | 79.8656 | 0.2635 | 0.0753 | -3.4996 | 0.0005 | 0.0212 | Epithelial |
| WTAP | 73.9677 | 0.2258 | 0.0646 | -3.4986 | 0.0005 | 0.0212 | NA |
| AC062004.1 | 2.2544 | -1.0333 | 0.2955 | 3.4970 | 0.0005 | 0.0212 | Stromal |
| LGI1 | 2.0735 | -0.8941 | 0.2556 | 3.4973 | 0.0005 | 0.0212 | NA |
| KCNC2 | 69.7373 | -1.6374 | 0.4683 | 3.4963 | 0.0005 | 0.0212 | Epithelial |
| SNORA38B | 12.2830 | -0.6558 | 0.1876 | 3.4954 | 0.0005 | 0.0213 | Epithelial |
| EFHC1 | 58.3314 | -0.3259 | 0.0934 | 3.4888 | 0.0005 | 0.0216 | Epithelial |
| DHCR7 | 55.9065 | 0.5131 | 0.1471 | -3.4882 | 0.0005 | 0.0216 | Epithelial |
| ZNF26 | 65.1569 | -0.2758 | 0.0790 | 3.4904 | 0.0005 | 0.0216 | Epithelial |
| CES3 | 4.3031 | 1.2790 | 0.3666 | -3.4887 | 0.0005 | 0.0216 | NA |
| ZNF44 | 69.0307 | -0.3238 | 0.0928 | 3.4882 | 0.0005 | 0.0216 | NA |

| | | | | | | | |
|---|---|---|---|---|---|---|---|
| ACO2 | 98.4239 | 0.2376 | 0.0681 | -3.4902 | 0.0005 | 0.0216 | Epithelial |
| UQCRB | 257.9997 | 0.2499 | 0.0717 | -3.4863 | 0.0005 | 0.0217 | NA |
| POMP | 142.7961 | 0.2221 | 0.0638 | -3.4839 | 0.0005 | 0.0218 | Epithelial |
| ZNF518A | 149.9459 | -0.2173 | 0.0624 | 3.4827 | 0.0005 | 0.0219 | Epithelial |
| FAM89B | 104.1775 | 0.2240 | 0.0643 | -3.4812 | 0.0005 | 0.0219 | NA |
| RPS5 | 773.5341 | 0.2798 | 0.0804 | -3.4781 | 0.0005 | 0.0221 | NA |
| AC114490.3 | 8.4655 | -0.5209 | 0.1499 | 3.4740 | 0.0005 | 0.0223 | NA |
| EIF4A2 | 210.5167 | 0.2102 | 0.0605 | -3.4738 | 0.0005 | 0.0223 | NA |
| ZFYVE16 | 163.3559 | -0.1931 | 0.0556 | 3.4737 | 0.0005 | 0.0223 | Stromal |
| OLFML2A | 63.6707 | -0.3283 | 0.0945 | 3.4741 | 0.0005 | 0.0223 | Stromal |
| AC005921.4 | 7.0003 | -0.6337 | 0.1824 | 3.4735 | 0.0005 | 0.0223 | NA |
| CCT6A | 240.3929 | 0.1985 | 0.0572 | -3.4713 | 0.0005 | 0.0224 | Epithelial |
| ARF4 | 80.9065 | 0.2889 | 0.0832 | -3.4704 | 0.0005 | 0.0224 | NA |
| MINDY3 | 32.9399 | -0.2910 | 0.0839 | 3.4701 | 0.0005 | 0.0224 | NA |
| LMNB2 | 30.0392 | 0.2786 | 0.0803 | -3.4685 | 0.0005 | 0.0225 | Epithelial |
| CEP290 | 207.9039 | -0.2385 | 0.0688 | 3.4672 | 0.0005 | 0.0226 | NA |
| VAMP8 | 236.7002 | 0.2375 | 0.0685 | -3.4662 | 0.0005 | 0.0226 | Epithelial |
| TMA7 | 78.1268 | 0.2935 | 0.0847 | -3.4656 | 0.0005 | 0.0226 | NA |
| RPS20P22 | 17.0242 | -0.6370 | 0.1840 | 3.4630 | 0.0005 | 0.0227 | NA |
| RPL8 | 2501.6919 | 0.3032 | 0.0875 | -3.4634 | 0.0005 | 0.0227 | Epithelial |
| GRB14 | 33.5959 | -1.2205 | 0.3525 | 3.4620 | 0.0005 | 0.0227 | Epithelial |
| ZNF236 | 57.4095 | -0.2533 | 0.0732 | 3.4615 | 0.0005 | 0.0227 | Epithelial |
| MT-ND4 | 11168.5621 | 0.3125 | 0.0903 | -3.4605 | 0.0005 | 0.0228 | Epithelial |
| CERS5 | 51.1223 | -0.2097 | 0.0606 | 3.4594 | 0.0005 | 0.0228 | NA |
| RP9 | 16.7409 | 0.3337 | 0.0966 | -3.4551 | 0.0006 | 0.0231 | NA |
| AC004825.3 | 2.8839 | -0.8011 | 0.2318 | 3.4558 | 0.0005 | 0.0231 | Stromal |
| MED1 | 401.9248 | 0.6233 | 0.1804 | -3.4548 | 0.0006 | 0.0231 | Epithelial |
| DHX16 | 51.7999 | -0.2590 | 0.0751 | 3.4490 | 0.0006 | 0.0235 | NA |
| ITGA2B | 1.9689 | -0.8818 | 0.2557 | 3.4488 | 0.0006 | 0.0235 | NA |
| PFKFB2 | 22.8852 | 0.4161 | 0.1208 | -3.4445 | 0.0006 | 0.0236 | NA |
| MPV17L | 40.6703 | 0.7253 | 0.2105 | -3.4463 | 0.0006 | 0.0236 | Epithelial |
| AC091153.1 | 2.0645 | 0.7727 | 0.2243 | -3.4441 | 0.0006 | 0.0236 | NA |

| | | | | | | | |
|---|---|---|---|---|---|---|---|
| MIA | 18.5782 | 0.6622 | 0.1922 | -3.4445 | 0.0006 | 0.0236 | NA |
| UFD1 | 71.2209 | 0.2409 | 0.0699 | -3.4459 | 0.0006 | 0.0236 | NA |
| PIN4 | 41.6164 | 0.2638 | 0.0766 | -3.4438 | 0.0006 | 0.0236 | Epithelial |
| S100A9 | 450.5395 | 1.0411 | 0.3026 | -3.4409 | 0.0006 | 0.0238 | Epithelial |
| PCSK2 | 1.5712 | -1.2019 | 0.3495 | 3.4389 | 0.0006 | 0.0240 | NA |
| LPL | 84.5276 | -0.5796 | 0.1686 | 3.4370 | 0.0006 | 0.0240 | Stromal |
| CHP1 | 97.8210 | 0.2366 | 0.0688 | -3.4372 | 0.0006 | 0.0240 | Epithelial |
| TXNL4A | 133.6549 | 0.2374 | 0.0691 | -3.4371 | 0.0006 | 0.0240 | Epithelial |
| SCN8A | 13.5754 | -0.7602 | 0.2214 | 3.4341 | 0.0006 | 0.0241 | Epithelial |
| STXBP6 | 4.1166 | -0.7681 | 0.2236 | 3.4346 | 0.0006 | 0.0241 | NA |
| SAP30 | 17.8093 | 0.3905 | 0.1138 | -3.4324 | 0.0006 | 0.0241 | Epithelial |
| HERC1 | 213.5943 | -0.2271 | 0.0661 | 3.4325 | 0.0006 | 0.0241 | Stromal |
| MT-ND4L | 238.5799 | 0.3945 | 0.1149 | -3.4334 | 0.0006 | 0.0241 | Epithelial |
| PLA2G2D | 7.5350 | 0.9721 | 0.2833 | -3.4309 | 0.0006 | 0.0242 | NA |
| MYOZ3 | 3.7975 | -0.8959 | 0.2611 | 3.4308 | 0.0006 | 0.0242 | NA |
| BRCA1 | 37.6428 | -0.3946 | 0.1151 | 3.4295 | 0.0006 | 0.0242 | Epithelial |
| ST6GAL1 | 44.3851 | 0.5323 | 0.1552 | -3.4289 | 0.0006 | 0.0243 | Stromal |
| PRR11 | 40.9090 | 0.5558 | 0.1623 | -3.4243 | 0.0006 | 0.0246 | NA |
| EXOC4 | 124.1939 | -0.2010 | 0.0587 | 3.4232 | 0.0006 | 0.0247 | NA |
| TMSB10 | 1768.3385 | 0.3286 | 0.0960 | -3.4218 | 0.0006 | 0.0247 | NA |
| SLC35C1 | 26.7921 | 0.3887 | 0.1136 | -3.4221 | 0.0006 | 0.0247 | NA |
| RARRES1 | 101.2983 | 0.6837 | 0.1999 | -3.4200 | 0.0006 | 0.0248 | NA |
| PFN1 | 678.8930 | 0.2656 | 0.0777 | -3.4190 | 0.0006 | 0.0249 | Stromal |
| ATP1A1 | 323.9202 | 0.2442 | 0.0714 | -3.4178 | 0.0006 | 0.0249 | Epithelial |
| AP005121.1 | 9.8503 | -1.3019 | 0.3809 | 3.4181 | 0.0006 | 0.0249 | Epithelial |
| NPHP3 | 72.6266 | -0.2324 | 0.0680 | 3.4164 | 0.0006 | 0.0249 | Stromal |
| KIAA1324L | 33.3473 | -0.4499 | 0.1317 | 3.4154 | 0.0006 | 0.0249 | NA |
| AC008264.2 | 12.5942 | -0.3474 | 0.1017 | 3.4157 | 0.0006 | 0.0249 | Stromal |
| PPP1R14B | 88.0169 | 0.3390 | 0.0993 | -3.4138 | 0.0006 | 0.0250 | Epithelial |
| TNFRSF12A | 113.9624 | 0.4140 | 0.1213 | -3.4125 | 0.0006 | 0.0251 | NA |
| DCX | 16.1216 | -0.9160 | 0.2685 | 3.4114 | 0.0006 | 0.0251 | NA |
| CDC34 | 54.5584 | 0.2676 | 0.0785 | -3.4094 | 0.0007 | 0.0253 | Epithelial |

| | | | | | | | |
|---|---|---|---|---|---|---|---|
| SERBP1 | 329.0849 | 0.1647 | 0.0483 | -3.4061 | 0.0007 | 0.0254 | NA |
| OLA1 | 60.7751 | 0.3401 | 0.0998 | -3.4068 | 0.0007 | 0.0254 | Epithelial |
| CPLX2 | 1.2203 | -3.0295 | 0.8895 | 3.4059 | 0.0007 | 0.0254 | NA |
| TMEM98 | 49.7544 | -0.4023 | 0.1181 | 3.4059 | 0.0007 | 0.0254 | Stromal |
| B3GALT5 | 24.6858 | -0.8210 | 0.2411 | 3.4052 | 0.0007 | 0.0254 | Epithelial |
| RGS5 | 659.7148 | -0.5406 | 0.1590 | 3.4006 | 0.0007 | 0.0258 | NA |
| ADAMTS9-AS2 | 7.7106 | -0.6100 | 0.1794 | 3.4007 | 0.0007 | 0.0258 | Stromal |
| RPS13 | 176.9819 | 0.5086 | 0.1496 | -3.4000 | 0.0007 | 0.0258 | NA |
| RPS19BP1 | 105.0341 | 0.2244 | 0.0660 | -3.3987 | 0.0007 | 0.0258 | Epithelial |
| GK5 | 123.1691 | -0.2370 | 0.0698 | 3.3972 | 0.0007 | 0.0259 | Epithelial |
| BCAN | 1.1052 | -1.2684 | 0.3736 | 3.3955 | 0.0007 | 0.0260 | NA |
| NCOA1 | 170.0628 | -0.1788 | 0.0527 | 3.3946 | 0.0007 | 0.0261 | Stromal |
| ZNF25 | 37.6739 | -0.2588 | 0.0763 | 3.3926 | 0.0007 | 0.0262 | Stromal |
| HYOU1 | 129.8035 | 0.2874 | 0.0847 | -3.3917 | 0.0007 | 0.0262 | Epithelial |
| LAD1 | 63.1272 | 0.5913 | 0.1744 | -3.3903 | 0.0007 | 0.0263 | Epithelial |
| RPL7 | 314.2710 | 0.4946 | 0.1460 | -3.3880 | 0.0007 | 0.0265 | NA |
| NDUFA3 | 150.9675 | 0.2713 | 0.0801 | -3.3880 | 0.0007 | 0.0265 | NA |
| MTR | 138.5731 | -0.1717 | 0.0507 | 3.3862 | 0.0007 | 0.0265 | Stromal |
| WNT11 | 8.9677 | -0.8533 | 0.2520 | 3.3857 | 0.0007 | 0.0265 | NA |
| TTC5 | 31.3741 | -0.2266 | 0.0669 | 3.3866 | 0.0007 | 0.0265 | NA |
| RPL28 | 1375.5966 | 0.2237 | 0.0661 | -3.3861 | 0.0007 | 0.0265 | Stromal |
| DEAF1 | 51.8180 | -0.2287 | 0.0676 | 3.3822 | 0.0007 | 0.0268 | Epithelial |
| TMCO1 | 354.7888 | 0.2536 | 0.0751 | -3.3787 | 0.0007 | 0.0271 | Epithelial |
| UNC13A | 4.3236 | -1.0060 | 0.2982 | 3.3740 | 0.0007 | 0.0275 | NA |
| PKM | 329.3862 | 0.2443 | 0.0724 | -3.3726 | 0.0007 | 0.0276 | Epithelial |
| TGFBR3 | 139.3440 | -0.4476 | 0.1328 | 3.3705 | 0.0008 | 0.0276 | Stromal |
| CYS1 | 25.3732 | -0.5641 | 0.1674 | 3.3710 | 0.0007 | 0.0276 | Stromal |
| AL109628.1 | 13.7715 | 0.3942 | 0.1169 | -3.3714 | 0.0007 | 0.0276 | NA |
| PPIA | 107.7663 | 0.2176 | 0.0646 | -3.3682 | 0.0008 | 0.0278 | Epithelial |
| CHMP3 | 225.4888 | 0.1441 | 0.0429 | -3.3623 | 0.0008 | 0.0282 | NA |
| ADGRB3 | 2.5631 | -1.0137 | 0.3014 | 3.3633 | 0.0008 | 0.0282 | Stromal |
| ASPH | 456.5756 | 0.3880 | 0.1154 | -3.3623 | 0.0008 | 0.0282 | Epithelial |

| | | | | | | | |
|---|---|---|---|---|---|---|---|
| AC002558.3 | 32.1220 | -0.3564 | 0.1060 | 3.3612 | 0.0008 | 0.0283 | Stromal |
| ABHD1 | 4.7078 | -0.4945 | 0.1472 | 3.3601 | 0.0008 | 0.0284 | NA |
| ARHGAP6 | 23.7311 | -0.3846 | 0.1145 | 3.3593 | 0.0008 | 0.0284 | Stromal |
| AL157838.1 | 1.9917 | -0.8755 | 0.2609 | 3.3562 | 0.0008 | 0.0287 | NA |
| PRC1 | 9.8601 | 0.5247 | 0.1564 | -3.3548 | 0.0008 | 0.0287 | Epithelial |
| GPM6A | 4.1254 | -1.0004 | 0.2983 | 3.3533 | 0.0008 | 0.0288 | NA |
| LMBR1L | 70.7537 | -0.2175 | 0.0649 | 3.3535 | 0.0008 | 0.0288 | Stromal |
| FIRRE | 16.1603 | 0.7192 | 0.2145 | -3.3526 | 0.0008 | 0.0288 | NA |
| HINT1 | 221.4778 | 0.2537 | 0.0757 | -3.3494 | 0.0008 | 0.0291 | Epithelial |
| ADIPOQ | 27.9307 | -0.8060 | 0.2410 | 3.3440 | 0.0008 | 0.0296 | Stromal |
| HIF3A | 4.8096 | -0.7224 | 0.2160 | 3.3441 | 0.0008 | 0.0296 | Stromal |
| EVC | 12.7339 | -0.4408 | 0.1319 | 3.3431 | 0.0008 | 0.0296 | Stromal |
| KAT2A | 82.5945 | -0.2847 | 0.0852 | 3.3424 | 0.0008 | 0.0296 | NA |
| CSMD1 | 2.6103 | -1.2330 | 0.3694 | 3.3380 | 0.0008 | 0.0300 | NA |
| MUCL1 | 1794.3958 | 1.2490 | 0.3742 | -3.3379 | 0.0008 | 0.0300 | Epithelial |
| GPR137C | 9.0497 | -0.5797 | 0.1738 | 3.3360 | 0.0008 | 0.0302 | NA |
| CD37 | 71.5520 | 0.5880 | 0.1764 | -3.3338 | 0.0009 | 0.0303 | Stromal |
| DOLPP1 | 13.0467 | 0.3560 | 0.1069 | -3.3299 | 0.0009 | 0.0307 | NA |
| ANKLE2 | 161.7664 | -0.1923 | 0.0577 | 3.3299 | 0.0009 | 0.0307 | NA |
| AC018362.1 | 4.6743 | -0.4961 | 0.1490 | 3.3293 | 0.0009 | 0.0307 | NA |
| LINC01348 | 2.9414 | 0.9803 | 0.2947 | -3.3264 | 0.0009 | 0.0307 | NA |
| FAM228B | 25.1765 | -0.2922 | 0.0878 | 3.3276 | 0.0009 | 0.0307 | NA |
| NOP10 | 54.2809 | 0.3166 | 0.0952 | -3.3267 | 0.0009 | 0.0307 | NA |
| NCCRP1 | 14.8823 | 1.1291 | 0.3393 | -3.3280 | 0.0009 | 0.0307 | Epithelial |
| TSPO | 197.8738 | 0.2935 | 0.0882 | -3.3265 | 0.0009 | 0.0307 | Epithelial |
| SDC1 | 293.6702 | 0.5674 | 0.1707 | -3.3250 | 0.0009 | 0.0308 | Epithelial |
| HLA-V | 1.4827 | 1.2364 | 0.3721 | -3.3225 | 0.0009 | 0.0310 | NA |
| HOXB3 | 65.2271 | -0.6660 | 0.2005 | 3.3223 | 0.0009 | 0.0310 | NA |
| MYDGF | 118.0437 | 0.2883 | 0.0868 | -3.3220 | 0.0009 | 0.0310 | NA |
| RN7SL4P | 6.2685 | -0.5236 | 0.1576 | 3.3213 | 0.0009 | 0.0310 | NA |
| CFAP69 | 38.9754 | -0.4118 | 0.1240 | 3.3202 | 0.0009 | 0.0310 | NA |
| CAPN15 | 12.8688 | 0.4493 | 0.1353 | -3.3204 | 0.0009 | 0.0310 | NA |

| | | | | | | | |
|---|---|---|---|---|---|---|---|
| AHNAK | 931.1495 | -0.2234 | 0.0673 | 3.3191 | 0.0009 | 0.0310 | Stromal |
| RBX1 | 102.8468 | 0.2296 | 0.0692 | -3.3195 | 0.0009 | 0.0310 | NA |
| BMP2K | 126.9361 | -0.3588 | 0.1082 | 3.3173 | 0.0009 | 0.0312 | NA |
| SOCS7 | 81.7016 | 0.5269 | 0.1590 | -3.3145 | 0.0009 | 0.0314 | Epithelial |
| HES6 | 9.5789 | 0.6781 | 0.2047 | -3.3130 | 0.0009 | 0.0316 | Epithelial |
| TAC1 | 11.8456 | -0.9722 | 0.2939 | 3.3081 | 0.0009 | 0.0321 | Stromal |
| GSTO1 | 97.6788 | 0.2543 | 0.0769 | -3.3064 | 0.0009 | 0.0321 | Stromal |
| RTCB | 96.1354 | 0.2008 | 0.0607 | -3.3065 | 0.0009 | 0.0321 | Epithelial |
| PMF1 | 57.4479 | 0.2684 | 0.0812 | -3.3051 | 0.0009 | 0.0322 | Epithelial |
| DNAJB11 | 71.0043 | 0.2093 | 0.0633 | -3.3043 | 0.0010 | 0.0323 | NA |
| TNS2 | 103.3789 | -0.2463 | 0.0746 | 3.3040 | 0.0010 | 0.0323 | Stromal |
| EIF3M | 82.0818 | 0.1742 | 0.0527 | -3.3027 | 0.0010 | 0.0323 | NA |
| LIG3 | 84.2878 | -0.2949 | 0.0893 | 3.3028 | 0.0010 | 0.0323 | Epithelial |
| ATP5MC3 | 215.1757 | 0.2181 | 0.0661 | -3.3018 | 0.0010 | 0.0323 | Epithelial |
| PRDM6 | 12.0138 | -0.6173 | 0.1870 | 3.3008 | 0.0010 | 0.0323 | Stromal |
| IFI27 | 586.8236 | -0.6902 | 0.2091 | 3.3009 | 0.0010 | 0.0323 | NA |
| COLEC12 | 98.3482 | -0.4106 | 0.1244 | 3.3003 | 0.0010 | 0.0323 | Stromal |
| HLA-DRB1 | 567.0707 | 0.4592 | 0.1392 | -3.2985 | 0.0010 | 0.0325 | Stromal |
| C18orf21 | 17.2745 | 0.2827 | 0.0857 | -3.2982 | 0.0010 | 0.0325 | NA |
| RBMS3 | 160.5167 | -0.2926 | 0.0888 | 3.2962 | 0.0010 | 0.0327 | Stromal |
| ATP5MGL | 5.0676 | 0.4768 | 0.1447 | -3.2950 | 0.0010 | 0.0327 | Stromal |
| SF3B4 | 8.1001 | 0.5431 | 0.1650 | -3.2914 | 0.0010 | 0.0329 | NA |
| DYNC1I2 | 184.0261 | 0.2111 | 0.0642 | -3.2914 | 0.0010 | 0.0329 | NA |
| AC005550.2 | 4.2622 | -0.6457 | 0.1961 | 3.2932 | 0.0010 | 0.0329 | NA |
| GGCT | 131.9023 | 0.3776 | 0.1147 | -3.2920 | 0.0010 | 0.0329 | Epithelial |
| TSIX | 7.2899 | -0.4714 | 0.1432 | 3.2922 | 0.0010 | 0.0329 | NA |
| KRT16 | 7.0167 | 0.7919 | 0.2407 | -3.2897 | 0.0010 | 0.0330 | Epithelial |
| CHD6 | 203.2044 | -0.2284 | 0.0694 | 3.2900 | 0.0010 | 0.0330 | Epithelial |
| NR2F2 | 463.3689 | -0.2579 | 0.0784 | 3.2891 | 0.0010 | 0.0330 | Stromal |
| TET1 | 28.3828 | -0.3892 | 0.1183 | 3.2886 | 0.0010 | 0.0330 | NA |
| RIC8B | 38.7839 | -0.3097 | 0.0942 | 3.2875 | 0.0010 | 0.0331 | NA |
| NPAS3 | 10.1017 | -0.5627 | 0.1712 | 3.2862 | 0.0010 | 0.0331 | NA |

| | | | | | | | |
|---|---|---|---|---|---|---|---|
| CLTC | 559.7193 | 0.3426 | 0.1042 | -3.2862 | 0.0010 | 0.0331 | Epithelial |
| HSPA12B | 19.8831 | -0.4223 | 0.1285 | 3.2869 | 0.0010 | 0.0331 | Stromal |
| RPL27A | 1819.9020 | 0.2186 | 0.0666 | -3.2820 | 0.0010 | 0.0335 | Stromal |
| UTP11 | 59.9357 | 0.1808 | 0.0551 | -3.2811 | 0.0010 | 0.0335 | NA |
| PELO | 21.8338 | 0.2990 | 0.0912 | -3.2787 | 0.0010 | 0.0338 | Stromal |
| AL049838.1 | 3.8978 | -0.7092 | 0.2165 | 3.2760 | 0.0011 | 0.0340 | Stromal |
| RECK | 25.3559 | -0.3615 | 0.1104 | 3.2747 | 0.0011 | 0.0341 | Stromal |
| TTC17 | 185.9962 | -0.1623 | 0.0495 | 3.2749 | 0.0011 | 0.0341 | NA |
| CALM2 | 473.1091 | 0.1973 | 0.0603 | -3.2714 | 0.0011 | 0.0343 | Epithelial |
| AC092620.1 | 18.6876 | -0.4847 | 0.1482 | 3.2713 | 0.0011 | 0.0343 | Epithelial |
| LMCD1-AS1 | 3.1337 | -0.7214 | 0.2206 | 3.2704 | 0.0011 | 0.0343 | NA |
| ITGA9-AS1 | 9.9439 | -0.4504 | 0.1377 | 3.2716 | 0.0011 | 0.0343 | NA |
| IMPDH2 | 187.4097 | 0.2461 | 0.0752 | -3.2708 | 0.0011 | 0.0343 | Epithelial |
| YY1AP1 | 116.6549 | -0.2519 | 0.0770 | 3.2696 | 0.0011 | 0.0344 | Epithelial |
| NOP58 | 210.4541 | 0.1743 | 0.0533 | -3.2668 | 0.0011 | 0.0346 | Epithelial |
| ATIC | 89.9696 | 0.2073 | 0.0635 | -3.2666 | 0.0011 | 0.0346 | Epithelial |
| KLK12 | 1.3456 | -2.2679 | 0.6951 | 3.2630 | 0.0011 | 0.0350 | Epithelial |
| ADAMTS5 | 102.5498 | -0.3929 | 0.1204 | 3.2631 | 0.0011 | 0.0350 | Stromal |
| NPIPB2 | 7.1652 | -0.5369 | 0.1646 | 3.2618 | 0.0011 | 0.0351 | NA |
| KRAS | 108.7721 | 0.2231 | 0.0684 | -3.2602 | 0.0011 | 0.0352 | NA |
| AL512770.1 | 5.3722 | -0.4772 | 0.1464 | 3.2592 | 0.0011 | 0.0353 | NA |
| ATP1B1 | 439.1032 | 0.4404 | 0.1352 | -3.2571 | 0.0011 | 0.0355 | Epithelial |
| UCP2 | 160.2975 | 0.3625 | 0.1114 | -3.2552 | 0.0011 | 0.0356 | NA |
| CBFA2T2 | 86.6974 | -0.2128 | 0.0654 | 3.2549 | 0.0011 | 0.0356 | Epithelial |
| RCOR3 | 133.4266 | -0.2582 | 0.0794 | 3.2531 | 0.0011 | 0.0358 | Epithelial |
| PCGF3 | 65.8560 | -0.2127 | 0.0654 | 3.2524 | 0.0011 | 0.0358 | NA |
| PSMA5 | 94.9886 | 0.2005 | 0.0617 | -3.2506 | 0.0012 | 0.0360 | Epithelial |
| RBM5 | 248.0450 | -0.2007 | 0.0617 | 3.2500 | 0.0012 | 0.0360 | NA |
| KMT5B | 164.0481 | -0.1920 | 0.0591 | 3.2483 | 0.0012 | 0.0362 | Epithelial |
| RGMB-AS1 | 3.1198 | 0.6735 | 0.2075 | -3.2461 | 0.0012 | 0.0364 | NA |
| AC073869.1 | 14.0922 | -0.4394 | 0.1354 | 3.2452 | 0.0012 | 0.0364 | NA |
| ATP5F1A | 317.2114 | 0.2051 | 0.0632 | -3.2452 | 0.0012 | 0.0364 | Epithelial |

| | | | | | | | |
|---|---|---|---|---|---|---|---|
| EID1 | 521.0178 | -0.1572 | 0.0485 | 3.2447 | 0.0012 | 0.0364 | Stromal |
| PLAGL1 | 66.6017 | -0.4008 | 0.1236 | 3.2429 | 0.0012 | 0.0365 | Stromal |
| BSPRY | 66.1595 | 0.4097 | 0.1263 | -3.2430 | 0.0012 | 0.0365 | Epithelial |
| ADHFE1 | 12.3683 | -0.4249 | 0.1310 | 3.2422 | 0.0012 | 0.0366 | Epithelial |
| RNU4-2 | 81.0032 | -0.2605 | 0.0804 | 3.2410 | 0.0012 | 0.0367 | NA |
| SNORD1B | 7.3125 | -0.5634 | 0.1738 | 3.2406 | 0.0012 | 0.0367 | NA |
| ATP13A4 | 21.8956 | 0.9382 | 0.2897 | -3.2384 | 0.0012 | 0.0369 | Epithelial |
| TP53BP1 | 160.5233 | -0.1937 | 0.0598 | 3.2379 | 0.0012 | 0.0369 | NA |
| OBSCN | 35.1556 | -0.3454 | 0.1067 | 3.2366 | 0.0012 | 0.0370 | NA |
| SMIM4 | 62.9278 | 0.2970 | 0.0918 | -3.2360 | 0.0012 | 0.0370 | Epithelial |
| PLIN1 | 60.4529 | -0.6839 | 0.2115 | 3.2341 | 0.0012 | 0.0372 | Stromal |
| SMC1A | 192.3521 | 0.1864 | 0.0576 | -3.2335 | 0.0012 | 0.0372 | Epithelial |
| VEGFD | 7.7826 | -0.8311 | 0.2571 | 3.2328 | 0.0012 | 0.0373 | Stromal |
| NPY1R | 183.3997 | -1.1260 | 0.3484 | 3.2323 | 0.0012 | 0.0373 | Epithelial |
| C1orf43 | 143.8168 | 0.2330 | 0.0721 | -3.2309 | 0.0012 | 0.0374 | Epithelial |
| SNHG16 | 112.9516 | 0.3272 | 0.1013 | -3.2298 | 0.0012 | 0.0375 | Epithelial |
| SRSF10 | 158.2835 | -0.1531 | 0.0474 | 3.2290 | 0.0012 | 0.0375 | NA |
| RPL22L1 | 49.1866 | 0.2790 | 0.0865 | -3.2253 | 0.0013 | 0.0379 | NA |
| ZNF136 | 37.3025 | -0.2138 | 0.0663 | 3.2253 | 0.0013 | 0.0379 | NA |
| AL450998.3 | 1.6989 | -0.7960 | 0.2470 | 3.2233 | 0.0013 | 0.0381 | NA |
| D2HGDH | 79.1685 | -0.2750 | 0.0853 | 3.2227 | 0.0013 | 0.0381 | Epithelial |
| RAP1B | 129.4365 | -0.2020 | 0.0627 | 3.2223 | 0.0013 | 0.0381 | Stromal |
| MTRNR2L5 | 10.3197 | -0.4389 | 0.1362 | 3.2217 | 0.0013 | 0.0381 | Epithelial |
| WDFY3-AS2 | 12.5303 | -0.3764 | 0.1169 | 3.2193 | 0.0013 | 0.0384 | Stromal |
| SLIRP | 131.3500 | 0.2082 | 0.0647 | -3.2191 | 0.0013 | 0.0384 | Epithelial |
| SLC9A7 | 64.8600 | 0.3072 | 0.0955 | -3.2160 | 0.0013 | 0.0387 | Epithelial |
| BCHE | 2.3468 | -0.9673 | 0.3010 | 3.2142 | 0.0013 | 0.0389 | NA |
| NDUFA8 | 55.3085 | 0.2421 | 0.0753 | -3.2139 | 0.0013 | 0.0389 | Epithelial |
| LDB2 | 47.0764 | -0.3352 | 0.1044 | 3.2114 | 0.0013 | 0.0391 | Stromal |
| TUFM | 249.4489 | 0.1842 | 0.0574 | -3.2113 | 0.0013 | 0.0391 | Epithelial |
| UBE2D2 | 144.0976 | 0.1382 | 0.0431 | -3.2095 | 0.0013 | 0.0393 | NA |
| UBOX5 | 28.2587 | -0.2360 | 0.0735 | 3.2095 | 0.0013 | 0.0393 | Epithelial |

| | | | | | | | |
|---|---|---|---|---|---|---|---|
| CD160 | 3.1491 | -0.5881 | 0.1834 | 3.2066 | 0.0013 | 0.0395 | NA |
| RPS15 | 127.4942 | 0.4835 | 0.1508 | -3.2063 | 0.0013 | 0.0395 | Epithelial |
| NOP53 | 486.6634 | 0.2256 | 0.0703 | -3.2069 | 0.0013 | 0.0395 | Stromal |
| HMGB1P5 | 8.3697 | 0.9913 | 0.3092 | -3.2058 | 0.0013 | 0.0395 | NA |
| PIK3C2A | 226.0220 | -0.1734 | 0.0541 | 3.2040 | 0.0014 | 0.0397 | NA |
| CFAP300 | 3.8412 | -0.7328 | 0.2288 | 3.2030 | 0.0014 | 0.0398 | NA |
| EIF2S2 | 114.6314 | 0.2177 | 0.0680 | -3.2026 | 0.0014 | 0.0398 | Epithelial |
| DENND4C | 76.1426 | -0.1832 | 0.0572 | 3.2019 | 0.0014 | 0.0398 | NA |
| HOXA11 | 1.6328 | 1.4662 | 0.4584 | -3.1984 | 0.0014 | 0.0402 | NA |
| ANKIB1 | 171.3588 | -0.1614 | 0.0505 | 3.1984 | 0.0014 | 0.0402 | NA |
| MRPL48 | 22.7613 | 0.3204 | 0.1002 | -3.1973 | 0.0014 | 0.0403 | NA |
| AL035409.1 | 2.9864 | -1.6108 | 0.5040 | 3.1960 | 0.0014 | 0.0404 | Epithelial |
| B4GALT3 | 72.3199 | 0.3100 | 0.0970 | -3.1945 | 0.0014 | 0.0406 | Epithelial |
| ULK1 | 86.3955 | -0.2127 | 0.0666 | 3.1938 | 0.0014 | 0.0406 | Epithelial |
| STS | 44.1307 | 0.4639 | 0.1453 | -3.1931 | 0.0014 | 0.0406 | NA |
| CLDN18 | 1.5511 | -0.7831 | 0.2453 | 3.1923 | 0.0014 | 0.0407 | NA |
| NOMO1 | 19.4955 | 0.3942 | 0.1235 | -3.1911 | 0.0014 | 0.0408 | NA |
| RN7SL792P | 11.2814 | -0.4715 | 0.1478 | 3.1909 | 0.0014 | 0.0408 | NA |
| KIAA2026 | 226.8925 | -0.1480 | 0.0464 | 3.1895 | 0.0014 | 0.0409 | Stromal |
| ZDHHC12 | 27.8956 | 0.2840 | 0.0891 | -3.1882 | 0.0014 | 0.0410 | Epithelial |
| RPS4X | 2023.6509 | 0.2320 | 0.0728 | -3.1879 | 0.0014 | 0.0410 | NA |
| RPS14 | 616.3547 | 0.3213 | 0.1009 | -3.1844 | 0.0015 | 0.0415 | NA |
| CCNB1IP1 | 30.5480 | 0.2908 | 0.0914 | -3.1804 | 0.0015 | 0.0420 | NA |
| RPS28 | 232.8941 | 0.4454 | 0.1401 | -3.1801 | 0.0015 | 0.0420 | Stromal |
| FBXW8 | 30.4072 | -0.2541 | 0.0800 | 3.1785 | 0.0015 | 0.0421 | Epithelial |
| UBTF | 170.2226 | -0.1409 | 0.0443 | 3.1783 | 0.0015 | 0.0421 | Stromal |
| EMC3 | 102.7446 | 0.1914 | 0.0603 | -3.1762 | 0.0015 | 0.0423 | Epithelial |
| NF1 | 178.2077 | -0.2249 | 0.0708 | 3.1764 | 0.0015 | 0.0423 | NA |
| KLHL11 | 4.8918 | -0.5626 | 0.1772 | 3.1756 | 0.0015 | 0.0423 | NA |
| CALY | 2.3295 | 1.3735 | 0.4326 | -3.1748 | 0.0015 | 0.0424 | NA |
| DLGAP2 | 2.4377 | -0.8196 | 0.2584 | 3.1722 | 0.0015 | 0.0425 | NA |
| RNA5SP378 | 1.1003 | -1.2588 | 0.3967 | 3.1731 | 0.0015 | 0.0425 | NA |

| | | | | | | | |
|---|---|---|---|---|---|---|---|
| SUZ12P1 | 41.6942 | -0.3270 | 0.1031 | 3.1719 | 0.0015 | 0.0425 | NA |
| RNU1-98P | 6.2849 | -0.5464 | 0.1722 | 3.1726 | 0.0015 | 0.0425 | NA |
| MT-CO3 | 10158.3779 | 0.2931 | 0.0924 | -3.1732 | 0.0015 | 0.0425 | Epithelial |
| MT-ND5 | 1421.6634 | 0.2803 | 0.0884 | -3.1717 | 0.0015 | 0.0425 | Epithelial |
| CNDP2 | 160.0044 | 0.2343 | 0.0739 | -3.1712 | 0.0015 | 0.0425 | Epithelial |
| REV1 | 97.4209 | -0.1579 | 0.0498 | 3.1702 | 0.0015 | 0.0426 | NA |
| SOX5 | 16.2232 | -0.4595 | 0.1450 | 3.1697 | 0.0015 | 0.0426 | Stromal |
| AC068580.4 | 2.2304 | -0.8723 | 0.2752 | 3.1691 | 0.0015 | 0.0426 | NA |
| DNAJC3 | 180.2157 | 0.2301 | 0.0726 | -3.1689 | 0.0015 | 0.0426 | NA |
| B3GALT5-AS1 | 5.0078 | -0.9341 | 0.2949 | 3.1680 | 0.0015 | 0.0427 | NA |
| XRCC5 | 383.5691 | 0.1333 | 0.0421 | -3.1652 | 0.0015 | 0.0430 | Epithelial |
| C16orf54 | 9.4596 | 0.6357 | 0.2009 | -3.1643 | 0.0016 | 0.0430 | Stromal |
| RPS6KB1 | 95.6182 | 0.3690 | 0.1166 | -3.1645 | 0.0016 | 0.0430 | Epithelial |
| ZDHHC17 | 71.6643 | -0.2340 | 0.0740 | 3.1636 | 0.0016 | 0.0430 | Stromal |
| CSAD | 222.4910 | -0.3054 | 0.0966 | 3.1622 | 0.0016 | 0.0432 | Epithelial |
| AC011379.2 | 9.4042 | -0.5066 | 0.1603 | 3.1608 | 0.0016 | 0.0434 | NA |
| SEMA7A | 4.3260 | 0.6326 | 0.2002 | -3.1598 | 0.0016 | 0.0434 | Stromal |
| APBB2 | 118.3163 | -0.2985 | 0.0945 | 3.1580 | 0.0016 | 0.0436 | NA |
| WDR5 | 30.8305 | 0.2453 | 0.0778 | -3.1535 | 0.0016 | 0.0443 | NA |
| CHCHD4 | 12.6747 | 0.3391 | 0.1076 | -3.1525 | 0.0016 | 0.0443 | NA |
| MYH9 | 809.5347 | 0.1636 | 0.0519 | -3.1527 | 0.0016 | 0.0443 | Stromal |
| MKRN2 | 27.9893 | 0.2228 | 0.0707 | -3.1504 | 0.0016 | 0.0445 | NA |
| AL022342.1 | 1.7947 | 0.7074 | 0.2247 | -3.1488 | 0.0016 | 0.0447 | NA |
| AP003086.2 | 1.7086 | -0.8103 | 0.2576 | 3.1456 | 0.0017 | 0.0450 | NA |
| PHLDB1 | 145.5016 | -0.2265 | 0.0720 | 3.1463 | 0.0017 | 0.0450 | Stromal |
| LYRM9 | 28.5390 | -0.2987 | 0.0950 | 3.1456 | 0.0017 | 0.0450 | Stromal |
| CSKMT | 54.1899 | 0.4264 | 0.1356 | -3.1448 | 0.0017 | 0.0451 | Epithelial |
| RILPL1 | 38.3165 | -0.2167 | 0.0689 | 3.1444 | 0.0017 | 0.0451 | Stromal |
| TAL1 | 7.7780 | -0.3677 | 0.1171 | 3.1411 | 0.0017 | 0.0452 | Stromal |
| ANGEL2 | 51.1979 | -0.1912 | 0.0609 | 3.1405 | 0.0017 | 0.0452 | NA |
| RN7SKP55 | 15.9280 | -0.4900 | 0.1560 | 3.1413 | 0.0017 | 0.0452 | NA |
| EIPR1 | 17.4270 | 0.2596 | 0.0826 | -3.1413 | 0.0017 | 0.0452 | Epithelial |

| | | | | | | | |
|---|---|---|---|---|---|---|---|
| NBEAL1 | 126.9912 | -0.1620 | 0.0515 | 3.1423 | 0.0017 | 0.0452 | NA |
| MANF | 118.7692 | 0.2485 | 0.0791 | -3.1405 | 0.0017 | 0.0452 | NA |
| AC087239.1 | 2.2180 | 0.6081 | 0.1936 | -3.1408 | 0.0017 | 0.0452 | Epithelial |
| AL049780.1 | 2.2465 | -0.8073 | 0.2570 | 3.1416 | 0.0017 | 0.0452 | NA |
| RSL1D1 | 325.9444 | 0.1669 | 0.0532 | -3.1401 | 0.0017 | 0.0452 | Epithelial |
| ACTR1B | 51.7867 | 0.1888 | 0.0601 | -3.1395 | 0.0017 | 0.0452 | Epithelial |
| RPLP2 | 1424.5660 | 0.3192 | 0.1017 | -3.1391 | 0.0017 | 0.0452 | Stromal |
| TWF2 | 47.2722 | 0.2417 | 0.0771 | -3.1367 | 0.0017 | 0.0455 | Stromal |
| PCSK1 | 5.5923 | -1.3016 | 0.4150 | 3.1363 | 0.0017 | 0.0456 | NA |
| AC011815.2 | 3.2884 | -0.7209 | 0.2299 | 3.1353 | 0.0017 | 0.0456 | NA |
| RPL5 | 832.3017 | 0.2382 | 0.0760 | -3.1327 | 0.0017 | 0.0459 | Stromal |
| CCT4 | 109.1307 | 0.1827 | 0.0583 | -3.1328 | 0.0017 | 0.0459 | Epithelial |
| MT-TV | 169.4806 | 0.5299 | 0.1691 | -3.1334 | 0.0017 | 0.0459 | Epithelial |
| ARF1 | 327.8007 | 0.2142 | 0.0684 | -3.1319 | 0.0017 | 0.0459 | Epithelial |
| PIKFYVE | 73.5372 | -0.1661 | 0.0531 | 3.1298 | 0.0017 | 0.0462 | Stromal |
| CTBP1-DT | 29.7469 | -0.2551 | 0.0815 | 3.1291 | 0.0018 | 0.0462 | Epithelial |
| CLDN4 | 269.5882 | 0.3588 | 0.1147 | -3.1283 | 0.0018 | 0.0462 | Epithelial |
| HECTD4 | 150.1558 | -0.2069 | 0.0661 | 3.1283 | 0.0018 | 0.0462 | Epithelial |
| FBL | 216.3621 | 0.2055 | 0.0657 | -3.1282 | 0.0018 | 0.0462 | NA |
| PTOV1 | 109.4092 | 0.2562 | 0.0819 | -3.1272 | 0.0018 | 0.0463 | Epithelial |
| CTNNBIP1 | 60.7606 | 0.3139 | 0.1006 | -3.1206 | 0.0018 | 0.0469 | Epithelial |
| FBXO42 | 63.3500 | -0.1718 | 0.0550 | 3.1225 | 0.0018 | 0.0469 | NA |
| MAEL | 1.9251 | -0.9853 | 0.3157 | 3.1206 | 0.0018 | 0.0469 | NA |
| REEP5 | 300.7409 | 0.2332 | 0.0747 | -3.1207 | 0.0018 | 0.0469 | Epithelial |
| UBN2 | 178.2958 | -0.2065 | 0.0662 | 3.1215 | 0.0018 | 0.0469 | Epithelial |
| GPT2 | 30.9993 | 0.4760 | 0.1525 | -3.1207 | 0.0018 | 0.0469 | Epithelial |
| NAGS | 5.3729 | -0.8211 | 0.2630 | 3.1225 | 0.0018 | 0.0469 | NA |
| AL133520.1 | 1.3277 | 0.8235 | 0.2638 | -3.1215 | 0.0018 | 0.0469 | NA |
| CLHC1 | 30.3029 | -0.3171 | 0.1017 | 3.1183 | 0.0018 | 0.0470 | Epithelial |
| SEMA3D | 15.5348 | -0.4202 | 0.1348 | 3.1180 | 0.0018 | 0.0470 | Stromal |
| AC115837.1 | 1.1701 | -0.8140 | 0.2609 | 3.1194 | 0.0018 | 0.0470 | NA |
| PPFIA1 | 216.4723 | -0.3010 | 0.0966 | 3.1176 | 0.0018 | 0.0470 | Epithelial |

| | | | | | | | |
|---|---|---|---|---|---|---|---|
| ITCH | 136.2446 | -0.1919 | 0.0615 | 3.1191 | 0.0018 | 0.0470 | NA |
| AL022476.1 | 1.1179 | -0.9935 | 0.3187 | 3.1177 | 0.0018 | 0.0470 | NA |
| F11R | 134.6530 | 0.2807 | 0.0901 | -3.1160 | 0.0018 | 0.0470 | Epithelial |
| CIDEC | 15.4688 | -0.7467 | 0.2396 | 3.1162 | 0.0018 | 0.0470 | Stromal |
| AC010623.1 | 1.4559 | -0.9396 | 0.3015 | 3.1166 | 0.0018 | 0.0470 | NA |
| GPD1 | 33.9058 | -0.7142 | 0.2292 | 3.1159 | 0.0018 | 0.0470 | Stromal |
| COX5B | 163.4785 | 0.2012 | 0.0646 | -3.1151 | 0.0018 | 0.0470 | Epithelial |
| ICA1L | 25.1031 | -0.3428 | 0.1101 | 3.1150 | 0.0018 | 0.0470 | Stromal |
| BCR | 9.3406 | 0.4409 | 0.1416 | -3.1143 | 0.0018 | 0.0470 | NA |
| TSPAN9 | 55.7077 | 0.2617 | 0.0841 | -3.1125 | 0.0019 | 0.0473 | NA |
| SEC61A1 | 229.2580 | 0.2002 | 0.0644 | -3.1097 | 0.0019 | 0.0475 | NA |
| OMD | 21.1451 | -0.4149 | 0.1334 | 3.1099 | 0.0019 | 0.0475 | Stromal |
| TUBB4B | 149.4579 | 0.3123 | 0.1004 | -3.1101 | 0.0019 | 0.0475 | Epithelial |
| TPT1 | 1957.7779 | 0.2491 | 0.0801 | -3.1099 | 0.0019 | 0.0475 | Stromal |
| ATG2B | 47.6423 | -0.2091 | 0.0673 | 3.1082 | 0.0019 | 0.0477 | NA |
| MTND3P17 | 1.1783 | -0.7924 | 0.2551 | 3.1067 | 0.0019 | 0.0478 | NA |
| BID | 29.4510 | 0.2850 | 0.0917 | -3.1066 | 0.0019 | 0.0478 | Stromal |
| POLRMT | 25.0831 | 0.2492 | 0.0802 | -3.1061 | 0.0019 | 0.0478 | NA |
| TRPM6 | 4.3285 | -0.6579 | 0.2119 | 3.1049 | 0.0019 | 0.0478 | Stromal |
| STN1 | 29.7006 | 0.2412 | 0.0777 | -3.1056 | 0.0019 | 0.0478 | NA |
| SHMT2 | 65.3636 | 0.3366 | 0.1084 | -3.1051 | 0.0019 | 0.0478 | NA |
| STOX2 | 18.9472 | -0.4132 | 0.1331 | 3.1032 | 0.0019 | 0.0479 | Stromal |
| SOWAHA | 11.1448 | -0.7872 | 0.2537 | 3.1028 | 0.0019 | 0.0479 | Epithelial |
| MALAT1 | 48925.2279 | -0.4062 | 0.1309 | 3.1029 | 0.0019 | 0.0479 | Epithelial |
| ARPP19 | 119.1772 | 0.1786 | 0.0575 | -3.1038 | 0.0019 | 0.0479 | Epithelial |
| KANTR | 32.9643 | -0.2841 | 0.0915 | 3.1041 | 0.0019 | 0.0479 | NA |
| PDZK1IP1 | 56.4222 | 0.7729 | 0.2492 | -3.1020 | 0.0019 | 0.0479 | Epithelial |
| AC011503.1 | 2.0343 | -1.0850 | 0.3500 | 3.1005 | 0.0019 | 0.0481 | Stromal |
| SNHG19 | 84.6142 | 0.3782 | 0.1221 | -3.0982 | 0.0019 | 0.0484 | Epithelial |
| TBK1 | 53.9059 | -0.1917 | 0.0619 | 3.0975 | 0.0020 | 0.0485 | NA |
| PNPLA7 | 35.9658 | -0.3046 | 0.0984 | 3.0960 | 0.0020 | 0.0485 | NA |
| SRSF8 | 65.6018 | 0.2047 | 0.0661 | -3.0961 | 0.0020 | 0.0485 | NA |

| | | | | | | | |
|---|---|---|---|---|---|---|---|
| AP1B1 | 117.1570 | 0.2154 | 0.0696 | -3.0963 | 0.0020 | 0.0485 | NA |
| SUN2 | 88.1320 | 0.2355 | 0.0761 | -3.0943 | 0.0020 | 0.0488 | Stromal |
| DIP2C | 73.3586 | -0.1890 | 0.0611 | 3.0934 | 0.0020 | 0.0489 | Stromal |
| TBCA | 168.7333 | 0.2115 | 0.0684 | -3.0924 | 0.0020 | 0.0489 | Epithelial |
| FAM193B | 92.6232 | -0.2106 | 0.0681 | 3.0927 | 0.0020 | 0.0489 | NA |
| CFL1 | 1550.3793 | 0.1616 | 0.0523 | -3.0900 | 0.0020 | 0.0492 | Epithelial |
| LSM4 | 152.3750 | 0.2299 | 0.0744 | -3.0900 | 0.0020 | 0.0492 | Epithelial |
| SRM | 134.5017 | 0.2260 | 0.0732 | -3.0890 | 0.0020 | 0.0492 | NA |
| AC044787.1 | 2.7654 | 0.6419 | 0.2078 | -3.0894 | 0.0020 | 0.0492 | NA |
| FOXRED2 | 46.8172 | 0.3135 | 0.1015 | -3.0887 | 0.0020 | 0.0492 | NA |
| ARMCX1 | 34.1835 | -0.3304 | 0.1070 | 3.0879 | 0.0020 | 0.0493 | NA |
| PSMD14 | 94.8451 | 0.1765 | 0.0572 | -3.0833 | 0.0020 | 0.0498 | Epithelial |
| ZZEF1 | 76.8722 | -0.2104 | 0.0682 | 3.0836 | 0.0020 | 0.0498 | Stromal |
| AC079336.5 | 2.7864 | -0.6102 | 0.1979 | 3.0834 | 0.0020 | 0.0498 | Stromal |
| AP2M1 | 235.2423 | 0.1484 | 0.0481 | -3.0825 | 0.0021 | 0.0499 | NA |
| PPA1 | 71.3369 | 0.2360 | 0.0766 | -3.0818 | 0.0021 | 0.0500 | NA |

**Table S4:** Associations between recurrent CNAs and sequencing coverage or cohort. Supplemental to Figure 4.

| | **Association with coverage** | | | **Association with cohort** | | |
|---|---|---|---|---|---|---|
| **Recurrent CNAs** | **Difference in medians** | **P-values** | **FDR** | **Difference in medians** | **P-value** | **FDR** |
| Amp_1q42.3 | 1550611 | 0.321 | 0.416 | 0.033 | 0.757 | 0.955 |
| Del_17p13.2 | 2800768 | 0.931 | 0.931 | 0.058 | 0.444 | 0.804 |
| Amp_17q12 | 2800768 | 0.651 | 0.697 | 0.044 | 0.587 | 0.895 |
| Amp_17q23.2 | 1737324 | 0.758 | 0.784 | 0.044 | 0.954 | 0.996 |
| Del_16q24.1 | 1217342 | 0.349 | 0.431 | 0.222 | 0.102 | 0.591 |
| Amp_8q24.21 | 2532147 | 0.882 | 0.897 | -0.071 | 0.348 | 0.745 |
| Del_11q25 | -6290327 | 0.618 | 0.674 | -0.196 | 0.644 | 0.903 |
| Amp_8q13.1 | -8252690 | 0.139 | 0.206 | 0 | 0.654 | 0.903 |
| Amp_20q13.2 | 7046467 | 0.132 | 0.201 | 0 | 0.436 | 0.804 |
| Del_11q22.3 | 13158102 | 0.032 | 0.058 | 0 | 0.314 | 0.745 |
| Amp_17q21.31 | -2459037 | 0.180 | 0.259 | 0 | 0.996 | 0.996 |
| Amp_8p11.23 | 20778292 | 0.036 | 0.062 | 0 | 0.957 | 0.996 |
| Del_3p14.1 | 2684015 | 0.729 | 0.767 | 0 | 0.163 | 0.599 |
| Del_8p11.22 | 1648757 | 0.518 | 0.585 | 0 | 0.343 | 0.745 |
| Amp_11q13.3 | 13932379 | 0.423 | 0.486 | 0 | 0.757 | 0.955 |
| Del_17q12 | -8452507 | 0.353 | 0.431 | 0 | 0.575 | 0.895 |
| Del_17q21.2 | 10296570 | 0.207 | 0.281 | 0 | 0.993 | 0.996 |
| Amp_19q13.12 | 10494689 | 0.341 | 0.431 | 0 | 0.338 | 0.745 |
| Amp_12q15 | -12560166 | 0.050 | 0.081 | 0 | 0.029 | 0.233 |
| Del_11p15.4 | 3708923 | 0.304 | 0.403 | 0 | 0.275 | 0.745 |
| Del_5q31.1 | 8548178 | 0.388 | 0.455 | 0 | 0.827 | 0.996 |
| Del_11q11 | -166682 | 0.610 | 0.674 | 0 | 0.531 | 0.895 |
| Del_1p21.3 | 18621923 | 0.044 | 0.073 | 0 | 0.002 | **0.028** |
| Del_11q13.1 | 10494689 | 0.061 | 0.095 | 0 | 0.165 | 0.599 |
| Amp_16q12.2 | -7615355 | 0.367 | 0.439 | 0 | 0.032 | 0.233 |
| Del_4q35.2 | -11531034 | 0.200 | 0.277 | 0 | 0.157 | 0.599 |
| Amp_6q25.1 | -7615355 | 0.182 | 0.259 | 0 | 0.360 | 0.745 |
| Del_10p15.3 | -21959310 | 0.027 | 0.050 | 0 | 0.000 | **0.005** |

| | | | | | | |
|---|---|---|---|---|---|---|
| Del_19p13.3 | -30105171 | 0.042 | 0.072 | 0 | 0.904 | 0.996 |

**Table S5:** Univariate Cox regression analysis of CSx cell type abundance towards progression (any iBE) in RAHBT LCM. Supplemental to Figure 5.

| | **coef** | **exp(coef)** | **se(coef)** | **z** | **P-value** | **FDR** |
|---|---|---|---|---|---|---|
| **mDC** | 25.36863 | 1.041E+11 | 6.383095 | 3.974347 | 7.06E-05 | **0.00101071** |
| **CD4** | 10.88261 | 53242.31 | 2.838826 | 3.833489 | 0.00012634 | **0.00101071** |
| **pDC** | 28.35189 | 2.06E+12 | 8.856115 | 3.201391 | 0.00136766 | **0.00554251** |
| **Immune** | 2.605316 | 13.5355 | 0.8147652 | 3.197628 | 0.00138563 | **0.00554251** |
| **NKT** | 19.62875 | 334702207 | 8.055231 | 2.436771 | 0.01481907 | **0.04595117** |
| **Fibroblast** | -2.624797 | 0.07245444 | 1.102057 | -2.381727 | 0.01723169 | **0.04595117** |
| Macrophage | 15.06461 | 3487218 | 6.808633 | 2.212576 | 0.02692691 | 0.06154722 |
| Mast | 19.44246 | 277813316 | 9.526299 | 2.040925 | 0.0412583 | 0.0825166 |
| Monocyte | 11.71033 | 121823.5 | 7.007859 | 1.671028 | 0.09471613 | 0.16838423 |
| Endothelial | -3.857939 | 0.02111147 | 2.48589 | -1.551935 | 0.1206778 | 0.19308448 |
| CD8 | 5.864071 | 352.1549 | 4.018355 | 1.459321 | 0.1444767 | 0.21014793 |
| Neutrophil | 20.47718 | 781856966 | 14.86406 | 1.377631 | 0.1683173 | 0.22442307 |
| TReg | 11.2979 | 80651.73 | 10.19587 | 1.108086 | 0.2678248 | 0.32963052 |
| B_Cells | 7.016958 | 1115.388 | 7.655518 | 0.9165882 | 0.3593585 | 0.41069543 |
| NK | 7.482623 | 1776.895 | 18.85228 | 0.396908 | 0.6914353 | 0.73753099 |
| Epithelial | -0.1520237 | 0.8589679 | 0.6357247 | -0.2391345 | 0.8110013 | 0.8110013 |
